# Multimodal mapping of systemic inflammation and immunity across the spectrum of CFTR dysfunction

**DOI:** 10.64898/2026.03.25.714282

**Authors:** L. Jonckheere, S. J. Tavernier, I. Janssens, Y. Vande Weygaerde, H. Schaballie, P. Schelstraete, S. Van Biervliet, R. Browaeys, N. Vandamme, E. Duthoo, S. Riemann, T. Maes, V. Bosteels, F. Haerynck, B. N. Lambrecht, C. Bosteels, E. Van Braeckel

**Author notes:** **Corresponding author:** Eva Van Braeckel, MD, PhD, Respiratory Infection and Defense Lab (RIDL) Lung Research Centre, MRB-II, Corneel Heymanslaan 10, B-9000 Ghent, Belgium.

## Abstract

Cystic fibrosis is traditionally framed as a dichotomy between affected individuals and clinically unaffected carriers, yet the systemic immune consequences across the spectrum of CFTR functionality remain incompletely defined. The advent of highly effective CFTR modulators now provides a unique momentum to examine whether partial restoration of CFTR function can influence systemic immunity. Using multimodal immune profiling, we constructed a single-cell atlas of circulating immune cells in people with cystic fibrosis (pwCF), healthy *F508del* carriers and non-carriers. In pwCF, systemic immunity was markedly altered following in vivo CFTR modulation with elexacaftor-tezacaftor-ivacaftor, with broad reductions in pro-inflammatory cytokines linked to improved clinical outcomes. Notably, healthy *F508del* carriers exhibited a CF-like immune signature characterised by low-grade systemic inflammation, including elevated IL-6, reduced mucosal-associated invariant T cells, and inflammatory monocyte features overlapping with pwCF. Together, these findings show that CFTR-related immune dysregulation extends beyond classical cystic fibrosis, challenging a strict dichotomy between health and disease.

## INTRODUCTION

Cystic fibrosis (CF) is a life-limiting autosomal recessive disorder caused by mutations in the *CFTR* gene (1). Loss of functional CFTR disrupts chloride and bicarbonate transport and dysregulates epithelial sodium absorption, leading to airway surface liquid dehydration, impaired mucociliary clearance, and chronic infection-driven neutrophilic inflammation. This cycle ultimately causes progressive structural lung disease, the major determinant of morbidity and mortality (2).

Beyond epithelial dysfunction throughout the body, loss of CFTR functionality has the potential to influence systemic immunity. Chronic microbial exposure sustains inflammatory activation of circulating immune cells, and accumulating evidence indicates that CFTR has intrinsic roles in immune regulation (3). Enhanced epithelial sodium influx and reduced bicarbonate transport promote NLRP3 inflammasome activation, increasing IL-1β and IL-6 (4) and priming immune cells for exaggerated responses. Importantly, CFTR is also expressed in circulating monocytes and macrophages, where its dysfunction has been linked to impaired phagocytosis, exaggerated inflammatory cytokine production, and dysregulated inflammasome signalling, indicating cell-intrinsic effects beyond the epithelial compartment (5–7). Neutrophils from people with CF (pwCF) display impaired bacterial clearance and excessive mediator release (8), while macrophages exhibit reduced phagocytic capacity and heightened inflammasome activity (9). Adaptive immunity is also affected: CFTR-deficient T cells show abnormal calcium signalling and cytokine secretion (10), naïve CD4⁺ T cells skew toward T-helper (Th)17 differentiation in the presence of functionally impaired CFTR (11), and CFTR-deficient B cells demonstrate increased activation and IL-6 production (12). Together, these innate and adaptive alterations create a state of chronic systemic inflammation akin to autoinflammatory diseases (13).

Highly effective CFTR modulators, particularly elexacaftor-tezacaftor-ivacaftor (ETI), have transformed CF care, improving lung function, nutritional status, and quality of life (14,15). ETI reduces airway neutrophilia, neutrophil elastase activity, and local inflammatory cytokine production (16–18), yet its impact on circulating immune cells remains incompletely characterized. Small ex-vivo studies suggest reduced NLRP3 activation and lower expression of pro-inflammatory cytokine genes in pwCF on ETI (19), but treatment durations were short and the extent to which these changes reflect direct effects on immune cells or secondary consequences of improved airway health is unclear.

Historically, pathological consequences were attributed only to individuals carrying biallelic pathogenic *CFTR* variants. However, large epidemiological studies now show that heterozygous carriers have increased risk of bronchiectasis, pancreatitis, diabetes, respiratory infections and malignancy (20–23). Importantly, one in 25 Caucasians carries a *CFTR* mutation, primarily *F508del*, making this enhanced morbidity risk a potential concern on a population level (21,24). Transcriptomic comparisons between CF probands, their parents, and unrelated controls further reveal that carriers share multiple immune-related gene expression shifts with pwCF, suggesting that even monoallelic CFTR dysfunction may influence immune homeostasis (25).

In this study, we use the momentum of introduction of CFTR modulation in routine CF care, to comprehensively map systemic immune dynamics across the spectrum of CFTR dysfunction without a priori assumptions. Using cellular indexing of transcriptomes and epitopes by sequencing (CITE-seq) and flow cytometry of peripheral blood mononuclear cells (PBMCs), alongside serum cytokine profiling, we characterise changes in the circulating immune compartment associated with in vivo restoration of CFTR function in pwCF. Furthermore, by incorporating healthy controls - including *F508del* carriers and non-carriers - we delineate the broader effects of CFTR dysfunction and CFTR modulation on systemic immunity.

## RESULTS

### Clinical characteristics of pwCF and healthy controls

We included 90 pwCF with paired serum samples obtained before and after at least six months of ETI therapy. Of these, 29 had previously been treated with tezacaftor-ivacaftor (TI) before transitioning to ETI (TI-to-ETI group), whereas 61 initiated ETI as their first CFTR modulator therapy (primary-ETI group). Median treatment duration at the time of sampling was 7.6 months (IQR 6.3 - 8.3) on TI and 7.4 months (IQR 6.7 - 8.3) on ETI. The TI-to-ETI group consisted of adult pwCF, the majority of whom were homozygous for *F508del* (79.3%), while the primary-ETI group included both children (aged < 18 years old) and adults homozygous for *F508del* and compound *F508del* heterozygotes. Clinical characteristics of both groups, baseline and after initiation of modulator therapy, are presented in **Table 1**. Consistent with the established clinical effects of CFTR modulation, the TI-to-ETI group showed a stepwise reduction in exacerbation frequency from baseline to TI and ETI, whereas improvements in lung function and BMI were primarily observed following ETI; overall, ETI therapy was associated with significant improvements in lung function and BMI alongside reductions in exacerbation frequency across both treatment groups.

**Table 1.** Clinical characteristics of study participants. People with cystic fibrosis (pwCF) who transitioned from tezacaftor-ivacaftor (TI) to elexacaftor-tezacaftor-ivacaftor (ETI) (TI-to-ETI cohort) are shown at baseline, after ≥6 months on TI, and after ≥6 months on ETI. For pwCF, within-subject comparisons are shown for baseline vs TI and TI vs ETI. PwCF initiating ETI as first CFTR modulator therapy (primary-ETI cohort) were stratified into children and adults and presented at baseline and after ≥6 months on ETI, with paired comparisons performed separately within each age group. Continuous variables are presented as mean ± standard deviation and categorical variables as number (percentage). Healthy controls (HC), including heterozygous *F508del* carriers and non-carriers were sampled once and are shown for descriptive reference only; no statistical comparisons with pwCF were performed. For the TI-to-ETI cohort, within-subject comparisons were performed using paired t-tests or Wilcoxon signed-rank tests as appropriate, with Bonferroni correction applied across the tested pairwise contrasts (baseline vs TI and TI vs ETI). For two-timepoint comparisons in the primary-ETI cohort, paired t-tests or Wilcoxon signed-rank tests were applied as appropriate. Categorical variables for paired pwCF comparisons were analysed using McNemar’s test. “ns” indicates not statistically significant. *, p < 0.05; **, p < 0.01; ***, p < 0.001; ****, p < 0.0001. Abbreviations: TI, tezacaftor-ivacaftor; ETI, elexacaftor-tezacaftor-ivacaftor; pwCF, people with cystic fibrosis; ppFEV₁, percent predicted forced expiratory volume in one second (post-bronchodilator); BMI, body mass index; PA, *Pseudomonas aeruginosa*; CFRD, cystic fibrosis-related diabetes; CFLD, cystic fibrosis-related liver disease; PI, pancreatic insufficiency; RF, residual function mutation; MF, minimal function mutation; O, other mutation class; WT, wild type.

|  | TI-to-ETI |  |  | Primary-ETI |  |  |  | Healthy controls |  |
| --- | --- | --- | --- | --- | --- | --- | --- | --- | --- |
|  | Adults with CF |  |  | Children with CF |  | Adults with CF |  | Carriers | Non-carriers |
|  | Baseline | TI | ETI | Baseline | ETI | Baseline | ETI |  |  |
| Total number, n | 29 | 29 | 29 | 34 | 34 | 27 | 27 | 23 | 9 |
| Age, years | 29.5±11.5 | 30.2±10.4 | 31.1±9.6 | 10.7±4.1 | 11.4±3.9 | 28.3±10 | 28.9±10.3 | 32.7±2.3 | 32.3±1.3 |
| Female sex, n | 16 (55.2%) | 16 (55.2%) | 16 (55.2%) | 20 (58.8%) | 20 (58.8%) | 14 (51.9%) | 14 (51.9%) | 15(65.2%) | 5 (55.5%) |
| Genotype |  |  |  |  |  |  |  |  |  |
| *F508del/F508del* | 23 (79.3%) | 23 (79.3%) | 23 (79.3%) | 14 (41.2%) | 14 (41.2%) | 2 (7.4%) | 2 (7.4%) | - | - |
| *F508del/RF* | 6 (20.7%) | 6 (20.7%) | 6 (20.7%) | 9 (26.5%) | 9 (26.5%) | 7 (25.9%) | 7 (25.9%) | - | - |
| *F508del/MF* | - | - | - | 11 (32.4%) | 11 (32.4%) | 17 (63%) | 17 (63%) | - | - |
| *RF/O* | - | - | - | - | - | 1 (3.7%) | 1 (3.7%) | - | - |
| *F508del/WT* | - | - | - | - | - | - | - | 23 (100%) | - |
| *WT/WT* | - | - | - | - | - | - | - | - | 9 (100%) |
| Post-BD ppFEV1, % | 71.2±20.2 | 74.4±18.8<sup>ns</sup> | 82.9±18.5<sup>***</sup> | 90.6±17.9 | 98.5±12.6<sup>***</sup> | 78.1±19.8 | 86.8±18.4<sup>***</sup> | - | - |
| BMI, kg/m<sup>2</sup> | 22.8±3.4 | 22.9±3.9<sup>ns</sup> | 23.6±3.3<sup>*</sup> | 16.8±3.4 | 17.6±3.5<sup>****</sup> | 21.5±3.3 | 22.5±3.2<sup>***</sup> | - | - |
| Chronic PA, n | 14 (48.3%) | 13 (44.8%)<sup>ns</sup> | 11 (37.9%)<sup>ns</sup> | 6 (17.6%) | 2 (6.1%)<sup>ns</sup> | 9 (33.3%) | 7 (25.9%)<sup>ns</sup> | - | - |
| CFRD, n | 12 (41.4%) | 12 (41.4%) | 12 (41.4%) | 2 (5.9%) | 2 (5.9%) | 7 (25.9%) | 7 (25.9%) | - | - |
| CFLD, n | 4 (13.8%) | 4 (13.8%) | 4 (13.8%) | 8 (23.5%) | 8 (23.5%) | 6 (22.2%) | 6 (22.2%) | - | - |
| PI, n | 23 (79.3%) | 23 (79.3%) | 23 (79.3%) | 31 (91.2%) | 31 (91.2%) | 20 (74.1%) | 20 (74.1%) | - | - |
| Exacerbations/yr, n | 1.66±1.78 | 0.83±1<sup>*</sup> | 0.31±0.54<sup>*</sup> | 1.76±1.65 | 0.79±0.91<sup>**</sup> | 1.93±1.8 | 0.33±1<sup>****</sup> | - | - |

For the CITE-seq and multiparameter flow cytometry experiments, PBMCs were selected from a subset of eight adult pwCF, selected for maximum homogeneity: homozygous for *F508del*, chronically colonised with *Pseudomonas aeruginosa*, and with established exocrine pancreatic insufficiency (mean age 25 years; four males, four females). Six participants belonged to the TI-to-ETI group and two to the primary-ETI group, with treatment durations of 7.6 months (IQR 6.4 - 8.0) on TI and 7.0 months (IQR 6.4 - 7.4) on ETI. Healthy controls for these experiments included four *F508del* carrier controls and four non-carrier controls (mean age 27 years for each group; two males and two females per group), confirmed by genetic screening for the 50 most frequent *CFTR* mutations, and frequency-matched for age and sex. These control groups will be further designated as *F508del* carriers and non-carriers, respectively. A detailed overview of cohort flow and sample availability for serum, CITE-seq and flow-cytometry analyses is provided in **Supplementary Figure 1**.

### *CFTR* heterozygosity is associated with a CF-like systemic immune phenotype

#### Single-cell PBMC atlas reveals reduced frequency of circulating mucosal-associated invariant T cells in both pwCF and *F508del* carriers

We constructed a single-cell atlas of PBMCs across distinct levels of CFTR functionality, integrating longitudinal samples from eight pwCF homozygous for F508del and healthy controls, including four heterozygous *F508del* carriers and four non-carriers **(Figure 1A)**. For pwCF, samples were obtained at baseline (pre-treatment), after at least six months on TI and subsequently after at least six months on ETI. Using CITE-seq, we profiled 213,479 PBMCs across all conditions, of which 193,985 cells were retained after quality-control filtering for downstream analysis. Quality-control metrics for the integrated CITE-seq dataset are shown in **Supplementary Figure 2**. Immune populations were identified using weighted-nearest neighbour clustering based on integrated RNA and antibody-derived tag (ADT) expression, followed by marker-based annotation using Azimuth reference-based mapping **(Figure 1B)**. Extended reference-mapping diagnostics and canonical RNA/ADT marker expression supporting cell-type annotation are shown in **Supplementary Figure 3**. Mucosal-associated invariant T (MAIT) cells showed consistently lower proportions in both pwCF (across all timepoints) and *F508del* carrier controls compared with non-carrier controls **(Figure 1C)**. In contrast, the overall distribution of major immune cell populations was largely preserved across groups and conditions. Quantification at the sample level using a linear mixed-effects model demonstrated marked reductions in MAIT-cell abundance across the spectrum of CFTR dysfunction. MAIT cells comprised 6% of PBMCs in healthy non-carriers, compared with 3% in both *F508del* carriers and pwCF. Model-derived estimated marginal mean differences corresponded to absolute reductions of 2.9 percentage points in *F508del* carriers (adjusted P = 0.026) and 3.9-4.0 percentage points in pwCF at baseline (adjusted P = 0.003), on TI (adjusted P = 0.003), and on ETI (adjusted P = 0.003) **(Figure 1D)**. No evidence of recovery in MAIT-cell abundance was observed in pwCF on CFTR modulation. Extended sample-level abundance analyses across all annotated PBMC populations are provided in **Supplementary Figure 4**.

**Figure 1.**
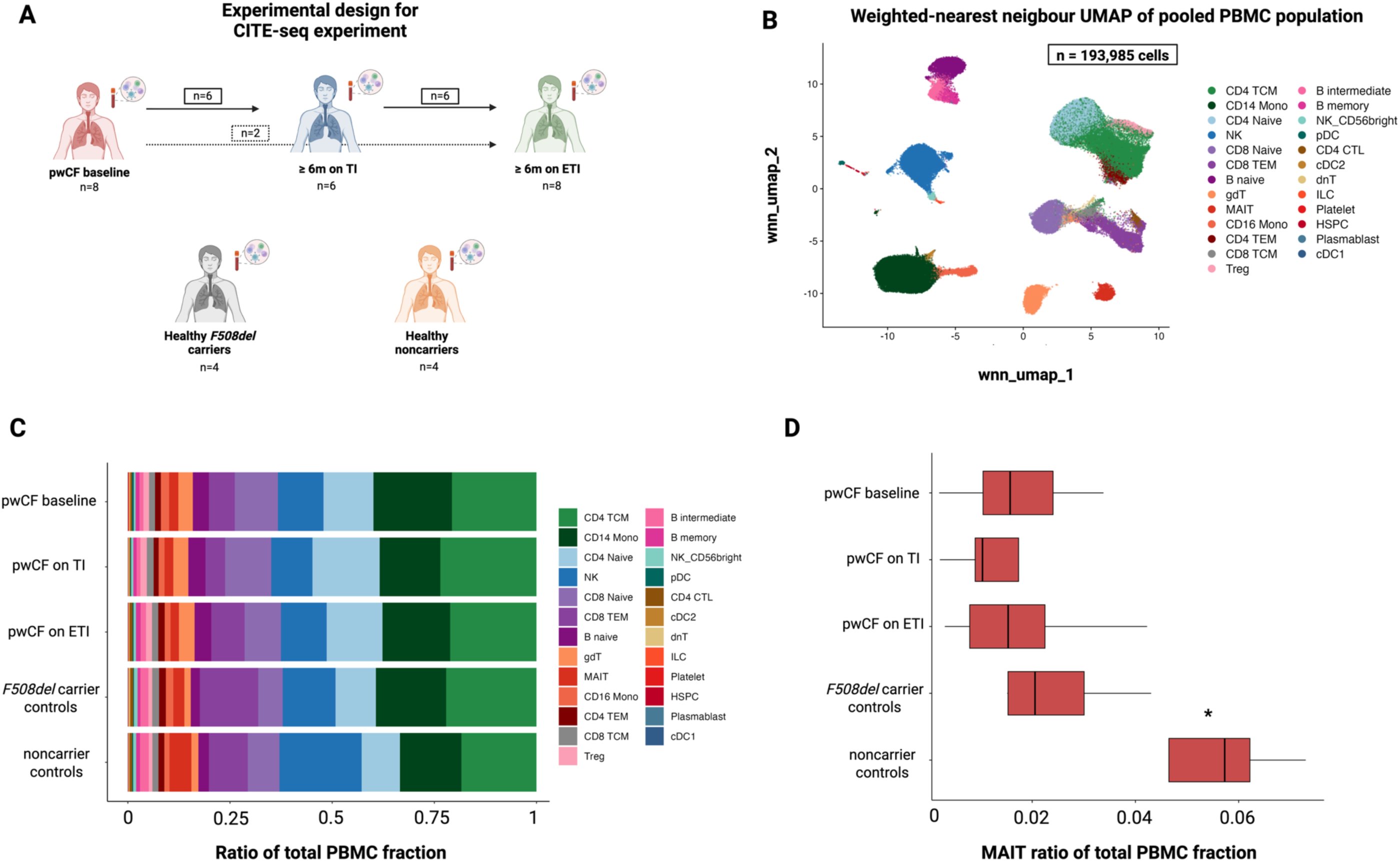
Single-cell CITE-seq profiling of peripheral blood immune cells across the spectrum of CFTR dysfunction. **(A) Overview of the experimental design**. Peripheral blood mononuclear cells (PBMCs) were obtained from people with cystic fibrosis (pwCF) at three disease states: prior to initiation of CFTR modulator therapy (baseline; *n* = 8), after ≥6 months of treatment with tezacaftor/ivacaftor (TI; *n* = 6), and after ≥6 months of treatment with elexacaftor/tezacaftor/ivacaftor (ETI; *n* = 8). PBMCs were also collected from healthy *F508del* heterozygous carriers (*n* = 4) and healthy non-carrier controls (*n* = 4). Cells were profiled using cellular indexing of transcriptomes and epitopes by sequencing (CITE-seq), enabling simultaneous measurement of transcriptomic and surface protein expression. **(B) Weighted-nearest neighbour (WNN) UMAP embedding of the integrated PBMC dataset coloured by annotated immune cell populations**. Cell types were defined using combined RNA and antibody-derived tag (ADT) expression following unsupervised clustering and marker-based annotation. **(C) Relative abundance of immune cell populations across study groups**. Bars represent the proportion of each annotated cell type within the pooled PBMC population for each condition, calculated as the fraction of cells assigned to each cell type relative to the total number of PBMCs in that condition. Colours correspond to the cell type annotations shown in panel B. **(D) Distribution of mucosal-associated invariant T (MAIT) cell proportions across study groups**. Each boxplot represents the per-sample ratio of MAIT cells relative to the total PBMC population. MAIT cell proportions were analysed using a linear mixed-effects model with condition included as a fixed effect and patient identity as a random intercept. Pairwise contrasts were estimated using estimated marginal means, with Benjamini-Hochberg correction for multiple testing. Boxes represent the interquartile range (IQR), the central line indicates the median, and whiskers extend to 1.5 × IQR. Significance is shown for comparisons between healthy non-carriers and each other group; asterisks indicate adjusted P values (\**P* < 0.05). Cell type abbreviations: CD4 TCM, CD4 central memory T cells; CD4 naïve, naïve CD4 T cells; CD4 TEM, CD4 effector memory T cells; CD8 TCM, CD8 central memory T cells; CD8 TEM, CD8 effector memory T cells; CD8 naïve, naïve CD8 T cells; NK, natural killer cells; NK_CD56bright, CD56^bright^ natural killer cells; CD14 Mono, classical CD14⁺ monocytes; CD16 Mono, CD16⁺ monocytes; B naïve, naïve B cells; B memory, memory B cells; B intermediate, intermediate B-cell population; gdT, γδ T cells; MAIT, mucosal-associated invariant T cells; pDC, plasmacytoid dendritic cells; cDC1, conventional dendritic cells type 1; cDC2, conventional dendritic cells type 2; CD4 CTL, cytotoxic CD4 T cells; dnT, double-negative T cells; ILC, innate lymphoid cells; PBMC, peripheral blood mononuclear cells; Platelet, platelet cluster; HSPC, haematopoietic stem and progenitor cells; Plasmablast, antibody-secreting plasmablasts; Treg, regulatory T cells.

#### High-dimensional flow cytometry of gated CD3⁺ lymphocytes shows altered T-cell subset distribution and state-marker expression in *F508del* carriers

To further characterise the immunophenotypic impact of *CFTR* mutation heterozygosity, we performed high-dimensional flow cytometry with FlowSOM clustering on gated CD3⁺ lymphocytes from healthy *F508del* carriers (n = 4) and non-carrier controls (n = 4). FlowSOM clustering with predefined metacluster resolution was used to define ten metaclusters, which upon annotation corresponded to major CD3⁺ T-cell subsets, including naïve CD4⁺ and CD8⁺ T cells, central memory CD4⁺ T cells, CD8⁺ effector memory T cells, regulatory T cells, CD8+ MAIT cells, and CD161⁺ γδ T cells (**Figure 2A)**. The gating strategy, FlowSOM metacluster tree and marker-expression patterns supporting metacluster annotation are shown in **Supplementary Figure 5**. When comparing subset abundances based on median frequencies, differences between *F508del* carriers and non-carriers became apparent. CD8+ MAIT cells in particular were reduced in carriers compared with non-carriers. However, after correction for multiple testing for all comparisons across all metaclusters, these differences were no longer statistically significant (adjusted p = 0.1940), and no other subsets showed major shifts in frequencies. Given the limited sample size and multiple testing burden inherent to this analysis, we next evaluated relative shifts in cluster representation using fold-change thresholds aligned with the FlowSOM framework. This effect size-based approach enables identification of biologically meaningful changes in cell population structure (26). Consistent with the observed fold changes (−4.23 for MAIT cells), CD8+ MAIT cells were classified as underrepresented in carriers, aligning with the reduction in circulating MAIT cells identified in the CITE-seq dataset **(Figure 2A)**. Beyond compositional differences, we next examined whether T-cell state was altered across subsets. To this end, the expression of selected activation and effector markers was compared between groups **(Figure 2B)**, with differences calculated as median fluorescence intensity (MFI) in carriers minus non-carriers for each marker-subset combination. Across multiple subsets, carriers exhibited reduced expression of CCR7 and CXCR3, whereas CCR6 expression was increased across several subsets, including CD8+ MAIT cells. In parallel, CD28 expression was generally lower, while CD39 expression was higher across multiple subsets. HLA-DR expression showed a more restricted pattern, with higher expression in CD4+ effector memory T cells and regulatory T cells, and to a lesser extent in CD8+ MAIT cells.

**Figure 2:**
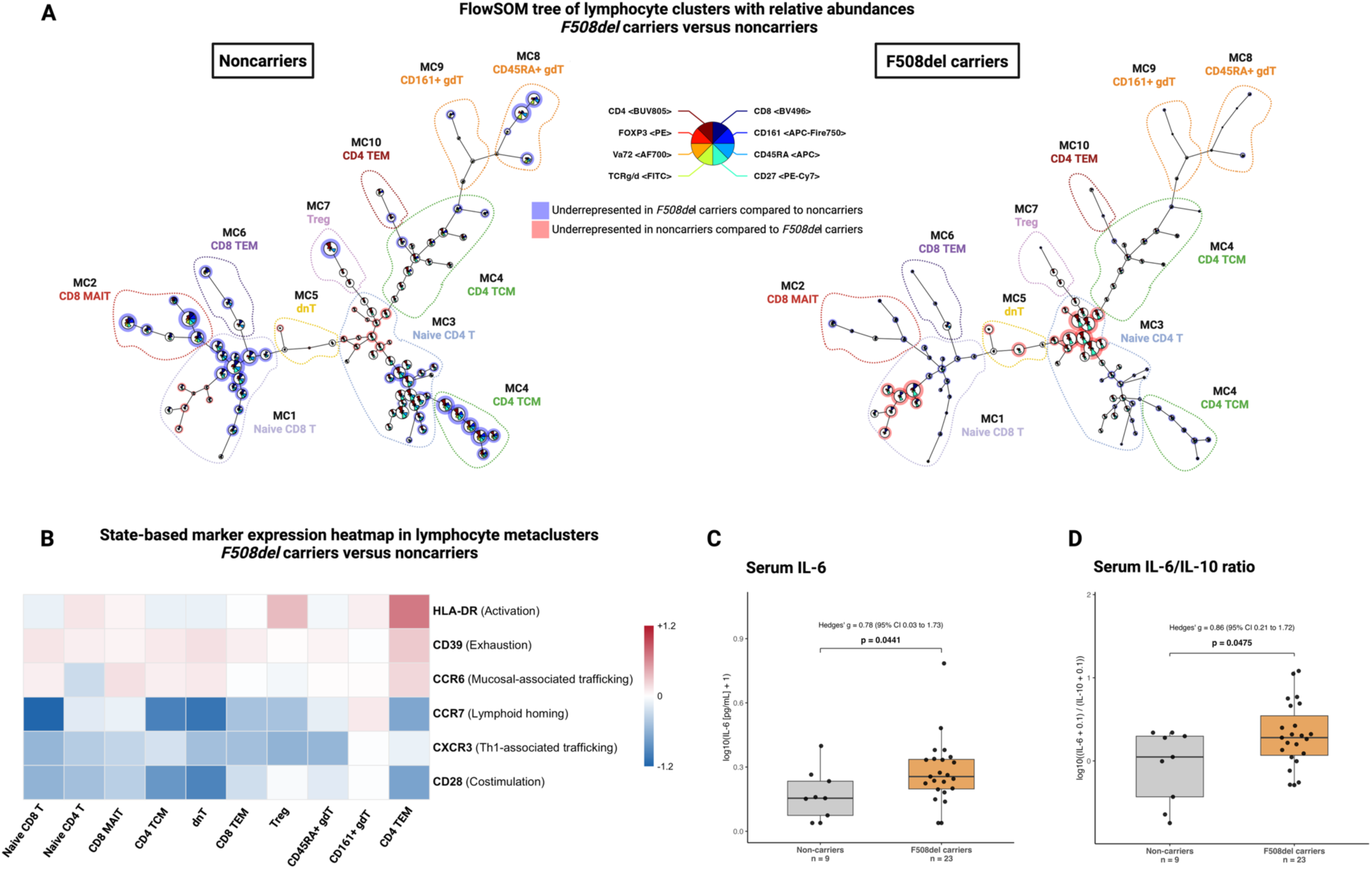
**(A) FlowSOM minimum spanning tree of CD3+ T-lymphocyte clusters derived from high-dimensional flow cytometry data**. Each node represents a FlowSOM cluster and edges indicate similarity relationships between clusters. Node size reflects the median relative abundance of the corresponding cluster within each group. Clusters are shown separately for healthy non-carriers (left) and healthy F508del carriers (right). Star charts within nodes depict the relative expression of the lineage-defining markers used for clustering (CD4, CD8, CD161, CD45RA, CD27, TCRγδ, Vα7.2, and FOXP3), as indicated in the legend. Background colouring indicates clusters with differences in relative abundance between groups based on the cluster-level statistics calculated from median frequencies: clusters with statistic values < −2.5 are coloured blue (lower abundance in *F508del* carriers compared with non-carriers), whereas clusters with statistic values > 2.5 are coloured red (lower abundance in non-carriers compared with carriers). Clusters with statistic values between −2.5 and 2.5 are shown without background colouring. The dashed outline highlights the region containing MAIT-cell clusters. **(B) Heatmap showing differences in state-marker expression across FlowSOM lymphocyte metaclusters between healthy *F508del* carriers and non-carriers**. Columns correspond to metaclusters (MC1-MC10), with MC2 representing the MAIT-cell metacluster and MC9 representing the γδ T-cell metacluster. Rows correspond to state markers included in the panel (HLA-DR, CD39, CCR6, CCR7, CXCR3, CD28). Values represent the median difference in marker expression (ΔMFI), calculated as the median expression in *F508del* carriers minus the median expression in non-carriers for each marker-metacluster combination. The colour scale is centred at zero; red indicates higher expression in carriers, blue indicates higher expression in non-carriers, and white indicates no difference. **(C) Serum IL-6 concentrations in healthy non-carriers and *F508del* carriers**. Individual points represent individual participants. Boxes indicate the interquartile range (IQR), the horizontal line indicates the median, and whiskers extend to 1.5 × IQR. Cytokine concentrations were log10-transformed before statistical analysis using log10(value + 1). **(D) Serum IL-6/IL-10 ratio in healthy non-carriers and *F508del* carriers**. Ratios were calculated from measured cytokine concentrations and subsequently log10-transformed prior to analysis using log10(value + 0.1). For Panels C-D, between-group comparisons were performed using two-sided Wilcoxon rank-sum tests. Exact p values are shown above the brackets. Effect sizes are reported as Hedges’ g with corresponding 95% confidence intervals. Sample sizes for each group are indicated below the x-axis labels. Abbreviations: MAIT, mucosal-associated invariant T cell; MFI, median fluorescence intensity; IQR, interquartile range

#### Reduced MAIT-cell abundance is associated with low-grade systemic inflammation in *F508del* carriers

Given the reduced abundance of MAIT cells observed in *F508del* carriers, we next assessed whether this alteration was accompanied by changes in circulating inflammatory mediators, consistent with the presence of low-grade inflammation, potentially of mucosal origin. To this end, serum cytokines were quantified in an expanded cohort of healthy individuals (non-carriers, n = 9; *F508del* carriers, n = 23). In line with this approach, serum IL-6 concentrations were higher in *F508del* carriers compared with non-carriers **(Figure 2C)**, with the broader cytokine profile of both groups shown in **Supplementary Figure 6**. Furthermore, the IL-6/IL-10 ratio was significantly increased in carriers, supporting an IL-6-skewed inflammatory profile. **(Figure 2D)**. Together, these findings suggest that the observed differences in T-cell subset distribution in *F508del* carriers are associated with measurable alterations in circulating IL-6.

#### Single-cell monocyte transcriptional signatures in carriers resemble inflammatory programmes in pwCF

Because classical CD14^+^ monocytes are key producers of inflammatory cytokines and have been implicated as intrinsically affected by CFTR dysfunction, we next examined whether the systemic and cellular differences observed above were reflected in transcriptional programmes within this myeloid compartment.

Differential expression analysis of classical monocytes revealed marked transcriptional changes in pwCF at baseline compared with healthy non-carriers **(Figure 3A)**. Among the most strongly upregulated genes were multiple regulators of inflammatory signalling, including *NFKBIA*, a canonical feedback inhibitor induced downstream of NF-κB activation, as well as immediate early response genes such as *JUNB* and *PIM3*. In contrast, several genes were significantly downregulated, including *CLEC12A*, an inhibitory C-type lectin receptor expressed on myeloid cells that dampens inflammatory responses.

**Figure 3.**
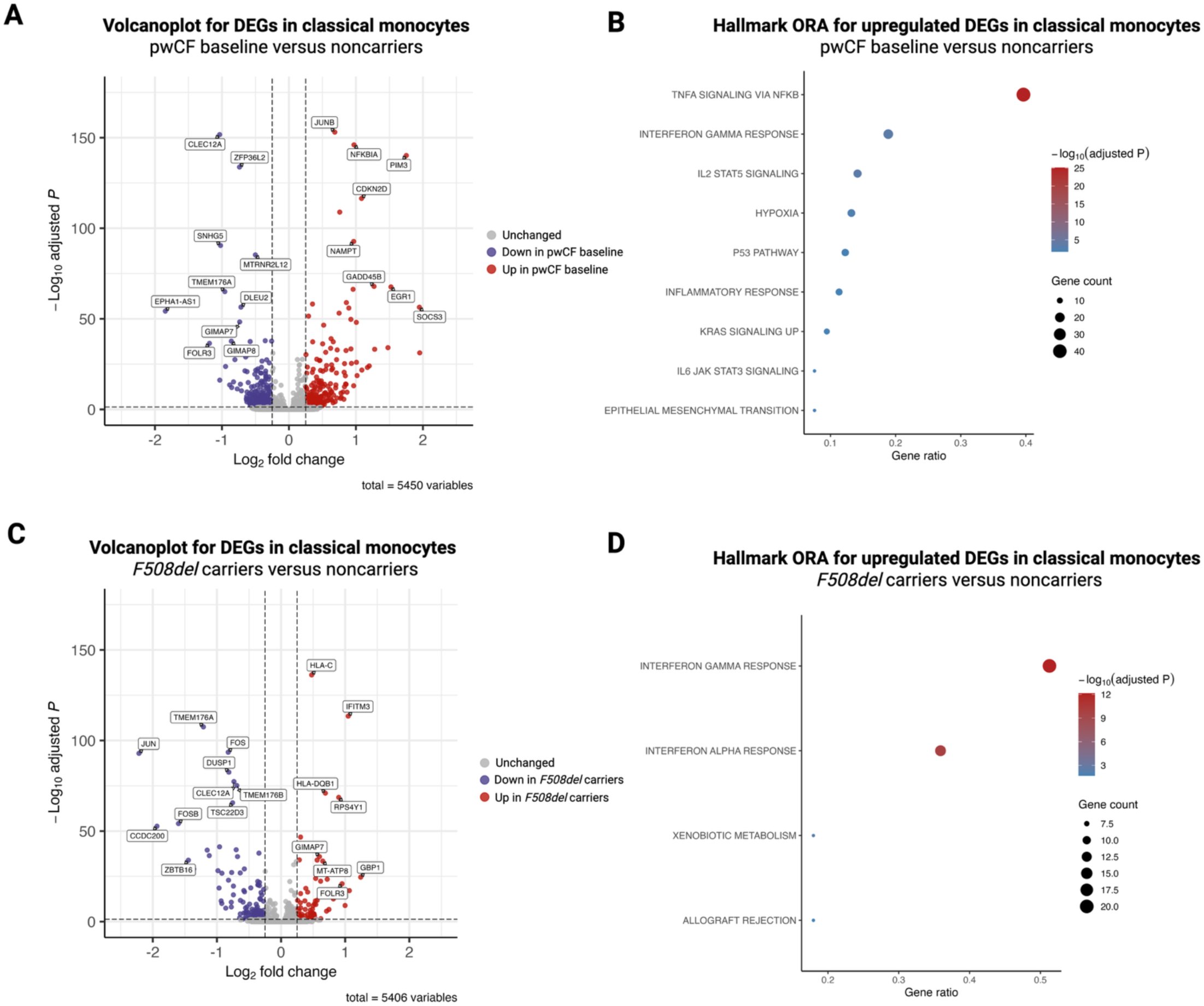
**(A) Volcano plot of differentially expressed genes in classical CD14^+^ monocytes comparing people with cystic fibrosis (pwCF) at baseline with healthy non-carriers**. Differential expression was assessed using Seurat in all genes detected in at least 10% of cells in either group. The x-axis shows log₂ fold change, and the y-axis shows −log₁₀ adjusted *P* value. Genes with adjusted *P* < 0.05 and absolute log₂ fold change ≥ 0.25 are highlighted (red, higher in pwCF at baseline; blue, higher in healthy non-carriers; grey, not significant). Labelled genes correspond to the top-ranking significantly upregulated and downregulated genes, prioritised by the product of absolute log₂ fold change and −log₁₀ adjusted *P* value. **(B) Hallmark over-representation analysis of genes upregulated in classical CD14^+^monocytes from pwCF at baseline compared with healthy non-carriers**. Enrichment analysis was performed against the MSigDB Hallmark gene set collection using the set of significantly upregulated genes from panel A, with all tested genes from the same differential expression comparison used as background. The plot shows the top enriched Hallmark terms ranked by adjusted *P* value. The x-axis represents the gene ratio, dot size indicates the number of input genes assigned to each pathway, and colour denotes −log₁₀ adjusted *P* value. (C) Volcano plot of differentially expressed genes in classical CD14^+^ monocytes comparing healthy *F508del* carriers with healthy non-carriers. Differential expression was assessed using the same strategy and thresholds as in panel A. The x-axis shows log₂ fold change, and the y-axis shows −log₁₀ adjusted *P* value. Genes with adjusted *P* < 0.05 and absolute log₂ fold change ≥ 0.25 are highlighted (red, higher in healthy *F508del* carriers; blue, higher in healthy non-carriers; grey, not significant). Labelled genes correspond to the top-ranking significantly upregulated and downregulated genes, prioritised by the product of absolute log₂ fold change and −log₁₀ adjusted *P* value. (D) Hallmark over-representation analysis of genes upregulated in classical CD14^+^ monocytes from healthy *F508del* carriers compared with healthy non-carriers. Enrichment analysis was performed against the MSigDB Hallmark gene set collection using the set of significantly upregulated genes from panel C, with all tested genes from the same differential expression comparison used as background. The plot shows the top enriched Hallmark terms ranked by adjusted *P* value. The x-axis represents the gene ratio, dot size indicates the number of input genes assigned to each pathway, and colour denotes −log₁₀ adjusted *P* value.

Hallmark over-representation analysis of genes upregulated in pwCF at baseline demonstrated enrichment of TNF-α/NF-κB signalling, interferon-γ response, IL-6/JAK-STAT3 signalling, and broader inflammatory response pathways **(Figure 3B**), consistent with a state of heightened innate immune activation in classical CD14^+^ monocytes. Comparing healthy *F508del* carriers with non-carriers, classical CD14^+^ monocytes from carriers also exhibited differential gene expression, although with smaller effect sizes **(Figure 3C)**. Notably, *CLEC12A* was again among the downregulated genes, suggesting reduced inhibitory signalling in this population. In parallel, upregulated genes included interferon- and antigen presentation-related transcripts such as *HLA-DQB1* and *IFITM3*. Over-representation analysis of genes upregulated in carriers revealed enrichment of interferon-γ and interferon-α response pathways, as well as allograft rejection-related signatures **(Figure 3D)**, which largely comprise interferon-inducible and antigen-processing genes. These pathways partially overlapped with those enriched in pwCF at baseline, albeit with lower magnitude, supporting the presence of a shared but attenuated inflammatory transcriptional programme in carriers. Extended analyses of CD14⁺ monocyte pathway enrichment, gene-level concordance between pwCF and *F508del* carriers, and representative pseudobulk gene expression are shown in **Supplementary Figure 7**. Combined, these data suggest that CFTR heterozygosity is associated with a CF-like immune phenotype at the transcriptional level, characterised by increased inflammatory and interferon-associated signalling and reduced expression of inhibitory regulators such as *CLEC12A* in classical CD14^+^ monocytes.

### Restoration of CFTR function in pwCF is associated with changes in monocyte inflammatory pathways, predicted PBMC ligand-receptor interactions and systemic cytokine levels

#### ETI attenuates TNF-associated pathways and enhances interferon-related programmes in classical monocytes

We next investigated whether partial restoration of CFTR function in pwCF through ETI in vivo modulates the inflammatory and interferon-associated programmes identified above. In classical monocytes, over-representation analysis of genes downregulated on ETI relative to baseline demonstrated enrichment of TNF-α/NF-κB signalling, IL2/STAT5 signalling, hypoxia, and IL6/JAK/STAT3 signalling **(Figure 4A)**, consistent with attenuation of inflammatory pathways that are more prominent prior to treatment. In contrast, genes upregulated on ETI were enriched for interferon-γ response and interferon-α response pathways (**Figure 4B)**, indicating a shift towards interferon-associated transcriptional programmes following CFTR modulation. Extended differential-expression, Hallmark GSEA and patient-level leading-edge gene analyses supporting these ETI-associated monocyte changes are provided in **Supplementary Figure 8**.

**Figure 4.**
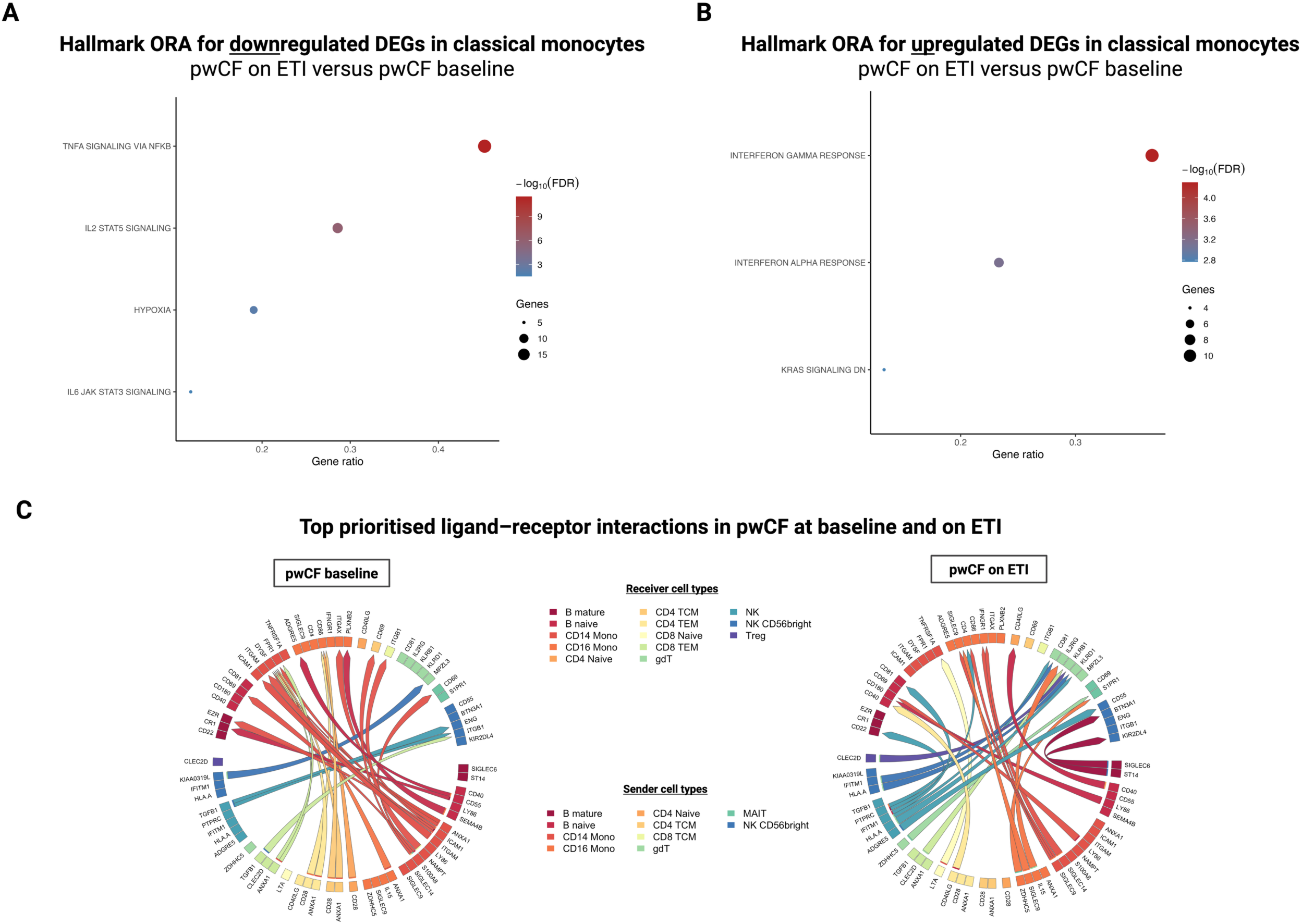
CFTR modulation is associated with changes in monocyte inflammatory pathways and predicted intercellular communication networks. **(A) Hallmark over-representation analysis (ORA) of genes downregulated in classical (CD14⁺) monocytes in people with cystic fibrosis (pwCF) on ETI compared with pwCF at baseline**. Each dot represents one Hallmark gene set. The x-axis shows the gene ratio (overlap between the input gene list and the gene set divided by the total number of genes in the gene set). Dot size indicates the number of overlapping genes, and colour represents significance (−log₁₀ false discovery rate, FDR). **(B) Hallmark ORA of genes upregulated in classical (CD14⁺) monocytes in pwCF on ETI compared with pwCF at baseline**. Each dot represents one Hallmark gene set. The x-axis shows the gene ratio. Dot size indicates the number of overlapping genes, and colour represents −log₁₀ FDR. **(C) Circos plots showing the top 25 prioritised ligand-receptor interactions per condition identified by MultiNicheNet in pwCF at baseline (left) and on ETI (right), ranked per group across sender-receiver combinations**. Each sector represents a cell type. Links between sectors represent prioritised ligand-receptor pairs connecting a sender cell type (ligand-expressing) to a receiver cell type (receptor-expressing). Link thickness is proportional to the prioritisation score.

#### ETI is associated with remodelling of predicted monocyte-derived intercellular communication

To assess whether these transcriptional changes were accompanied by alterations in intercellular immune communication, we applied MultiNicheNet to paired PBMC data from pwCF at baseline and on ETI. Prioritisation of ligand-receptor interactions across sender-receiver pairs revealed marked differences in the dominant predicted communication patterns between conditions (**Figure 4C; Supplementary Figure 9**). At baseline, prioritised interactions were dominated by CD14⁺ monocyte-derived *S100A8*-associated signalling, most prominently *S100A8-CD69* interactions directed towards multiple lymphocyte populations, including MAIT cells, Treg, γδ T cells and conventional CD4⁺ and CD8⁺ T-cell subsets. Ligand-target analysis further supported this baseline axis in MAIT cells, where *S100A8* showed high predicted ligand activity and was linked to a broad set of predicted target genes that were generally more highly expressed across baseline samples. On ETI, the dominant CD14⁺/CD16⁺ monocyte-associated prioritised interactions shifted towards *SIGLEC9-IFNGR1*, particularly involving monocyte and MAIT-cell receiver populations. These findings suggest qualitative remodelling of predicted intercellular communication following ETI, with reduced dominance of a baseline myeloid alarmin-to-lymphocyte axis and emergence of a distinct *SIGLEC9-IFNGR1*-associated communication pattern.

Taken together, these data suggest that ETI not only attenuates TNF-associated inflammatory programmes in classical CD14⁺ monocytes, but also influences predicted intercellular communication, shifting the dominant inferred CD14⁺ monocyte-derived axis from baseline *S100A8-CD69* interactions towards a distinct *SIGLEC9-IFNGR1*-associated pattern on ETI.

#### Restoration of CFTR function is associated with reduced circulating pro-inflammatory cytokines while increasing IFN-γ in a subset of pwCF

We next assessed how these cellular changes relate to changes in serum cytokine levels. Paired serum cytokine profiling was performed in adult pwCF (n = 56) before initiation of any modulator and after at least six months of ETI therapy, comparing individuals who initiated ETI as their first modulator therapy (primary-ETI, n = 27) with those who transitioned from prior TI treatment (TI-to-ETI, n = 29). The primary analyses focused on within-group changes from baseline to on-ETI, while recognising that pwCF in the TI-to-ETI group had received CFTR modulators for a longer cumulative duration (≥12 months) than those in the primary-ETI group (≥6 months on ETI). Both groups exhibited a consistent reduction in circulating TNF-α (**Figure 5A**). In primary-ETI pwCF, median TNF-α decreased from 2.97 pg/mL to 1.99 pg/mL (p < 0.05), and in TI-to-ETI pwCF from 3.67 pg/mL to 1.73 pg/mL (p < 0.001). IL-6 levels similarly decreased, from 1.31 pg/mL to 0.62 pg/mL in the primary-ETI group (p < 0.01) and from 2.35 pg/mL to 0.75 pg/mL in the TI-to-ETI group (p < 0.001). IL-13 declined significantly only in the TI-to-ETI group (0.61 to 0.42 pg/mL; p < 0.01), while remaining stable in primary-ETI pwCF. In contrast, IFN-γ increased significantly in the TI-to-ETI group (3.15 to 5.10 pg/mL; p < 0.01) but remained unchanged in the primary-ETI group. IL-17A showed the most pronounced decrease, falling from 4.11 pg/mL to 2.02 pg/mL in primary-ETI pwCF and from 29.72 pg/mL to 13.13 pg/mL in TI-to-ETI pwCF (both p < 0.0001). A similar pattern was observed for IL-8. IL-10 levels were unchanged across groups, while IL-1β decreased significantly on ETI in both cohorts (primary-ETI: 0.18 to 0.10 pg/mL; p < 0.01; TI-to-ETI: 0.22 to 0.10 pg/mL; p < 0.01). Extended adult subgroup analyses, including mean paired cytokine changes, three-timepoint trajectories in the TI-to-ETI cohort and baseline cytokine comparisons between adult primary-ETI and TI-to-ETI groups, are shown in **Supplementary Figure 11**. Overall, ETI in pwCF was associated with a broad reduction in serum pro-inflammatory cytokines, most notably TNF-α, IL-6, IL-17A, IL-8, and IL-1β, accompanied by stable IL-10 and a modest increase in IFN-γ in the TI-to-ETI subgroup. Paediatric primary-ETI cytokine trajectories and adult-paediatric comparisons are shown in **Supplementary Figure 12**.

**Figure 5.**
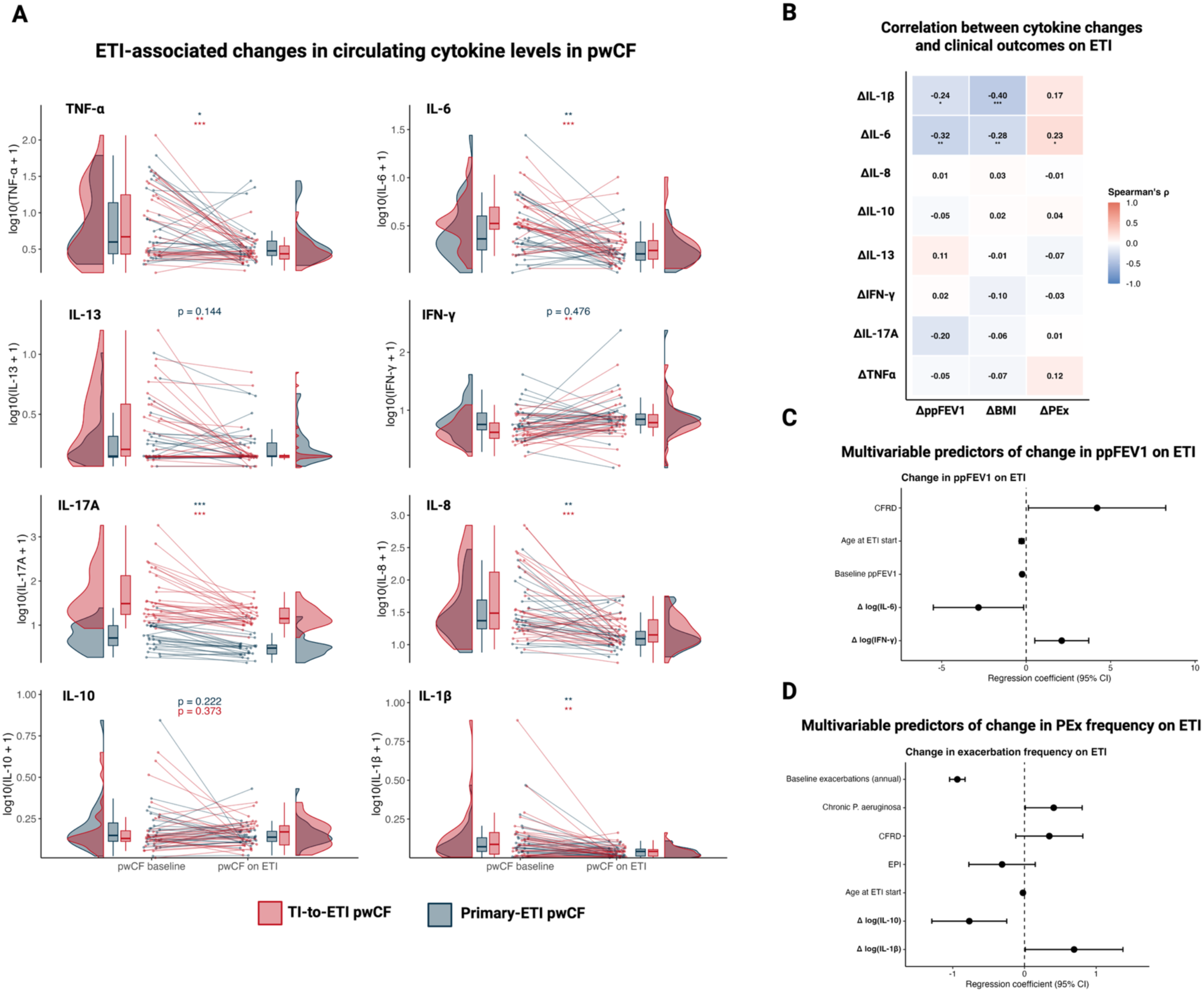
Systemic inflammatory remodelling on elexacaftor/tezacaftor/ivacaftor (ETI) and its association with clinical outcomes in cystic fibrosis. **(A) Paired analysis of circulating cytokine levels in people with cystic fibrosis (pwCF) before and after initiation of ETI. Cytokine concentrations are shown as log₁₀(x + 1)-transformed values.** Each line represents an individual patient trajectory from baseline to ETI. Data are stratified by treatment history, distinguishing individuals transitioning from tezacaftor/ivacaftor (TI-to-ETI; red) and those initiated directly on ETI (primary-ETI; blue). Violin plots depict the distribution of values at each time point, with embedded boxplots indicating median and interquartile range. Statistical comparisons were performed using two-sided Wilcoxon signed-rank tests; p-values are indicated where relevant, and significance is denoted as *p < 0.05, **p < 0.01, ***p < 0.001. Where applicable, p-values were adjusted for multiple testing using the Benjamini-Hochberg procedure**. (B) Heatmap showing Spearman rank correlation coefficients (ρ) between changes in cytokine levels on ETI and changes in clinical outcomes, including percent predicted FEV1 (ΔppFEV1), body mass index (ΔBMI), and exacerbation frequency (ΔPEx)**. Correlation coefficients are displayed within each tile and colour-coded according to magnitude and direction. Two-sided p-values were derived from Spearman correlation tests and adjusted for multiple testing using the Benjamini-Hochberg procedure. Statistical significance is indicated by asterisks (*p < 0.05, **p < 0.01, ***p < 0.001). **(C) Forest plot of a multivariable linear regression model assessing predictors of change in ppFEV1 on ETI**. Regression coefficients with 95% confidence intervals are shown for variables retained in the final model following stepwise model selection based on the Akaike information criterion. The dashed vertical line indicates no effect (β = 0). **(D) Forest plot of a multivariable linear regression model evaluating predictors of change in exacerbation frequency on ETI**. Regression coefficients with 95% confidence intervals are shown for variables retained in the final model following stepwise model selection based on the Akaike information criterion. The dashed vertical line indicates no effect (β = 0). **Abbreviations:** ETI, elexacaftor/tezacaftor/ivacaftor; TI, tezacaftor/ivacaftor; pwCF, people with cystic fibrosis; ppFEV₁, percent predicted forced expiratory volume in one second; BMI, body mass index; PEx, pulmonary exacerbation frequency; Δ, change between baseline and on ETI; TNF-α, tumour necrosis factor alpha; IFN-γ, interferon gamma; IL, interleukin.

#### Cytokine changes in pwCF on ETI are associated with improvements in clinical endpoints

To evaluate the clinical relevance of ETI-associated cytokine changes, we analysed their relationships with longitudinal changes in ppFEV₁, BMI and exacerbation frequency. In bivariate correlation analysis, reductions in IL-1β and IL-6 were associated with increases in ppFEV₁ and BMI, while IL-6 reduction also correlated with changes in exacerbation frequency (**Figure 5B**). Baseline clinical and inflammatory features associated with cytokine changes, together with a model of BMI change, are shown in **Supplementary Figure 13**. In multivariable regression, larger reductions in IL-6 and increases in IFN-γ were associated with ppFEV₁ gain (**Figure 5C**). In the exacerbation model, IL-10 and IL-1β changes were retained alongside clinical covariates, supporting a relationship between systemic cytokine remodelling and changes in exacerbation burden (**Figure 5D**).

## DISCUSSION

This study maps systemic immunity across the spectrum of CFTR dysfunction, showing that (i) healthy *F508del* carriers display a CF-like immunophenotype, and (ii) partial restoration of CFTR function in pwCF is associated with broad attenuation of inflammatory signalling, enrichment of interferon-associated programmes, and remodelling of in-silico predicted intercellular communication. By combining CITE-seq, high-dimensional flow cytometry, multiplex serum cytokine analysis, and ligand-receptor inference modelling, we generated a multimodal characterisation of how CFTR dysfunction shapes systemic immunity and how these alterations can be modulated through ETI. Beyond these disease-specific observations, our findings support a broader conceptual shift in how CFTR dysfunction is understood. Rather than a strict binary distinction between affected individuals and unaffected carriers, our data suggest that CFTR function exists along a continuum reflected in systemic immune phenotypes. This aligns with emerging perspectives that challenge strictly Mendelian frameworks and instead emphasise graded quantitative effects of genetic variation on human phenotypes (27).

Inclusion of heterozygous *F508del* carriers allowed us to examine whether one pathogenic *CFTR* variant is sufficient to influence systemic immunity. Carriers exhibited elevated serum IL-6 levels together with increased IL-6/IL-2 and IL-6/IL-10 ratios, consistent with a subtle but measurable pro-inflammatory bias. At the cellular level, asymptomatic carriers demonstrated reduced circulating MAIT-cell abundance together with inflammatory and interferon-associated transcriptional programmes in classical monocytes. Exploratory ligand-receptor inference in healthy controls further suggested altered monocyte-centred predicted communication patterns in *F508del* carriers, although these analyses were hypothesis-generating given the small single-cell control cohort (**Supplementary Figure 10**). These findings extend prior epidemiological and transcriptomic observations linking *CFTR* heterozygosity to increased risk of bronchiectasis, pancreatitis, respiratory infections, and chronic respiratory disease (20–24). Consistent with this concept, population-scale phenome-wide association analyses in UK Biobank further support clinically relevant respiratory and systemic phenotypic consequences of *F508del* carriership, including associations with bronchiectasis, pancreatitis, and asthma (28). Importantly, our data provide evidence that *CFTR* heterozygosity is associated with measurable systemic immune remodelling even in the absence of overt disease. Taken together, these data challenge the traditional notion of ‘silent’ carriership in autosomal recessive disorders and support the concept that *CFTR* heterozygosity represents a biologically relevant state of altered immune homeostasis.

Among circulating immune populations, MAIT cells emerged as the most consistently altered compartment across the spectrum of CFTR dysfunction. MAIT cells contribute to mucosal immune surveillance and rapidly produce IFN-γ and IL-17A in response to microbial metabolites (29,30). We observed persistent reductions in circulating MAIT-cell abundance in both pwCF and healthy *F508del* carriers, extending prior observations in CF. Reduced MAIT-cell frequencies have also been associated with chronic airway disease and infection risk in other conditions characterised by CFTR dysfunction or mucosal inflammation (31,32). While redistribution of MAIT cells from blood to inflamed tissues has historically been proposed as a mechanism for peripheral depletion, alternative explanations including activation-induced apoptosis or impaired maintenance have also been suggested. Notably, Bardin et al. reported partial restoration of circulating MAIT-cell numbers after ETI in a predominantly paediatric cohort with limited chronic *P. aeruginosa* colonisation (33), whereas persistent reduction was observed in our adult cohort with established chronic colonisation. Given the known association between reduced circulating MAIT-cell abundance and chronic *P. aeruginosa* infection in CF (31), ETI-mediated restoration of CFTR function alone may not be sufficient to normalise MAIT-cell homeostasis in the setting of longstanding airway inflammation and chronic infection.

At the transcriptomic level, ETI was associated with attenuation of inflammatory monocyte programmes together with enrichment of interferon-associated signalling. Classical monocytes from pwCF on ETI demonstrated downregulation of TNF-NF-κB, IL6/JAK/STAT3, and hypoxia-associated pathways alongside upregulation of interferon-stimulated genes including *ISG15*, *IFI44L*, and *MX1*. These findings align with prior studies suggesting impaired or blunted interferon responses in CF (34–36) and support the concept that ETI may partially restore antiviral or interferon-associated immune competence in circulating myeloid cells. In parallel, paired serum cytokine profiling demonstrated broad reductions in IL-6, IL-8, IL-17A, TNF-α, and IL-1β following ETI, consistent with attenuation of systemic inflammatory signalling beyond the airway compartment (17–19). Reductions in IL-6 and IL-1β correlated with clinically relevant improvements, including gains in ppFEV₁ and BMI. In retained multivariable models, IL-6 reduction and IFN-γ increase were associated with ppFEV₁ gain, whereas IL-1β reduction was associated with BMI gain. These associations support a link between systemic inflammatory remodelling and clinical improvement.

Strikingly, IFN-γ levels increased significantly in the TI-to-ETI group, who had more advanced disease than the primary-ETI group at baseline, and this rise was associated with a modest ppFEV₁ improvement. Given the central role of IFN-γ in immunoregulation and antiviral defence, this pattern may reflect enhanced interferon-associated immune activity following partial CFTR correction, complementing the interferon-stimulated gene signature observed in classical CD14^+^ monocytes. Notably, this increase in IFN-γ occurred alongside a reduction in IL-13 in the same subgroup, raising the possibility of a broader shift away from type 2-associated inflammatory signalling towards a more interferon-associated immune state after ETI. Given the partially reciprocal relationship between interferon- and type 2-associated immune programmes, including cross-regulatory effects between IFN-γ- and IL-13-associated pathways (37,38), this pattern may reflect qualitative immune rebalancing rather than generalized immunosuppression. Together, these findings suggest that ETI exerts both anti-inflammatory and immunorestorative effects in pwCF. Importantly, these observations indicate that highly effective CFTR modulation may reshape immune signalling networks rather than simply suppress inflammatory activity.

To further dissect how ETI remodels intercellular communication, we applied the MultiNicheNet framework (39) to paired PBMC samples from pwCF before and on ETI. At baseline, prioritised ligand-receptor interactions were dominated by CD14⁺ monocyte-derived *S100A8*-associated interactions, particularly *S100A8-CD69* links directed towards lymphocyte populations. This was further supported by ligand-target inference in MAIT cells, where *S100A8* showed high predicted ligand activity and was linked to a set of predicted target genes that were broadly upregulated across evaluable baseline samples. S100A8 and S100A9 form calprotectin, the protein complex historically described as the “cystic fibrosis antigen” in early studies from the 1970s, predating its molecular characterisation (40,41). While initially identified in neutrophils, S100A8/A9 is also abundantly expressed by activated classical monocytes, supporting its role as a mediator of monocyte-driven inflammatory signalling in this context (42). In line with this interpretation, S100A8/S100A9 has been identified as an endogenous ligand for CD69, supporting a plausible link between myeloid alarmin release and CD69-expressing lymphocyte populations (43). Following ETI, the dominant prioritised myeloid-associated interactions shifted towards *SIGLEC9-IFNGR1*, particularly involving CD14⁺/CD16⁺ monocyte sender populations and monocyte or MAIT-cell receivers. Although the inferred *SIGLEC9-IFNGR1* interaction should not be interpreted as direct receptor-level signalling, its emergence may provide insight into the altered immune state on ETI. SIGLEC9 belongs to the Siglec family of sialic-acid-binding receptors, which regulate immune-cell functions through glycan recognition in infectious disease, inflammation, autoimmunity and cancer (44). IFNGR1, by contrast, forms part of the IFN-γ receptor system, through which IFN-γ coordinates transcriptional programmes with central roles in macrophage activation, antigen presentation and host defence (45). The combination of reduced *S100A8-CD69* prioritisation and increased *SIGLEC9-IFNGR1* prioritisation therefore suggests a shift from baseline damage-associated myeloid-to-lymphocyte communication towards a monocyte-centred network in which IFN-γ-responsive cells are positioned within a more immunoregulatory glycan-recognition context. This interpretation fits the broader observation that ETI reduced inflammatory cytokines and *TNF/NF-κB*-associated monocyte programmes while increasing interferon-associated signatures, supporting immune rebalancing rather than generalised immunosuppression.

This study has several notable strengths. We combined longitudinal sampling in well-characterised pwCF with multimodal immune profiling across transcriptomic, cellular, and circulating protein levels, while also incorporating genotyped healthy *F508del* carriers and non-carriers. This design enabled us to characterise systemic immune alterations across different levels of CFTR functionality and to relate immune remodelling to clinically meaningful outcomes following ETI therapy. Several limitations should also be acknowledged. First, the number of individuals included for CITE-seq and flow cytometry was modest, and larger independent cohorts will be required to validate and refine these observations. Nevertheless, the dataset provides a unique high-resolution multimodal characterisation of circulating immunity across distinct states of CFTR dysfunction in vivo. Second, we focused primarily on systemic rather than airway immune responses, although prior studies have demonstrated local anti-inflammatory effects of ETI within the airway compartment (17,18). Third, the MultiNicheNet analyses are predictive and require experimental validation. Fourth, functional studies will be required to determine whether the observed interferon-associated transcriptional programmes and inferred ligand-receptor communication changes translate into altered pathogen responses, antiviral immunity, immune regulation, or long-term disease modification. Finally, the *F508del CFTR* variant, used in this study to evaluate the consequences of reduced CFTR dosage on immune processes, does not capture the full breadth of molecular effects of *CFTR*-variants. Future studies, incorporating larger cohorts of heterozygous carriers, should also include other functional classes.

More broadly, our findings support reconsideration of the prevailing all-or-nothing view of CFTR dysfunction. Through multimodal mapping of systemic immunity, we show that apparently healthy *F508del* carriers resemble pwCF more closely than non-carrier controls across several immune parameters, including circulating cytokine profiles, MAIT-cell abundance, and monocyte inflammatory programmes. These observations suggest that even partial reduction in CFTR activity may be sufficient to perturb immune homeostasis and provide a potential biological framework linking CFTR heterozygosity to increased susceptibility to chronic inflammatory and respiratory disease. One potential mechanism underlying heterogeneity among carriers may relate to allele-specific expression. Recent work in inborn errors of immunity has shown that autosomal random monoallelic expression can influence disease penetrance, whereby preferential expression of either the wild-type or mutant allele shapes cellular phenotype despite an identical germline genotype (46). Although this mechanism has not been demonstrated for CFTR in immune cells, it provides a conceptual framework through which heterozygous CFTR variants could produce variable effective CFTR dosage across individuals or cell types. Future studies assessing allele-specific CFTR expression in sorted innate immune populations, particularly monocytes and neutrophils, may help determine whether such mechanisms contribute to the inflammatory carrier phenotypes observed here. At the same time, our data position ETI as both an anti-inflammatory and immunorestorative therapy, highlighting how pharmacological restoration of CFTR activity has the potential to influence systemic immune signalling in vivo.

In conclusion, this study demonstrates that CFTR dysfunction is associated with systemic immune remodelling across a spectrum of CFTR activity. Healthy *F508del* carriers exhibited a distinct CF-like immune phenotype characterised by low-grade systemic inflammation, reduced circulating MAIT cells, and inflammatory monocyte transcriptional programmes, while ETI therapy in pwCF was associated with attenuation of inflammatory signalling, enrichment of interferon-associated monocyte programmes, and remodelling of predicted intercellular communication. By leveraging CF as a modulable model, these findings advance our understanding of systemic immunity and inflammation across the spectrum of CFTR dysfunction in humans.

## ONLINE METHODS

### Study Design and Participants

Participants were enrolled between 1 March 2021 and 31 December 2024 at the CF Reference Centre of Ghent University Hospital, Belgium, part of the Belgian CF Registry (BCFR, Sciensano, EC/2010/226). Blood samples from 229 people with cystic fibrosis (pwCF), were prospectively biobanked during routine follow-up after obtaining written informed consent (BR-151, B.U.N. 6702020001009; FAGG BB210012). The study was approved by the local ethics committee of Ghent University Hospital (ONZ-2022-0295).

Samples from pwCF were collected for inclusion in the biobank at three defined timepoints: baseline (n = 151), after at least six months of dual-modulator therapy (tezacaftor-ivacaftor, TI; n = 69) and following at least six months of triple-modulator therapy (ETI; n = 143). All pwCF who received TI were subsequently transitioned to ETI. Inclusion criteria for the TI-to-ETI cohort required that participants have samples at each of the three timepoints (baseline, on TI, and on ETI). For the cohort treated directly with ETI (primary-ETI), inclusion required samples at baseline and during ETI treatment.

For the serum cytokine analysis, paired serum samples were available for 90 pwCF who had been treated with CFTR modulators for at least six months. These individuals were stratified into two clinically defined groups: primary-ETI, comprising pwCF who initiated ETI as their first CFTR modulator therapy, and TI-to-ETI, consisting of pwCF who had previously received TI and subsequently transitioned to ETI. This stratification reflects real-world clinical practice, as treatment decisions were made independently of study enrolment and were determined by drug availability and clinical indication. The duration of therapy at the time of sampling was 7.6 months (IQR 6.3-8.3) on TI and 7.4 months (IQR 6.7-8.3) on ETI. For cross-sectional comparison, serum samples from 23 *F508del* carriers and nine non-carriers were recruited, primarily from family members of pwCF. Carriers were confirmed through genetic screening to harbour a single *F508del* mutation on one allele and no additional mutations among the 50 most frequent *CFTR* variants. Healthy controls were frequency-matched to the pwCF cohort by age and sex; clinical characteristics of these participants are summarised in **Table 1**.

A subset of eight pwCF homozygous for *F508del* was selected from the biobank for single-cell sequencing and high-dimensional flow cytometry. All eight had paired PBMC samples collected at baseline (pre-treatment) and after at least six months on CFTR modulator therapy. Six of these participants belonged to the TI-to-ETI group, and two were primary-ETI pwCF, with treatment durations of 7.6 months (IQR 6.4-8.0) on TI and 7.0 months (IQR 6.4-7.4) on ETI. Eight healthy controls - four *F508del* carriers and four non-carriers were also included in these experiments.

All pwCF were clinically stable at the time of sampling, defined by the absence of pulmonary exacerbation and no antibiotic use in the preceding four weeks. Healthy controls completed a structured health questionnaire confirming the absence of chronic disease, active infection, regular medication use, or recent vaccination or dental procedures.

### Sample Collection

Peripheral blood was collected by venepuncture into EDTA and serum tubes and processed within one hour. Serum and plasma were isolated by centrifugation and stored at -80 °C until analysis. PBMCs were isolated using the standard Ficoll-Hypaque technique with Leucosep tubes prefilled with Lymphoprep (Serumwerk Bemburg AG), washed, assessed for viability, and cryopreserved in freezing medium at -150 °C.

### Cellular Indexing of Transcriptomes and Epitopes by Sequencing (CITE-seq)

#### Experimental procedures

Cryopreserved PBMCs were rapidly thawed in a 37 °C water bath, washed once in PBS with 0.04 % BSA, counted by trypan-blue exclusion (viability ≥ 85 %), and Fc-blocked with TruStain FcX (BioLegend) for 10 min on ice. Cells were stained on ice for 30 min with a 1:250 dilution of TotalSeq™ C antibody cocktail (BioLegend Panel P0042) and oligonucleotide-conjugated hashing antibodies (Hashtags 14 & 15; BioLegend) in PBS + 0.04 % BSA, washed twice, filtered through a 40 µm strainer, then stained with DAPI (1 µg/mL) for live/dead discrimination.

Live DAPI⁻ cells were sorted on a BD FACS Melody (BL-2 configuration). From each sample, 100 000 events were collected into RPMI-1640 + 10 % FCS; post-sort viability consistently exceeded 90 %. Cells were pelleted, resuspended at \∼1 × 10^6 cells/mL in PBS + 0.04 % BSA, and immediately loaded for encapsulation.

Approximately 17 000 cells per sequencing lane were processed on a Chromium Single-Cell A Chip (10x Genomics, 5′ v2 Dual Index, SI-TT-A11) using the 10x Genomics Chromium Controller. GEM generation and reverse transcription were performed with the Single Cell 5′ Library & Gel Bead Kit v2 (53 °C for 45 min; 85 °C for 5 min). After GEM breakage, barcoded cDNA was purified with Dynabeads MyOne SILANE and amplified by PCR (13 cycles). The product was split for gene-expression library construction (14 PCR cycles) and feature-barcode (ADT/HTO) library construction (8 cycles) using the Single Cell 5′ Feature Barcode Library Kit. Libraries were quantified by Qubit dsDNA HS assay, sized on an Agilent Bioanalyzer High Sensitivity DNA chip, and verified on an Advanced Analytics Fragment Analyzer. ADT and hashing libraries were pooled at 5 % of total molarity; GEX libraries comprised the remaining 95 %.

Sequencing was performed on an Illumina NovaSeq 6000 (paired-end 28 × 91 bp) across 15 S4 lanes over two days (eight lanes on day 1; seven on day 2), with two dual-indexed samples per lane. Demultiplexing used bcl2fastq and data were processed with Cell Ranger v7.1.0 against GRCh38-2020-A (intronic reads included). Across all lanes, estimated cell recoveries ranged from ∼12 000 to ∼25 000 per lane, mean reads per cell from ∼20 000 to ∼37 000, and sequencing saturation from 64 % to 78 %. Median genes detected per cell ranged from ∼1 000 to ∼1 700 and median UMIs per cell from ∼2 000 to ∼4 100. Genome mapping rates exceeded 94 % (63-66% exonic; 6-8 % intronic), with 85-96 % of reads assigned to cells. Antibody libraries achieved 29-36 % saturation, with 65-75 % of reads usable per cell and a median of ∼2 800-4 200 ADT UMIs per cell.

#### Data processing and analysis

Raw BCL files from paired single-cell gene expression and antibody-derived tag (ADT) libraries were demultiplexed using *bcl2fastq* and processed with the Cell Ranger *count* pipeline (v7.1.0) against the GRCh38 reference, using Feature Barcoding to generate UMI count matrices for RNA and ADT modalities. Count matrices were analysed in Seurat (v5). Ambient RNA contamination was corrected using *decontX*. Cells with low or extreme gene counts, high mitochondrial transcript fractions, or aberrant UMI counts were excluded based on dataset-specific thresholds. The RNA assay was log-normalized, and the ADT assay was normalized using centred log-ratio (CLR) transformation. Highly variable genes were identified with *FindVariableFeatures*, and data were scaled using *ScaleData*. Principal component analysis (PCA) was performed separately for RNA and ADT assays, and donor-level batch effects were corrected using Harmony on PCA embeddings. Multimodal integration was performed using weighted nearest neighbour (WNN) analysis (*FindMultiModalNeighbors*), followed by joint UMAP embedding (*RunUMAP*) and clustering (*FindClusters*). Cell-type annotation was performed by reference mapping using Azimuth (*MapQuery*) at the level 2 (l2) PBMC taxonomy (47). MAIT cell proportions were calculated on a per-sample basis and analysed using linear mixed-effects models (LMMs), ensuring statistical inference at the level of biological replicates rather than individual cells. This approach accounts for repeated measurements within individuals and provides robust estimates in the presence of unbalanced group sizes across conditions. Condition was included as a fixed effect, and patient identity was modelled as a random intercept. Estimated marginal means were computed using the *emmeans* package and pairwise contrasts between conditions were performed with Benjamini-Hochberg correction for multiple testing.

For differential expression analyses presented in the main figures, cell-level comparisons were performed within Seurat using *FindMarkers*, restricting to genes expressed in at least 10% of cells in either group. For cross-sectional comparisons (e.g. pwCF at baseline versus healthy non-carriers, and *F508del* carriers versus non-carriers), Wilcoxon rank-sum tests were used. For longitudinal comparisons within pwCF (pre- versus on ETI), the MAST framework was applied with patient identity included as a latent variable to account for repeated measures. P-values were adjusted using the Benjamini-Hochberg method, and genes with adjusted P < 0.05 and absolute log₂ fold change ≥ 0.25 were considered significant. Over-representation analysis (ORA) was performed separately for genes upregulated and downregulated in each comparison using the MSigDB Hallmark gene set collection (via *msigdbr*). Background gene sets consisted of all genes tested in the corresponding differential expression analysis. Enrichment significance was assessed using Fisher’s exact test with multiple testing correction, and pathways with adjusted P < 0.05 were considered significant.

Cell-cell communication analysis was performed using the MultiNicheNet framework (39). A prior knowledge ligand-receptor network and ligand-target regulatory matrix were obtained from publicly available resources and mapped to HGNC gene symbols. The filtered Seurat object was converted to a SingleCellExperiment, and metadata including condition, sample, patient identity and cell type were incorporated. For the paired pwCF baseline versus on-ETI analysis, patient identity was included as a covariate to account for repeated sampling. Analyses were restricted to cell types represented across conditions and to genes expressed in at least 5% of cells in at least 50% of samples per condition. Pseudobulk expression profiles were computed per gene, cell type and sample. Differential expression between conditions was calculated for each sender and receiver cell type, and ligand activities were inferred based on the ability of candidate ligands to explain observed transcriptional changes in receiver populations using the ligand-target prior model. Ligand-receptor interactions were prioritised by integrating ligand activity, ligand expression, receptor expression, expression prevalence, cell-type specificity and condition specificity. Prioritisation was performed per condition, allowing identification of condition-associated predicted communication axes. MultiNicheNet outputs were visualised using circos plots of top-ranked interactions per condition, sender-focused ligand-receptor evidence-layer plots, and ligand-target visualisations for selected receiver populations. It should be noted that ligand-receptor pairs are derived from a prior knowledge network and represent ligand-driven regulatory hypotheses constrained by receptor expression, rather than direct evidence of receptor-mediated signalling.

### Flow Cytometry

#### Experimental procedures

After CITE-seq sorting, leftover PBMCs (∼2-5 × 10⁶ cells/sample) were washed and counted. For each donor, ∼1 × 10⁶ cells were allocated to the T/NK panel and 1 × 10⁶ to the B/DC panel. To track instrument performance over the multi-day acquisitions, Rainbow Calibration Particles (BD Biosciences) were run daily, and PMT voltages were adjusted only if bead median fluorescence intensities drifted by >5%. Single-stained compensation controls were generated using BD CompBeads (BD Biosciences) stained with each antibody individually; spillover matrices were calculated in FlowJo v10.9 and applied to all samples.

All staining was performed at 4 °C unless otherwise noted. Cells were first blocked with human Fc block and labelled with Zombie viability dye for 30 min, protected from light. For the T/NK panel, cells were then incubated for 30 min with a first surface-antibody mix containing CD8, HLA-DR, CD4, CD16, CD56, CD27 and biotin-Vα24-Jα18. After washing, cells were incubated overnight with a second surface-antibody mix containing CD278/ICOS, CCR7, CD95, CXCR3, CXCR5, CTLA-4/CD152, CD39, CD25, CD28, TCRγδ, PD-1, CD134/OX40, CCR6, streptavidin, CD45RA, Vα7.2 and CD161. The following day, cells were washed, fixed and permeabilised for 30 min, followed by intracellular staining for CD3 (BUV395) and FoxP3 (PE) for at least 1 h.

For the B/DC panel, cells were stained in an analogous sequential workflow. After viability staining and Fc blocking, cells were incubated for 30 min with a first surface-antibody mix containing CD24, HLA-DR, CD16, IgD, CD11c, CD14, CD20, CD19, CD141 and CD27. After washing, cells were incubated overnight with a second surface-antibody mix containing CXCR3/CD183, CD86, CCR7, CD1c, CD25, CD169, CD123, PD-1, CD94, CD21, CD64, IgM and CD38. The following day, cells were washed, fixed and permeabilised for 30 min, followed by intracellular CD3 staining (BUV805) for at least 1 h. After final washes, samples were acquired on a BD FACSymphony using FACSDiva software, recording ≥50,000 live singlets per tube.

#### Data processing and analysis

Raw FCS files were imported into R (v4.4.0) using the flowCore package, preserving full dynamic range. Within-sample quality control included removal of margin events (PeacoQC::RemoveMargins), exclusion of doublets based on FSC-A versus FSC-H (PeacoQC::RemoveDoublets), and gating of live singlets using a fixed polygon gate on FSC-A versus viability dye. Manual gating boundaries for lymphocytes, singlets, live cells, and CD3⁺ populations were defined in FlowJo v10.9, exported via CytoML (open_flowjo_xml), and cross-validated with automated gating to ensure consistency across samples. All downstream analyses were performed on gated CD3⁺ lymphocytes. Compensation matrices were initially calculated from single-stained controls in FlowJo, manually reviewed and adjusted where required, and subsequently applied to the raw FCS files before downstream analysis. Fluorescence channels were transformed using logicle transformation (estimateLogicle), and scatter channels were linearly rescaled (linearTransform).

High-dimensional cytometry data were analysed using the FlowSOM workflow as described by Quintelier et al (26). To correct for inter-day batch effects, CytoNorm was trained on healthy control samples from each acquisition day using lineage and state markers, and the resulting normalisation functions were applied to all samples. Pre-and post-normalisation distributions were evaluated using density plots and UMAP visualisation to confirm effective removal of technical variation without distortion of biological signal. Normalised flow frames were aggregated into a single downsampled flowFrame using FlowSOM::AggregateFlowFrames, and unsupervised clustering was performed on canonical lineage markers using FlowSOM without prior dimensionality reduction, scaling all channels equally. A self-organising map grid of 10 × 10 nodes was generated, followed by metaclustering into ten clusters, selected based on the expected number of biologically relevant T-cell subsets. Cluster abundances (relative frequencies) and median fluorescence intensities (MFIs) of state markers were extracted using FlowSOM::GetFeatures.

Differences in cluster abundance between groups were quantified using FlowSOM::GroupStats, which computes group-wise medians, fold changes, and associated p-values. Statistical comparisons between *F508del* carriers and non-carriers were performed using unpaired Wilcoxon rank-sum tests, with p-values adjusted for multiple testing using the Benjamini-Hochberg procedure. For visualisation of differential abundance on FlowSOM trees, clusters were classified based on fold-change thresholds as implemented in the FlowSOM workflow: clusters with fold-change values < −2.5 were classified as underrepresented in *F508del* carriers, whereas clusters with fold-change values > 2.5 were classified as underrepresented in non-carriers, and clusters with fold-change values between −2.5 and 2.5 were not considered differentially represented. State-marker expression differences between groups were assessed by comparing MFIs per cluster and visualised as differences (*F508del* carriers minus non-carriers).

### Serum Cytokine Profiling

#### Experimental procedures

Serum concentrations of inflammatory cytokines were measured using multiplex electrochemiluminescence assays on the Meso Scale Discovery (MSD) V-PLEX platform (Meso Scale Diagnostics, Rockville, Maryland). Two commercial kits were employed: the Human Proinflammatory Panel 1 (K15049D-1; IFN-γ, IL-1β, IL-2, IL-4, IL-6, IL-8, IL-10, IL-13, TNF-α) and the Human Cytokine Panel 1 (K15050D-1; GM-CSF, IL-5, IL-17A).

Serum samples were thawed on ice and assayed in singlicate (consistent with manufacturer recommendations), with a two-fold in-plate dilution prepared by mixing 25 µL of sample with 25 µL of assay diluent (Diluent 2 for the proinflammatory panel, Diluent 43 for the cytokine panel). Plates pre-coated with capture antibodies were incubated with 50 µL per well of calibrators or diluted samples for 2 hours at room temperature on a plate shaker (≈650 rpm). Following three washes with PBS-Tween 20 (0.05%), 25 µL per well of SULFO-TAG conjugated detection antibodies was added and incubated for 2 hours under the same conditions. After a final wash, 150 µL per well of 2× MSD Read Buffer T was applied immediately before reading on the SQ120MM instrument.

Eight-point standard curves were generated from lyophilised calibrator blends reconstituted and serially diluted according to the manufacturer’s instructions. Analyte concentrations were back-calculated using MSD Discovery Workbench software (v4.0) with a four-parameter logistic fit and 1/y² weighting. Values below the lower limit of detection were assigned the plate-specific LLOD divided by two. All concentrations were subsequently log₁₀-transformed using a pseudocount of one (log₁₀[pg/mL + 1]) for downstream analyses.

#### Data processing and statistical analysis

All statistical analyses were conducted in R (v4.4.0; R Foundation for Statistical Computing, Vienna, Austria) within the RStudio IDE (v2026.01.0+392). Raw paired serum cytokine concentrations were imported from Excel and matched to clinical metadata. Categorical covariates (sex, *F508del* homozygosity, cystic fibrosis-related diabetes (CFRD), cystic fibrosis-related liver disease (CFLD), exocrine pancreatic insufficiency (EPI), and chronic *Pseudomonas aeruginosa* infection at baseline) were encoded as factors. Continuous variables (post-bronchodilator percent predicted FEV₁ (ppFEV₁), body mass index (BMI), annual exacerbation frequency, and age at ETI initiation) were visually inspected for distributional properties.

To account for right-skewed cytokine distributions, log₁₀ transformation with a pseudocount of one was applied to both baseline and on-ETI values. Treatment-associated changes were defined as Δlog₁₀ = log₁₀(on-ETI + 1) - log₁₀(baseline + 1). Paired comparisons of cytokine levels before and after ETI were performed using the Wilcoxon signed-rank test. Where multiple cytokines were evaluated simultaneously, P-values were adjusted using the Benjamini-Hochberg method.

Associations between cytokine changes (Δlog₁₀) and clinical outcomes (ΔppFEV₁, ΔBMI, Δexacerbation frequency) were assessed using Spearman rank correlation. Resulting P-values were adjusted for multiple testing using the Benjamini-Hochberg procedure, and only associations remaining significant after correction (adjusted P < 0.05) were considered in downstream interpretation and visualisation.

To further evaluate the relationship between cytokine changes and clinical outcomes, multivariable linear regression models were constructed with change in clinical endpoints (ΔppFEV₁, ΔBMI, or Δexacerbation frequency) as dependent variables. Candidate predictors included demographic and clinical covariates (age at ETI initiation, sex, genotype, baseline disease severity measures, and comorbidities) together with cytokine changes (Δlog₁₀ values). Final models were derived using stepwise selection based on the Akaike Information Criterion (stepAIC).

Model assumptions were assessed by examining variance inflation factors, residual distributions, homoscedasticity, and influential observations using the performance package. Model fit was summarised using R² values.

### Data and code availability

The CITE-seq data generated in this study will be deposited in the Gene Expression Omnibus (GEO) repository and made publicly available upon publication. Flow cytometry data will be deposited in FlowRepository, a public repository for annotated flow cytometry datasets compliant with MIFlowCyt standards. De-identified serum cytokine data, including both raw and processed values and accompanying metadata, will be deposited in Zenodo and made publicly available upon publication, in accordance with applicable data protection regulations. All code used to perform the analyses and generate the figures will be made publicly available via a GitHub repository upon publication.

## Notes

### Competing Interest Statement

The authors have declared no competing interest.

### Summary of Updates

This revised version substantially expands and refines the original preprint. We have updated the title, abstract and manuscript framing to more clearly present the study as a multimodal analysis of systemic inflammation and immunity across different states of CFTR dysfunction. Cohort descriptions, sample availability, subgroup definitions and figure references have been corrected and harmonised throughout. We have added extensive supplementary material, including cohort flow and sample availability, CITE-seq quality-control metrics, reference-mapping and cell-type annotation diagnostics, extended PBMC abundance analyses, healthy-control cytokine analyses, expanded monocyte pathway analyses, clinical-cytokine association analyses and additional supporting analyses for predicted intercellular communication. The Results and Discussion have been revised to improve internal consistency, sharpen the interpretation of the healthy carrier findings, and better integrate the longitudinal ETI-associated immune changes with the broader concept of systemic immune variation across CFTR function. The discussion has been substantially shortened in order to preserve the focus of the manuscript. Overall, this revised version provides a more complete, better documented and more internally consistent presentation of the study, with expanded supporting analyses and refined interpretation of the main findings.

### Funding

- Research Foundation - Flanders: 11J2525N (Leander Jonckheere), G024423N (Eva Van Braeckel)
- Fund Alphonse and Jean Forton and Belgian Cystic Fibrosis Association through the King Baudouin Foundation: 365N08923 (Eva Van Braeckel)
- Special Research Fund (BOF) Ghent University: BOF/STA/202109/009 (Eva Van Braeckel)

## REFERENCES

1. Elborn JS. Cystic fibrosis. The Lancet. 2016 Nov 19;388(10059):2519–31. doi:10.1016/S0140-6736(16)00576-6

2. Hanssens LS, Duchateau J, Casimir GJ. CFTR Protein: Not Just a Chloride Channel? Cells. 2021 Oct 22;10(11):2844. doi:10.3390/cells10112844 PubMed PMID: 34831067; PubMed Central PMCID: PMC8616376.

3. Ratner D, Mueller C. Immune Responses in Cystic Fibrosis: Are They Intrinsically Defective? American Journal of Respiratory Cell and Molecular Biology (Online) [Internet]. 2012 Jun [cited 2023 Aug 4];46(6):715–22. Available from: https://www.proquest.com/docview/1020892428/abstract/EB45CA7222ED47E6PQ/1

4. Scambler T, Jarosz-Griffiths HH, Lara-Reyna S, Pathak S, Wong C, Holbrook J, et al. ENaC-mediated sodium influx exacerbates NLRP3-dependent inflammation in cystic fibrosis. eLife. 8:e49248. doi:10.7554/eLife.49248 PubMed PMID: 31532390; PubMed Central PMCID: PMC6764826.

5. Zhang X, Moore CM, Harmacek LD, Domenico J, Rangaraj VR, Ideozu JE, et al. *CFTR*-mediated monocyte/macrophage dysfunction revealed by cystic fibrosis proband-parent comparisons. JCI Insight. 2022 Mar 22;7(6). doi:10.1172/jci.insight.152186 PubMed PMID: 35315363.

6. Sorio C, Buffelli M, Angiari C, Ettorre M, Johansson J, Vezzalini M, et al. Defective CFTR Expression and Function Are Detectable in Blood Monocytes: Development of a New Blood Test for Cystic Fibrosis. PLOS ONE. 2011 Jul 21;6(7):e22212. doi:10.1371/journal.pone.0022212

7. del Fresno C, Gómez-Piña V, Lores V, Soares-Schanoski A, Fernández-Ruiz I, Rojo B, et al. Monocytes from cystic fibrosis patients are locked in an LPS tolerance state: down-regulation of TREM-1 as putative underlying mechanism. PLoS One. 2008 Jul 16;3(7):e2667. doi:10.1371/journal.pone.0002667 PubMed PMID: 18628981; PubMed Central PMCID: PMC2442190.

8. Robledo-Avila FH, Ruiz-Rosado J de D, Brockman KL, Kopp BT, Amer AO, McCoy K, et al. Dysregulated Calcium Homeostasis in Cystic Fibrosis Neutrophils Leads to Deficient Antimicrobial Responses. J Immunol. 2018 Oct 1;201(7):2016–27. doi:10.4049/jimmunol.1800076 PubMed PMID: 30120123; PubMed Central PMCID: PMC6143431.

9. Lara-Reyna S, Scambler T, Holbrook J, Wong C, Jarosz-Griffiths HH, Martinon F, et al. Metabolic Reprograming of Cystic Fibrosis Macrophages via the IRE1α Arm of the Unfolded Protein Response Results in Exacerbated Inflammation. Front Immunol. 2019;10:1789. doi:10.3389/fimmu.2019.01789 PubMed PMID: 31428093; PubMed Central PMCID: PMC6687873.

10. Mueller C, Braag SA, Keeler A, Hodges C, Drumm M, Flotte TR. Lack of Cystic Fibrosis Transmembrane Conductance Regulator in CD3+ Lymphocytes Leads to Aberrant Cytokine Secretion and Hyperinflammatory Adaptive Immune Responses. Am J Respir Cell Mol Biol. 2011 Jun;44(6):922–9. doi:10.1165/rcmb.2010-0224OC PubMed PMID: 20724552; PubMed Central PMCID: PMC3135852.

11. Kushwah R, Gagnon S, Sweezey NB. Intrinsic predisposition of naïve cystic fibrosis T cells to differentiate towards a Th17 phenotype. Respiratory Research. 2013 Dec 17;14(1):138. doi:10.1186/1465-9921-14-138

12. Polverino F, Lu B, Quintero JR, Vargas SO, Patel AS, Owen CA, et al. CFTR regulates B cell activation and lymphoid follicle development. Respiratory Research. 2019 Jul 1;20(1):133. doi:10.1186/s12931-019-1103-1

13. de Torre-Minguela C, Mesa Del Castillo P, Pelegrín P. The NLRP3 and Pyrin Inflammasomes: Implications in the Pathophysiology of Autoinflammatory Diseases. Front Immunol. 2017;8:43. doi:10.3389/fimmu.2017.00043 PubMed PMID: 28191008; PubMed Central PMCID: PMC5271383.

14. Middleton PG, Mall MA, Dřevínek P, Lands LC, McKone EF, Polineni D, et al. Elexacaftor-Tezacaftor-Ivacaftor for Cystic Fibrosis with a Single Phe508del Allele. New England Journal of Medicine. 2019 Nov 7;381(19):1809–19. doi:10.1056/NEJMoa1908639 PubMed PMID: 31697873.

15. Heijerman HGM, McKone EF, Downey DG, Van Braeckel E, Rowe SM, Tullis E, et al. Efficacy and safety of the elexacaftor plus tezacaftor plus ivacaftor combination regimen in people with cystic fibrosis homozygous for the F508del mutation: a double-blind, randomised, phase 3 trial. Lancet. 2019 Nov 23;394(10212):1940–8. doi:10.1016/S0140-6736(19)32597-8 PubMed PMID: 31679946; PubMed Central PMCID: PMC7571408.

16. Sheikh S, Britt Jr. RD, Ryan-Wenger NA, Khan AQ, Lewis BW, Gushue C, et al. Impact of elexacaftor-tezacaftor-ivacaftor on bacterial colonization and inflammatory responses in cystic fibrosis. Pediatric Pulmonology. 2023;58(3):825–33. doi:10.1002/ppul.26261

17. Casey M, Gabillard-Lefort C, McElvaney OF, McElvaney OJ, Carroll T, Heeney RC, et al. Effect of elexacaftor/tezacaftor/ivacaftor on airway and systemic inflammation in cystic fibrosis. Thorax. 2023 Aug 1;78(8):835–9. doi:10.1136/thorax-2022-219943 PubMed PMID: 37208188.

18. De Vuyst RC, Bennard E, Kam CW, McKinzie CJ, Esther CR. Elexacaftor/tezacaftor/ivacaftor treatment reduces airway inflammation in cystic fibrosis. Pediatric Pulmonology. 2023;58(5):1592–4. doi:10.1002/ppul.26334

19. Jarosz-Griffiths HH, Gillgrass L, Caley LR, Spoletini G, Clifton IJ, Etherington C, et al. Anti-inflammatory effects of elexacaftor/tezacaftor/ivacaftor in adults with cystic fibrosis heterozygous for F508del. PLoS One. 2024;19(5):e0304555. doi:10.1371/journal.pone.0304555 PubMed PMID: 38820269; PubMed Central PMCID: PMC11142445.

20. Miller AC, Comellas AP, Hornick DB, Stoltz DA, Cavanaugh JE, Gerke AK, et al. Cystic fibrosis carriers are at increased risk for a wide range of cystic fibrosis-related conditions. Proceedings of the National Academy of Sciences. 2020 Jan 21;117(3):1621–7. doi:10.1073/pnas.1914912117

21. Polgreen PM, Comellas AP. Clinical Phenotypes of Cystic Fibrosis Carriers. Annu Rev Med. 2022 Jan 27;73:563–74. doi:10.1146/annurev-med-042120-020148 PubMed PMID: 35084992; PubMed Central PMCID: PMC8884701.

22. Polgreen PM, Brown GD, Hornick DB, Ahmad F, London B, Stoltz DA, et al. CFTR Heterozygotes Are at Increased Risk of Respiratory Infections: A Population-Based Study. Open Forum Infectious Diseases. 2018 Nov 1;5(11):ofy219. doi:10.1093/ofid/ofy219

23. Shi Z, Wei J, Na R, Resurreccion WK, Zheng SL, Hulick PJ, et al. Cystic fibrosis F508del carriers and cancer risk: Results from the UK Biobank. International Journal of Cancer. 2021;148(7):1658–64. doi:10.1002/ijc.33431

24. Fisman D. Cystic fibrosis heterozygosity: Carrier state or haploinsufficiency? Proc Natl Acad Sci USA. 2020 Feb 11;117(6):2740–2. doi:10.1073/pnas.1921730117

25. Zhang X, Moore CM, Harmacek LD, Domenico J, Rangaraj VR, Ideozu JE, et al. *CFTR*-mediated monocyte/macrophage dysfunction revealed by cystic fibrosis proband-parent comparisons. JCI Insight. 2022 Mar 22;7(6). doi:10.1172/jci.insight.152186 PubMed PMID: 0.

26. Quintelier K, Couckuyt A, Emmaneel A, Aerts J, Saeys Y, Van Gassen S. Analyzing high-dimensional cytometry data using FlowSOM. Nat Protoc. 2021 Aug;16(8):3775–801. doi:10.1038/s41596-021-00550-0

27. Tautz D, Pallares LF, Andersson L, Barghi N, Barton N, Bay R, et al. Beyond Mendel: a call to revisit the genotype-phenotype map through new experimental paradigms. Genetics. 2026 Feb 17;iyag024. doi:10.1093/genetics/iyag024

28. Karczewski KJ, Solomonson M, Chao KR, Goodrich JK, Tiao G, Lu W, et al. Systematic single-variant and gene-based association testing of thousands of phenotypes in 394,841 UK Biobank exomes. Cell Genom. 2022 Sep 14;2(9):100168. doi:10.1016/j.xgen.2022.100168 PubMed PMID: 36778668; PubMed Central PMCID: PMC9903662.

29. Anil N. Mucosal-associated invariant T cells: new players in CF lung disease? Inflamm Res. 2019 Aug 1;68(8):633–8. doi:10.1007/s00011-019-01259-3

30. Germain L, Veloso P, Lantz O, Legoux F. MAIT cells: Conserved watchers on the wall. J Exp Med. 2025 Jan 6;222(1):e20232298. doi:10.1084/jem.20232298 PubMed PMID: 39446132; PubMed Central PMCID: PMC11514058.

31. Smith DJ, Hill GR, Bell SC, Reid DW. Reduced Mucosal Associated Invariant T-Cells Are Associated with Increased Disease Severity and Pseudomonas aeruginosa Infection in Cystic Fibrosis. PLOS ONE. 2014 Oct 8;9(10):e109891. doi:10.1371/journal.pone.0109891

32. Pincikova T, Paquin-Proulx D, Moll M, Flodström-Tullberg M, Hjelte L, Sandberg JK. Severely Impaired Control of Bacterial Infections in a Patient With Cystic Fibrosis Defective in Mucosal-Associated Invariant T Cells. Chest. 2018 May 1;153(5):e93–6. doi:10.1016/j.chest.2018.01.020

33. Bardin E, Dietrich C, Attailia M, Ferroni A, Jamet A, Lezmi G, et al. Restored Cytokine-Producing Capacities of Mucosal-associated Invariant T Cells in Pediatric Cystic Fibrosis Patients Treated with Elexacaftor/Tezacaftor/Ivacaftor. Am J Respir Crit Care Med. 2024 Jul 15;210(2):243–5. doi:10.1164/rccm.202401-0201LE

34. Mulcahy EM, Cooley MA, McGuire H, Asad S, Fazekas de St Groth B, Beggs SA, et al. Widespread alterations in the peripheral blood innate immune cell profile in cystic fibrosis reflect lung pathology. Immunology & Cell Biology. 2019;97(4):416–26. doi:10.1111/imcb.12230

35. Hu Y, Bojanowski CM, Britto CJ, Wellems D, Song K, Scull C, et al. Aberrant immune programming in neutrophils in cystic fibrosis. Journal of Leukocyte Biology. 2024 Mar 1;115(3):420–34. doi:10.1093/jleuko/qiad139

36. Parker D, Cohen TS, Alhede M, Harfenist BS, Martin FJ, Prince A. Induction of type I interferon signaling by Pseudomonas aeruginosa is diminished in cystic fibrosis epithelial cells. Am J Respir Cell Mol Biol. 2012 Jan;46(1):6–13. doi:10.1165/rcmb.2011-0080OC PubMed PMID: 21778412; PubMed Central PMCID: PMC3262660.

37. Romagnani S. T-cell subsets (Th1 versus Th2). Annals of Allergy, Asthma & Immunology. 2000 Jul 1;85(1):9–21. doi:10.1016/S1081-1206(10)62426-X

38. Hu X, Ivashkiv LB. Cross-regulation of Signaling Pathways by Interferon-γ: Implications for Immune Responses and Autoimmune Diseases. Immunity. 2009 Oct 16;31(4):539–50. doi:10.1016/j.immuni.2009.09.002 PubMed PMID: 19833085.

39. Browaeys R, Gilis J, Sang-Aram C, Bleser PD, Hoste L, Tavernier S, et al. MultiNicheNet: a flexible framework for differential cell-cell communication analysis from multi-sample multi-condition single-cell transcriptomics data [Internet]. bioRxiv; 2023 [cited 2024 May 15]. p. 2023.06.13.544751. Available from: https://www.biorxiv.org/content/10.1101/2023.06.13.544751v1 doi:10.1101/2023.06.13.544751

40. Wilson GB, Fudenberg HH, Jahn TL. Studies on Cystic Fibrosis Using Isoelectric Focusing. I. An Assay for Detection of Cystic Fibrosis Homozygotes and Heterozygote Carriers from Serum. Pediatr Res. 1975 Aug;9(8):635–40. doi:10.1203/00006450-197508000-00005

41. Wilson GB, Fudenberg HH. Letter to the Editor: Is Cystic Fibrosis Protein a Diagnostic Marker for Individuals Who Harbor the Defective Gene? Pediatr Res. 1978 Jul;12(7):801–4. doi:10.1203/00006450-197807000-00014

42. Wang S, Song R, Wang Z, Jing Z, Wang S, Ma J. S100A8/A9 in Inflammation. Frontiers in Immunology [Internet]. 2018 [cited 2023 Jan 9];9. Available from: https://www.frontiersin.org/articles/10.3389/fimmu.2018.01298

43. Lin CR, Wei TYW, Tsai HY, Wu YT, Wu PY, Chen ST. Glycosylation-dependent interaction between CD69 and S100A8/S100A9 complex is required for regulatory T-cell differentiation. The FASEB Journal. 2015;29(12):5006–17. doi:10.1096/fj.15-273987

44. Macauley MS, Crocker PR, Paulson JC. Siglec regulation of immune cell function in disease. Nat Rev Immunol. 2014 Oct;14(10):653–66. doi:10.1038/nri3737 PubMed PMID: 25234143; PubMed Central PMCID: PMC4191907.

45. Schroder K, Hertzog PJ, Ravasi T, Hume DA. Interferon-gamma: an overview of signals, mechanisms and functions. J Leukoc Biol. 2004 Feb;75(2):163–89. doi:10.1189/jlb.0603252 PubMed PMID: 14525967.

46. Stewart O, Gruber C, Randolph HE, Patel R, Ramba M, Calzoni E, et al. Monoallelic expression can govern penetrance of inborn errors of immunity. Nature. 2025 Jan;637(8048):1186–97. doi:10.1038/s41586-024-08346-4 PubMed PMID: 39743591; PubMed Central PMCID: PMC11804961.

47. Hao Y, Hao S, Andersen-Nissen E, Mauck WM, Zheng S, Butler A, et al. Integrated analysis of multimodal single-cell data. Cell. 2021 Jun 24;184(13):3573–3587.e29. doi:10.1016/j.cell.2021.04.048 PubMed PMID: 34062119.

