## Supplementary Figures for "Multimodal mapping of systemic inflammation and immunity across the spectrum of CFTR dysfunction"

#### Supplementary Figure 1

##### A. Cohort flow

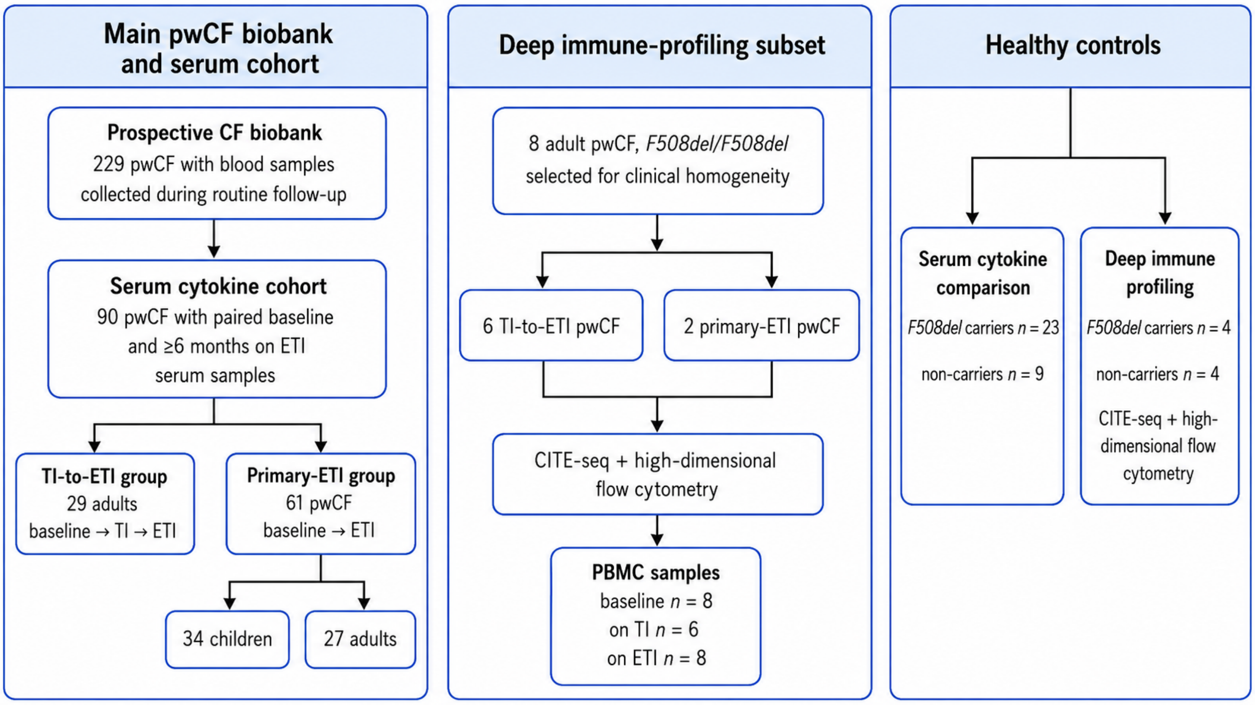


##### B. Sample availability matrix

| Group | Participants | Serum baseline | Serum on TI | Serum on ETI | PBMC baseline | PBMC on TI | PBMC on ETI |
|---|---|---|---|---|---|---|---|
| TI-to-ETI pwCF | 29 adults | 29 | 29 | 29 | 6 | 6 | 6 |
| Primary-ETI pwCF, adults | 27 adults | 27 | — | 27 | 2 | — | 2 |
| Primary-ETI pwCF, children | 34 children | 34 | — | 34 | — | — | — |
| Healthy *F508del* carriers | 23 adults | 23 | — | — | 4 | — | — |
| Non-carriers | 9 adults | 9 | — | — | 4 | — | — |

**Supplementary Figure 1. Cohort flow and sample availability across study modalities. (A)** Cohort flow diagram showing the inclusion of people with cystic fibrosis (pwCF) and healthy controls in the serum cytokine and deep immune-profiling analyses. Blood samples were collected prospectively from 229 pwCF during routine clinical follow-up. The serum cytokine cohort included 90 pwCF with paired samples obtained at baseline and after at least six months of elexacaftor/tezacaftor/ivacaftor (ETI). This cohort comprised 29 adults who transitioned from tezacaftor/ivacaftor (TI) to ETI, with samples available at baseline, on TI and on ETI, and 61 pwCF who initiated ETI as first CFTR modulator therapy, including 34 children and 27 adults, with samples available at baseline and on ETI. The deep immune-profiling subset included eight adult pwCF homozygous for *F508del*, selected for clinical homogeneity, of whom six belonged to the TI-to-ETI group and two to the primary-ETI group. Healthy controls included 23 *F508del* carriers and nine non-carriers for serum cytokine comparison, of whom four carriers and four non-carriers were included in the deep immune-profiling analyses. **(B)** Sample availability matrix showing the number of serum and peripheral blood mononuclear cell (PBMC) samples available across study groups and timepoints. PBMC profiling comprised CITE-seq and high-dimensional flow cytometry. Baseline refers to samples collected before initiation of CFTR modulator therapy. Abbreviations: pwCF, people with cystic fibrosis; TI, tezacaftor/ivacaftor; ETI, elexacaftor/tezacaftor/ivacaftor; PBMC, peripheral blood mononuclear cell.

#### Supplementary Figure 2

##### A. Number of cells retained per sample after QC filtering

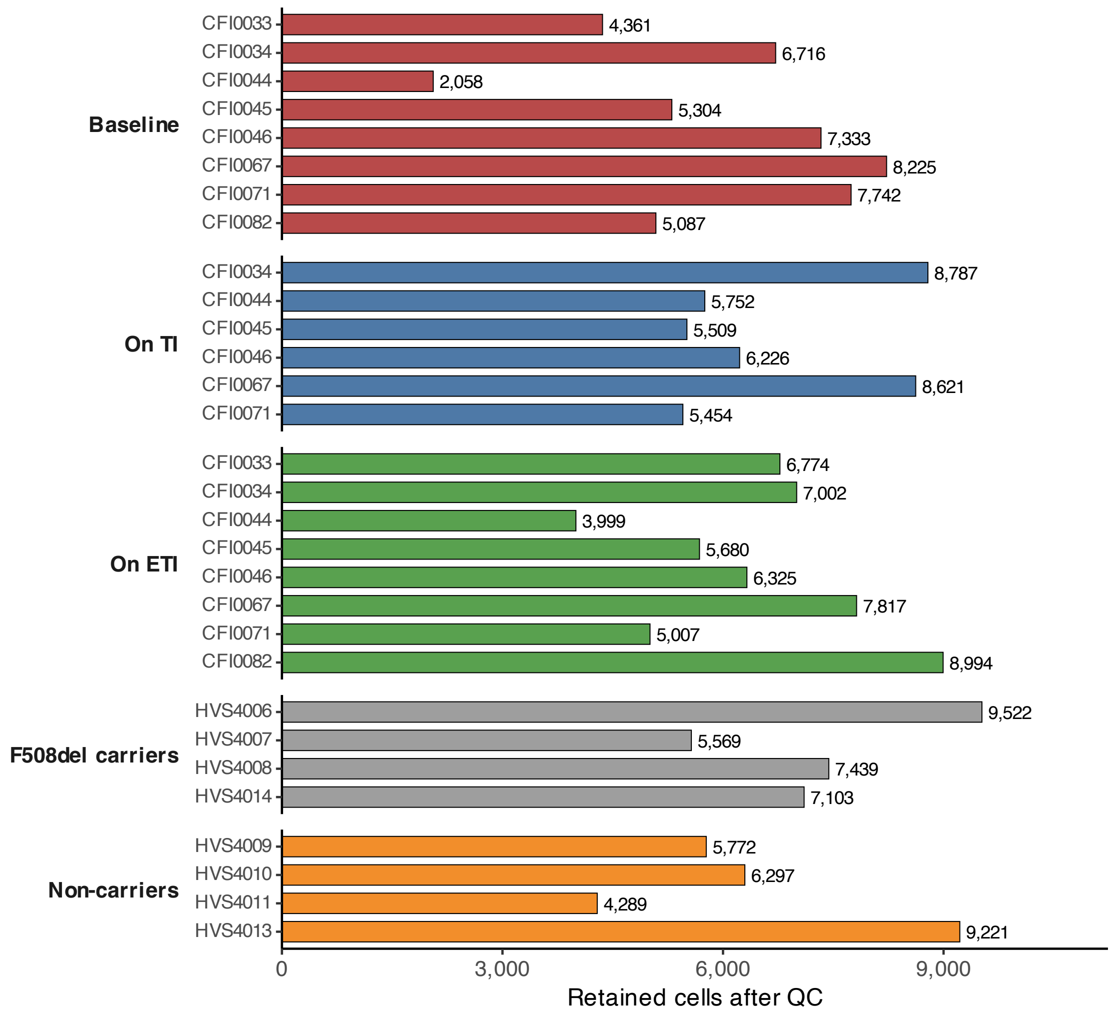


##### B. RNA quality-control metrics across study conditions

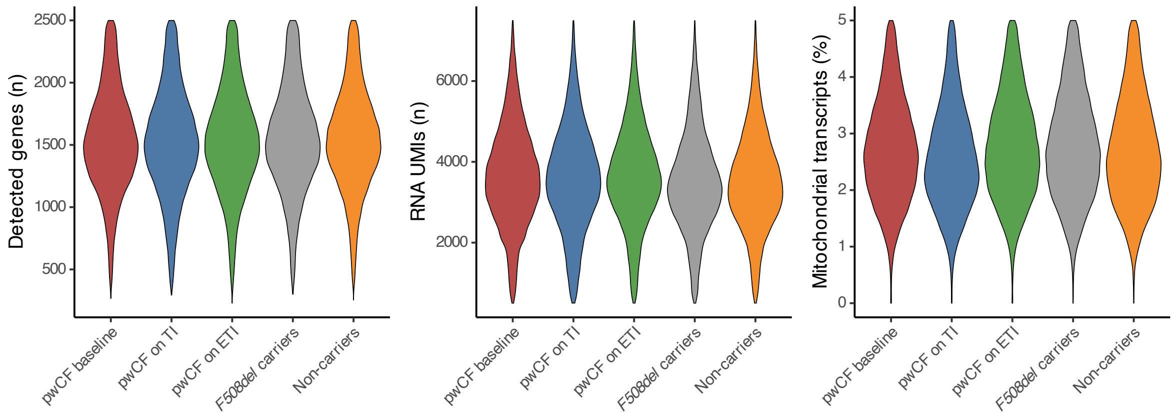

**C. ADT library metrics across study conditions**

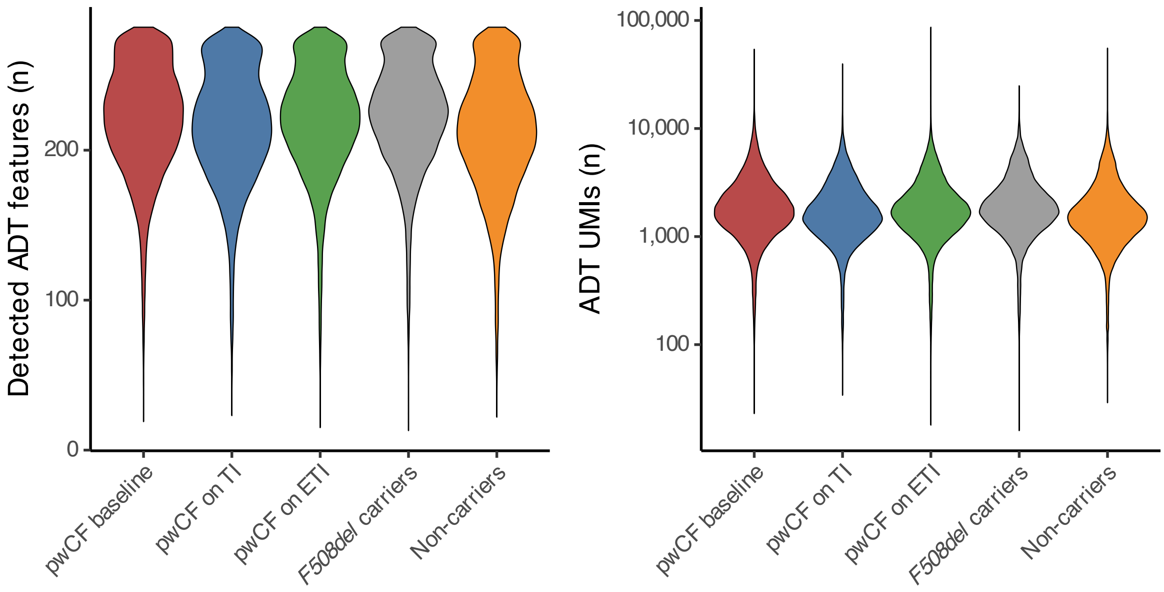


**Supplementary Figure 2. Quality-control metrics for the CITE-seq PBMC atlas. (A)** Number of cells retained after quality-control filtering for each individual sample included in the CITE-seq dataset. Bars are coloured according to study condition. **(B)** Distribution of RNA quality-control metrics across study conditions, including the number of detected genes per cell, total RNA unique molecular identifiers (UMIs) per cell, and percentage of mitochondrial transcripts. Violin plots show single-cell distributions, with embedded boxplots indicating medians and interquartile ranges. **(C)** Distribution of antibody-derived tag (ADT) library metrics across study conditions, including the number of detected ADT features per cell and total ADT UMIs per cell. ADT data were CLR-normalised and integrated jointly with RNA data for downstream weighted-nearest neighbour (WNN) analysis. Abbreviations: ADT, antibody-derived tag; CLR, centred log-ratio; ETI, elexacaftor–tezacaftor–ivacaftor; PBMC, peripheral blood mononuclear cell; TI, tezacaftor–ivacaftor; UMI, unique molecular identifier.

#### Supplementary Figure 3

**A. Weighted-nearest neighbour (WNN) UMAP embedding of the integrated PBMC dataset coloured by study condition.**

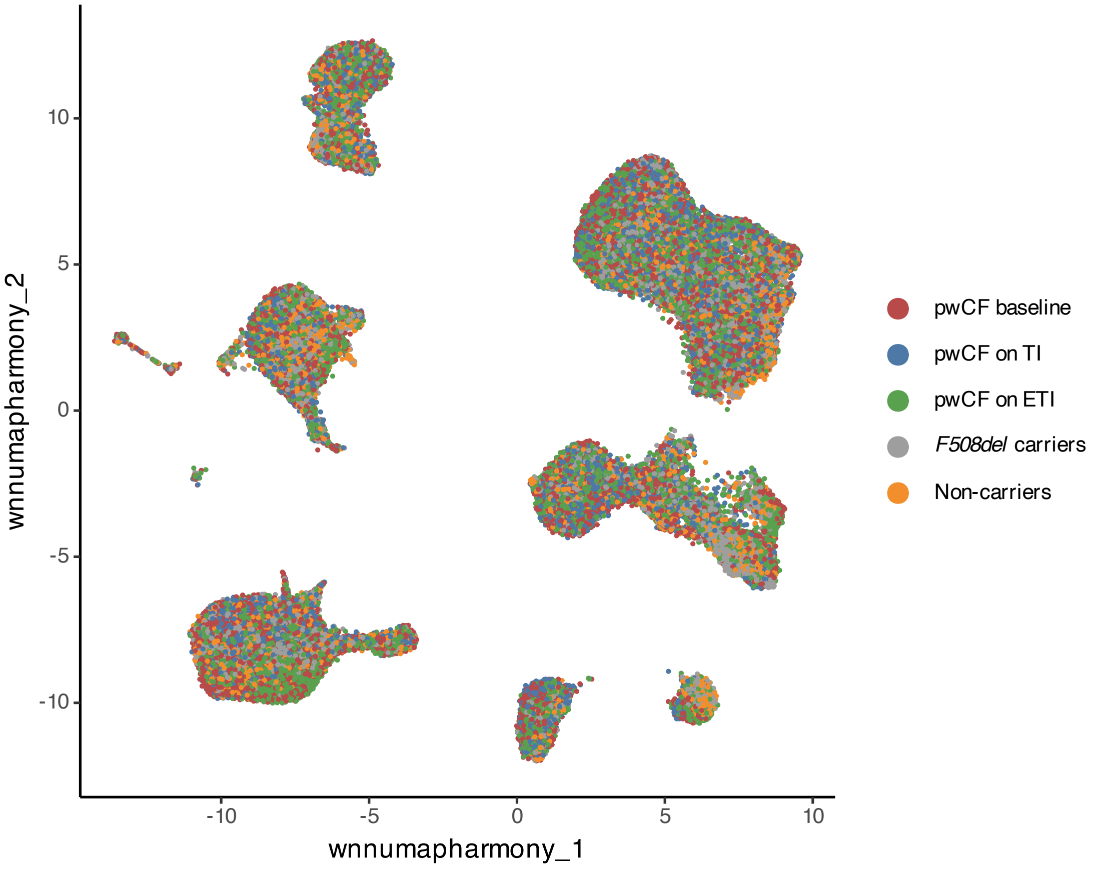

**B. Azimuth reference-mapping prediction scores across annotated cell populations**

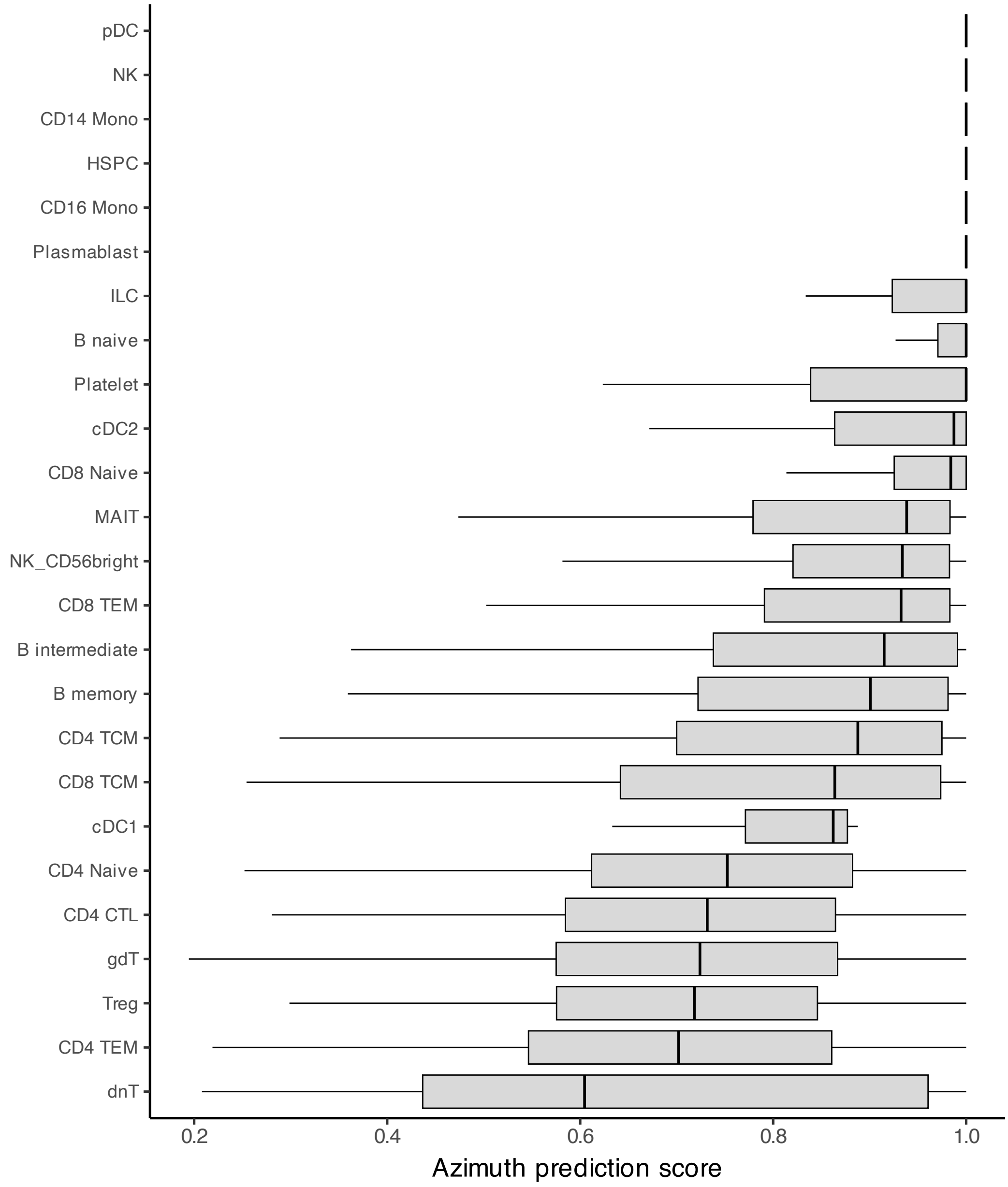

#### C. Expression of canonical Azimuth reference markers across annotated immune-cell populations

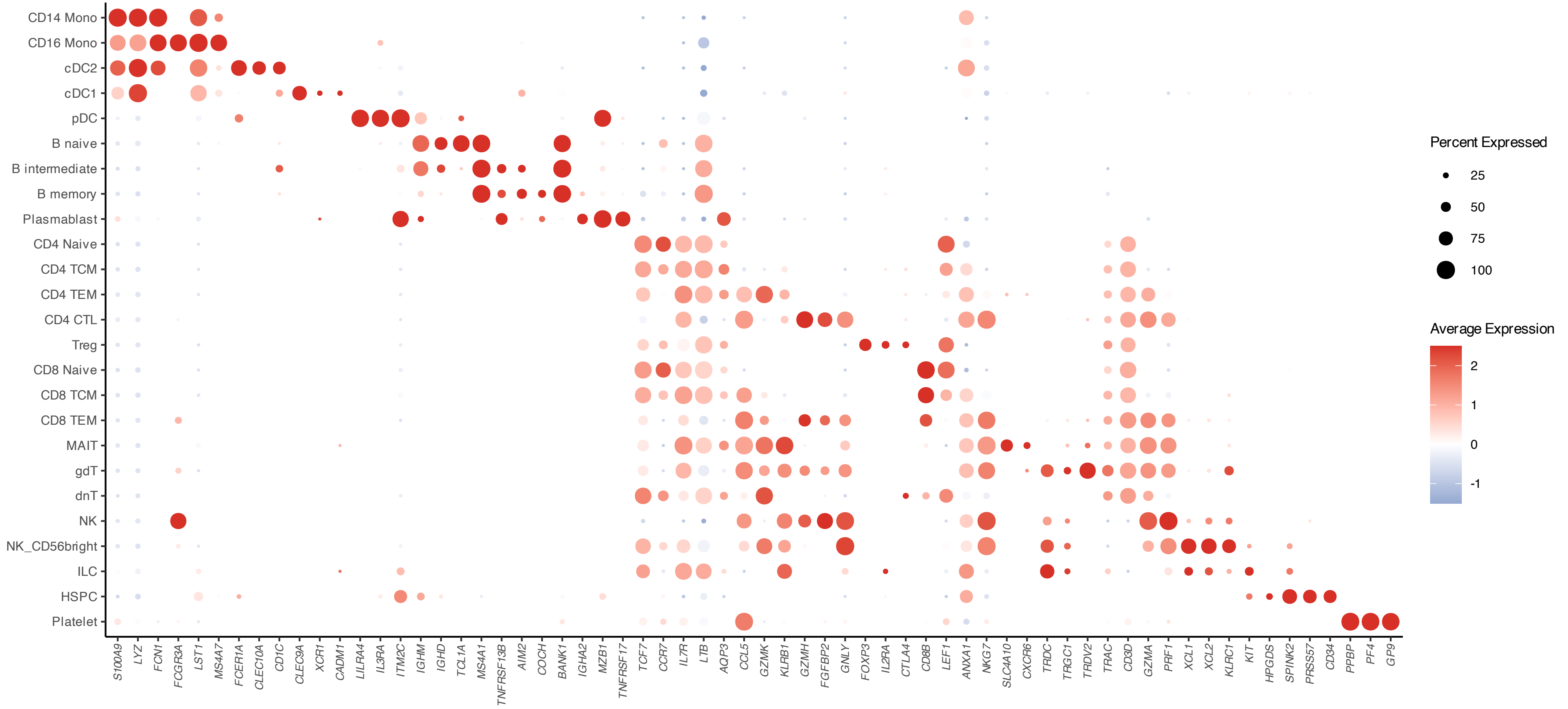

#### D. Expression of canonical ADT markers across annotated immune-cell populations

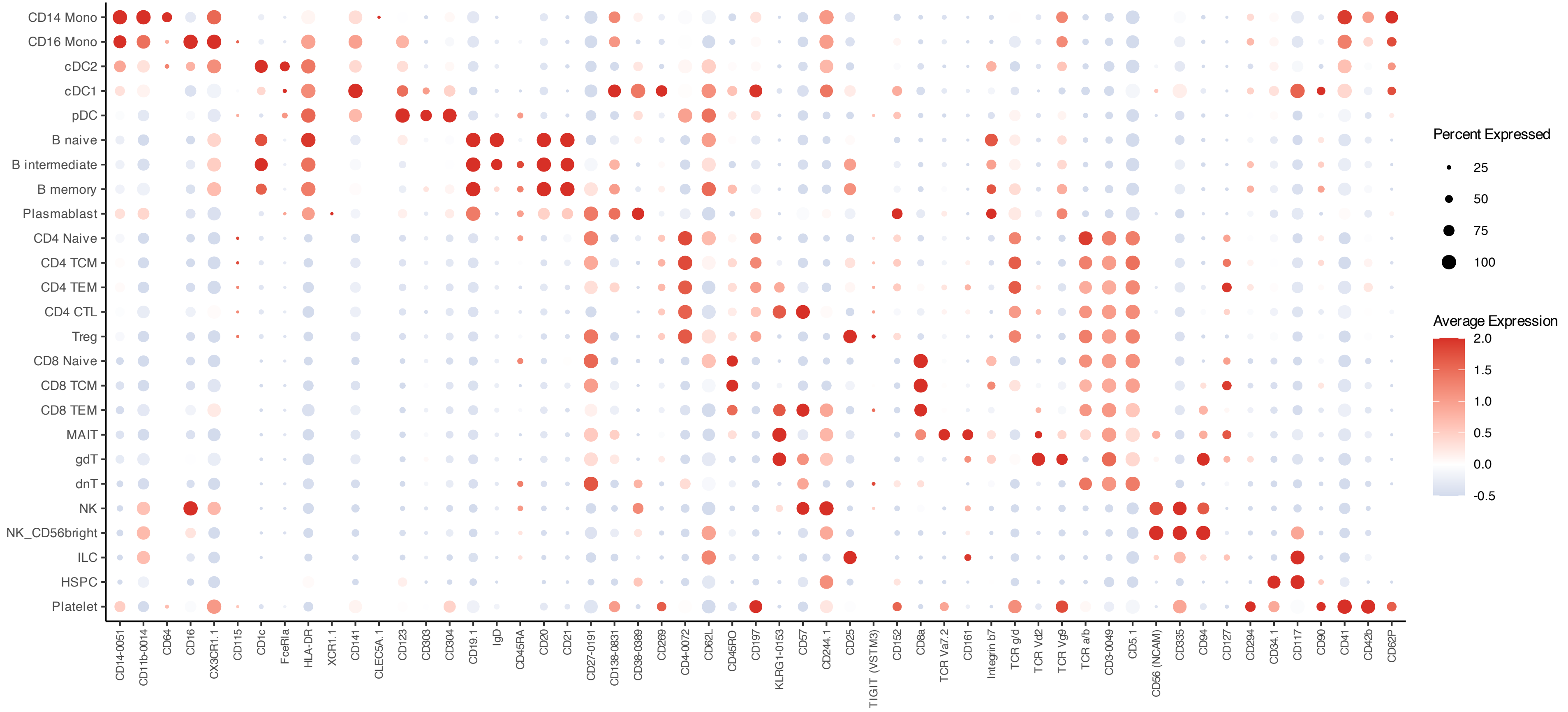

**Supplementary Figure 3. Multimodal reference mapping and annotation of the integrated PBMC CITE-seq dataset. (A)** Weighted-nearest neighbour (WNN) UMAP embedding of the integrated PBMC dataset coloured by study condition. Cells from people with cystic fibrosis (pwCF) at baseline, on tezacaftor/ivacaftor (TI), and on elexacaftor/tezacaftor/ivacaftor (ETI), together with healthy *F508del* carriers and non-carrier controls, are jointly embedded using integrated RNA and antibody-derived tag (ADT) information. **(B)** Distribution of Azimuth prediction scores across final annotated immune-cell populations. Prediction scores reflect the confidence of reference-based cell-type assignment for each cell following mapping to the Human PBMC Azimuth reference atlas. **(C)** Dot plot showing canonical RNA marker expression across annotated immune-cell populations. Markers were selected based on canonical lineage-defining transcripts from the Azimuth Human PBMC reference atlas. Dot size represents the fraction of cells expressing each marker, and colour indicates scaled average expression. **(D)** Dot plot showing canonical ADT marker expression across annotated immune-cell populations. CLR-normalised ADT values were used for visualisation. For visualisation purposes, low CLR-normalised ADT values (< 0.5) were set to zero to reduce background signal. Dot size represents the fraction of cells expressing each marker, and colour indicates scaled average expression. Cell-type annotation was performed using Azimuth reference mapping against the Human PBMC multimodal reference atlas from the HuBMAP consortium, based on the WNN framework described by Hao et al. (Cell 2021). Multimodal integration jointly incorporated RNA and ADT modalities for cell-state definition and annotation.

### Supplementary Figure 4

#### A. Relative abundance of PBMC cell populations across study populations

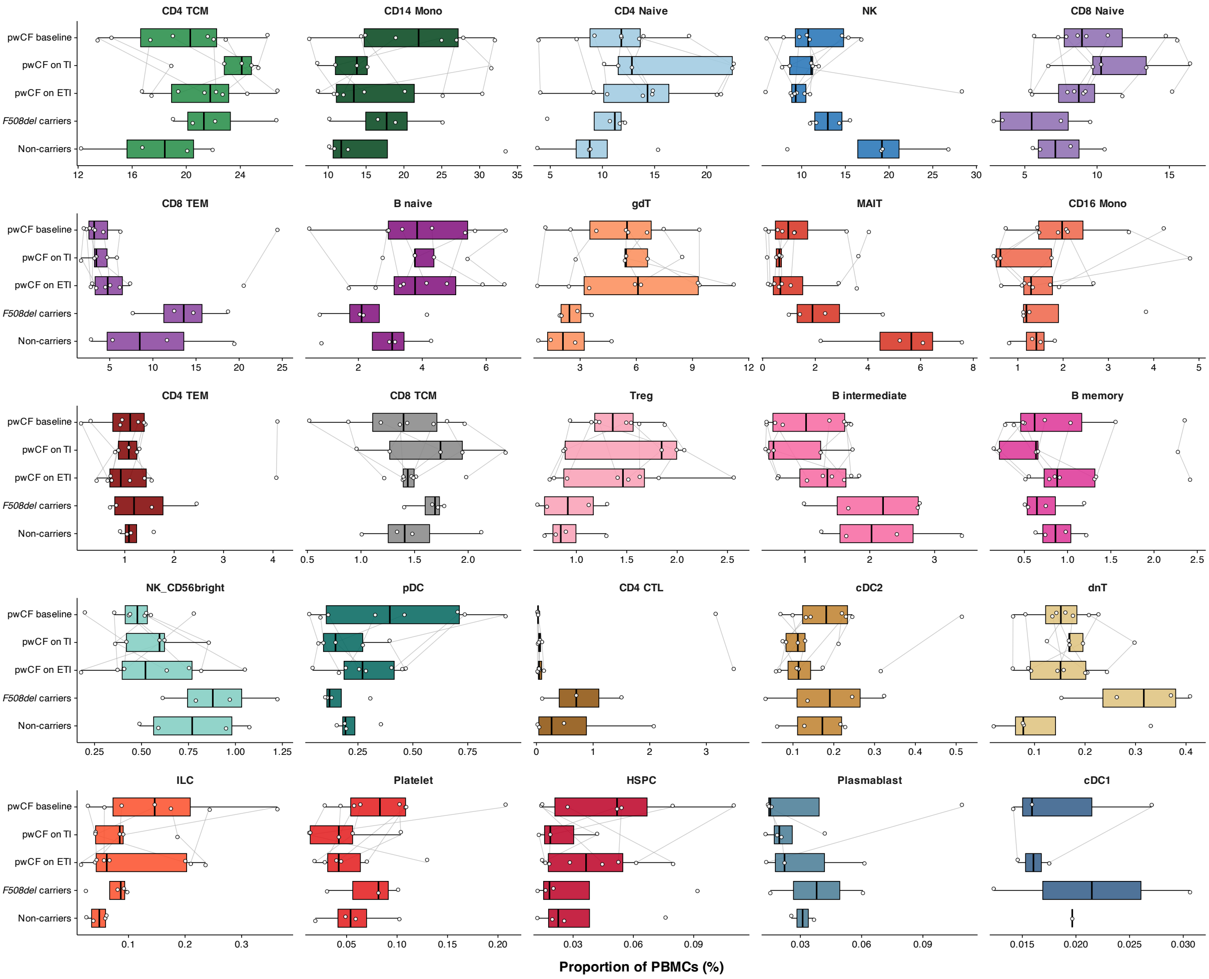

#### B. Pairwise linear mixed-effects estimates for the ten most abundant PBMC populations

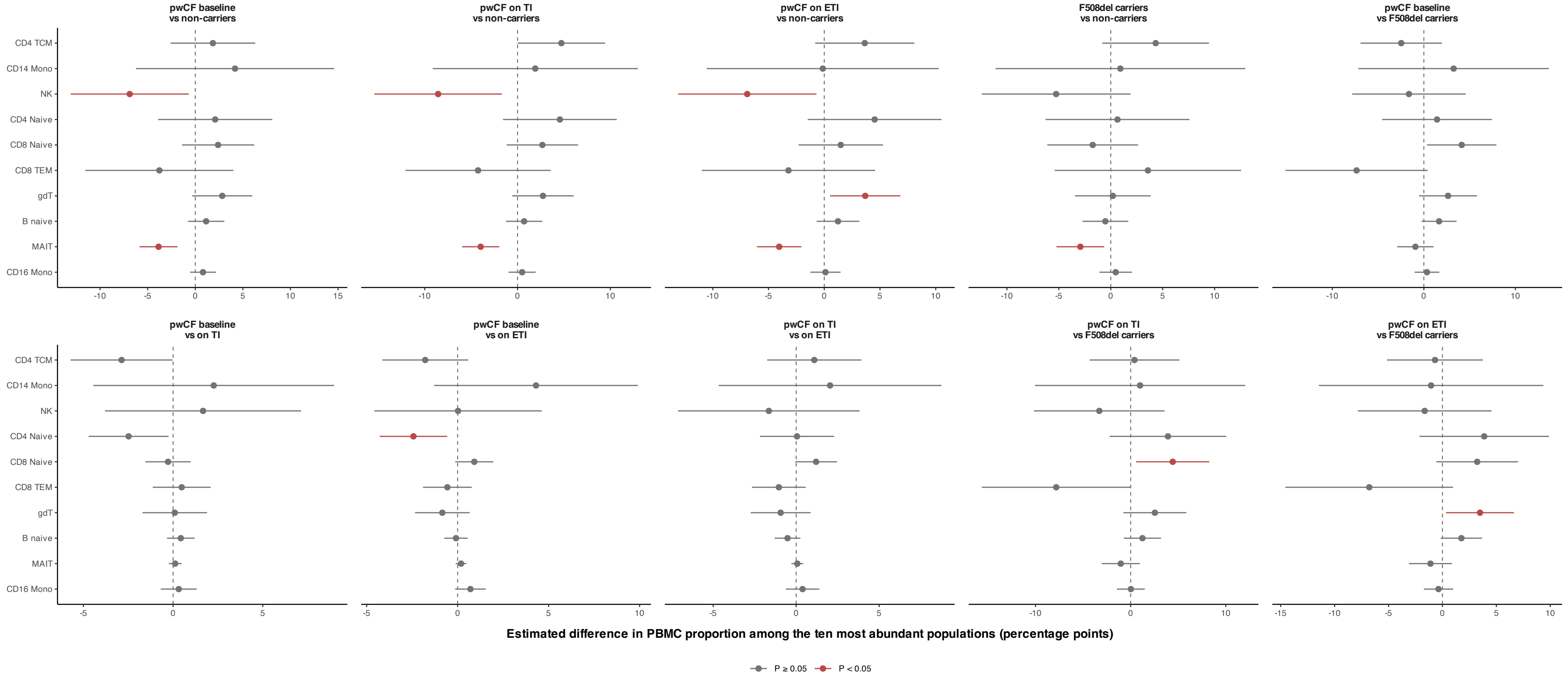

**Supplementary Figure 4. Extended exploratory analyses of PBMC population abundances across study conditions. (A)** Distribution of PBMC population abundances across study groups and treatment conditions. Cell-type proportions were calculated at the sample level and are shown as the percentage of total PBMCs within each sample. Each point represents one individual sample, while boxplots summarise the median and interquartile range across samples. Grey lines connect longitudinal samples obtained from the same pwCF participant across baseline, TI and ETI conditions, illustrating within-subject compositional changes over time. Facets are shown for all annotated PBMC populations and ordered according to overall abundance within the dataset. **(B)** Forest plot summarising pairwise linear mixed-effects model estimates for the ten most abundant PBMC populations. Models were fitted separately for each population using sample-level PBMC proportions as the dependent variable, with condition included as a fixed effect and participant identity as a random intercept to account for repeated longitudinal sampling in pwCF. Estimates are expressed as differences in PBMC proportion (percentage points) between conditions. Points indicate estimated effect sizes and horizontal bars denote 95% confidence intervals. Red points indicate nominal statistical significance ($P < 0.05$). Pairwise comparisons include contrasts between pwCF at baseline, on TI and on ETI, as well as comparisons with healthy *F508del* carriers and non-carrier controls. Abbreviations: ETI, elexacaftor–tezacaftor–ivacaftor; PBMC, peripheral blood mononuclear cell; pwCF, people with cystic fibrosis; TI, tezacaftor–ivacaftor.

#### Supplementary Figure 5

##### A. Representative preprocessing workflow defining the live CD3$^{+}$ T-cell input for FlowSOM analysis

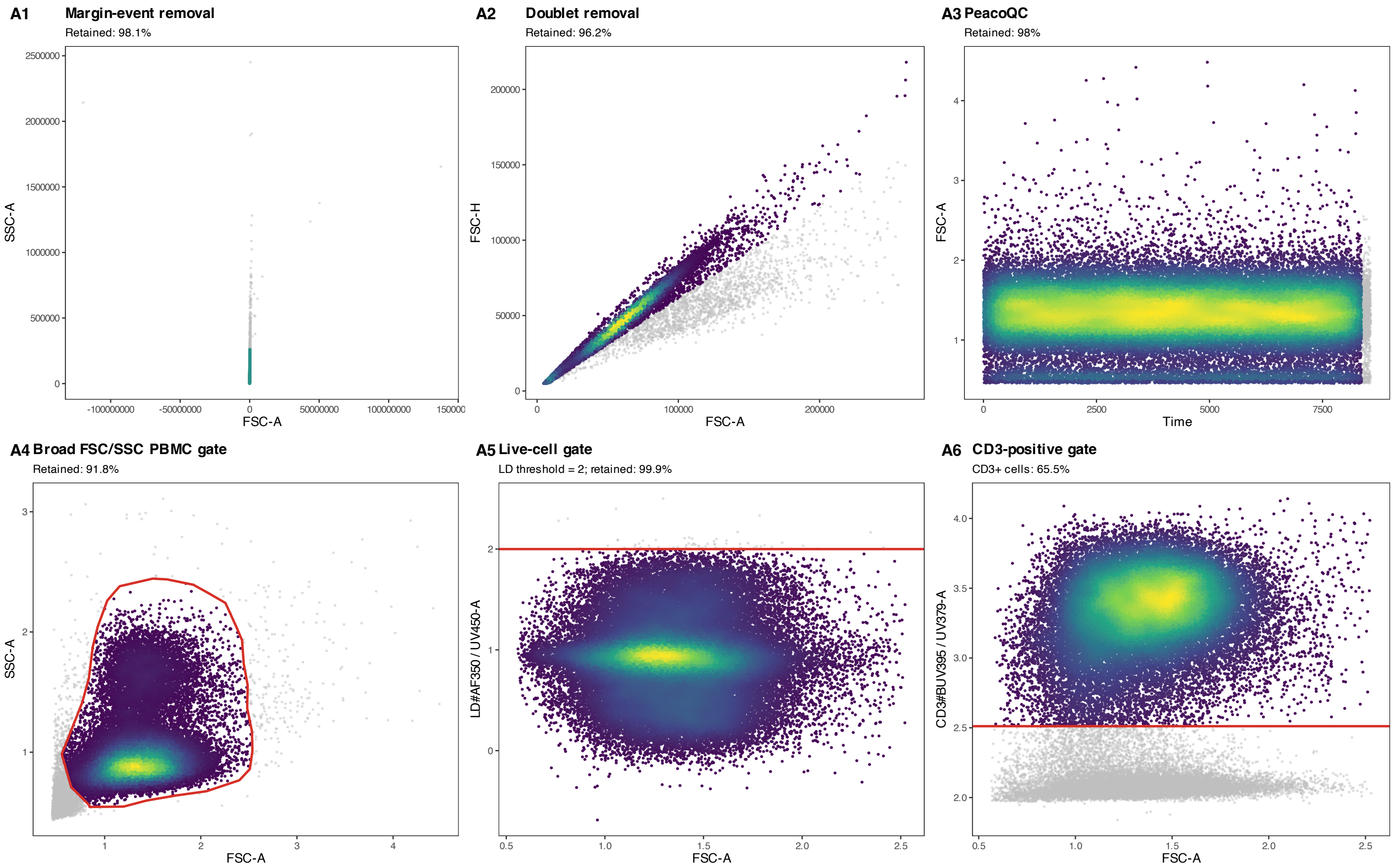

#### B. FlowSOM metacluster tree of CD3$^{+}$ T cells, pooled for healthy controls

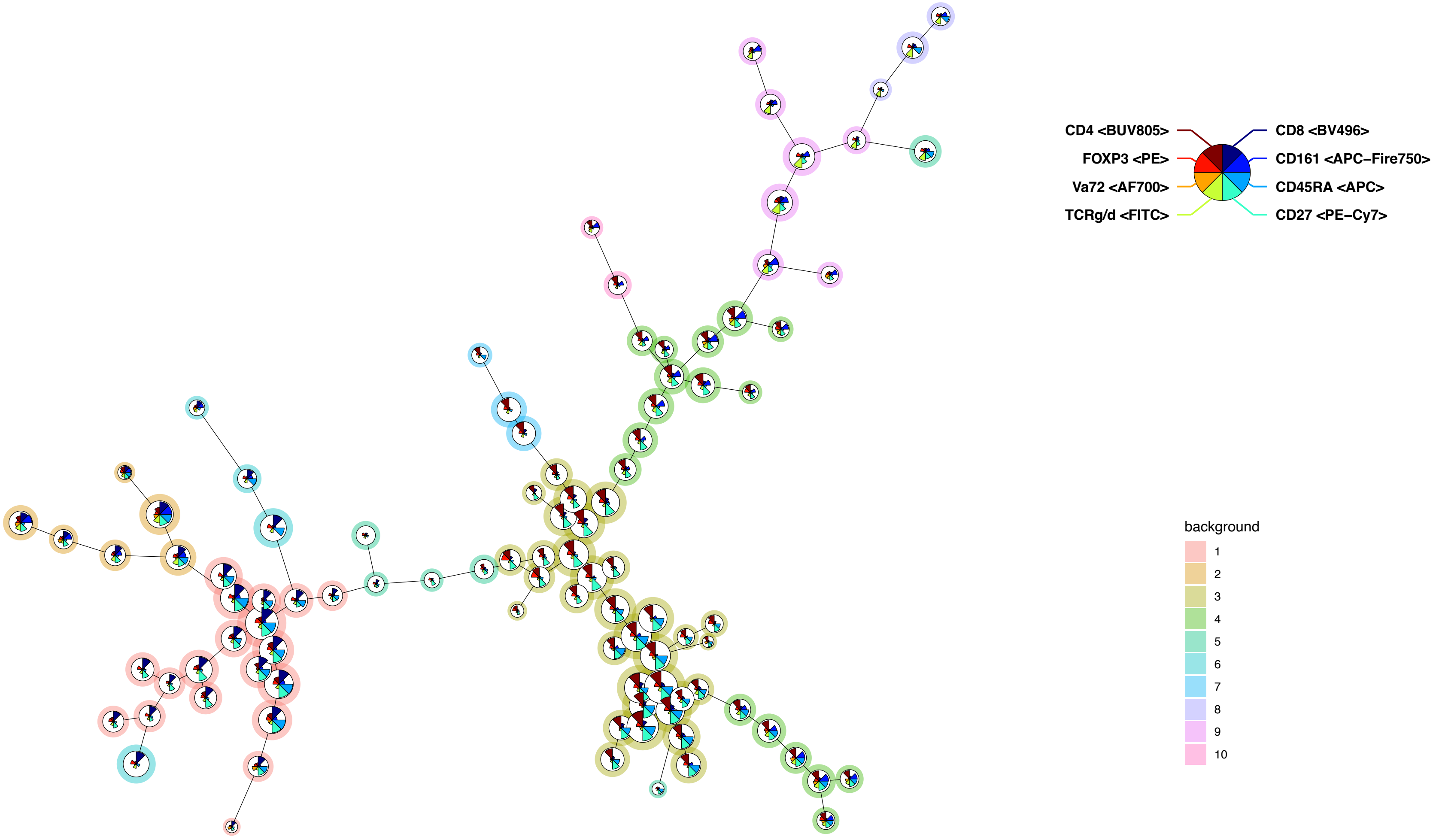

#### C. Median type-marker expression across FlowSOM metaclusters

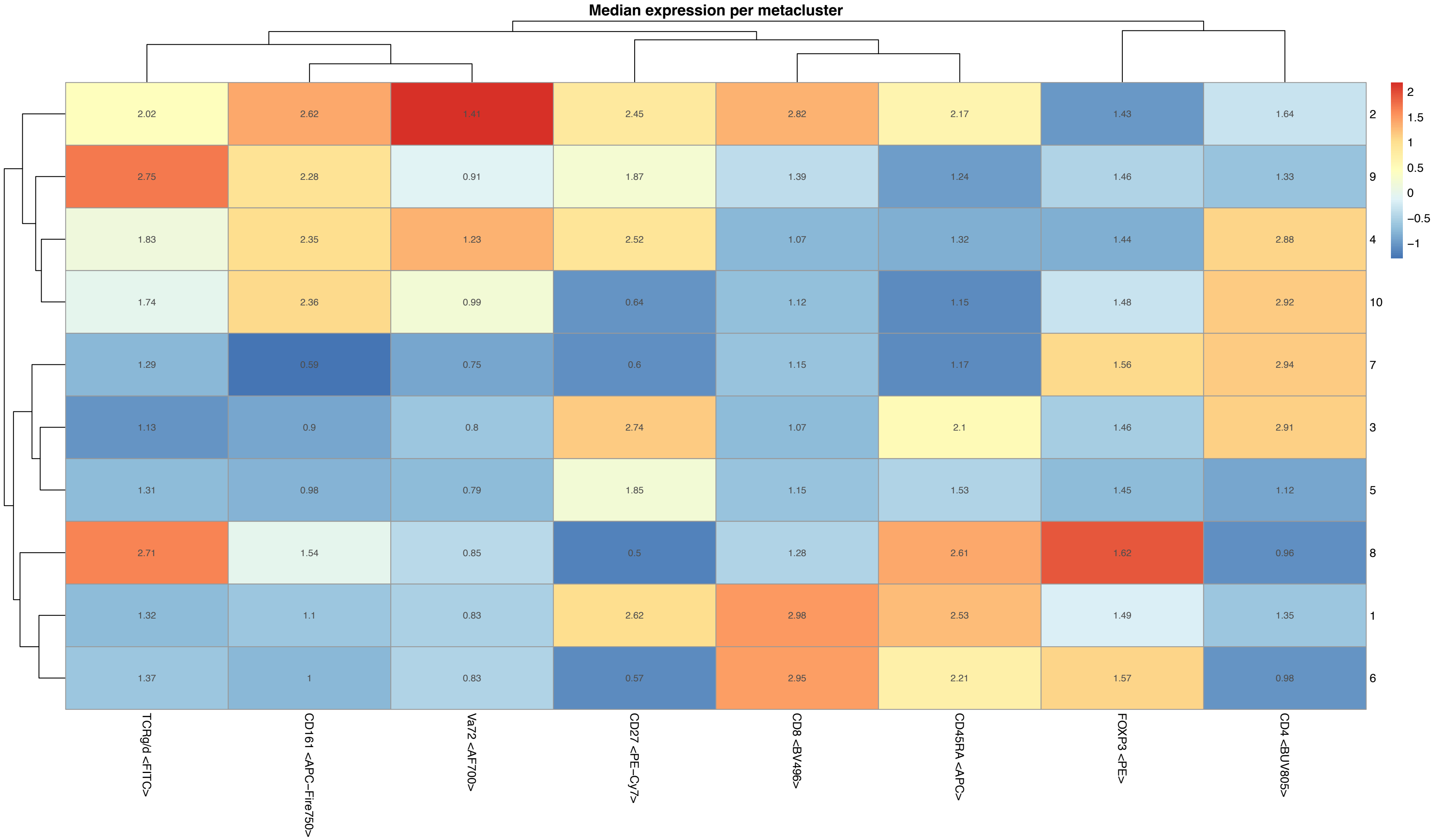

**D. Metacluster abundance in healthy *F508del* carriers and non-carriers**

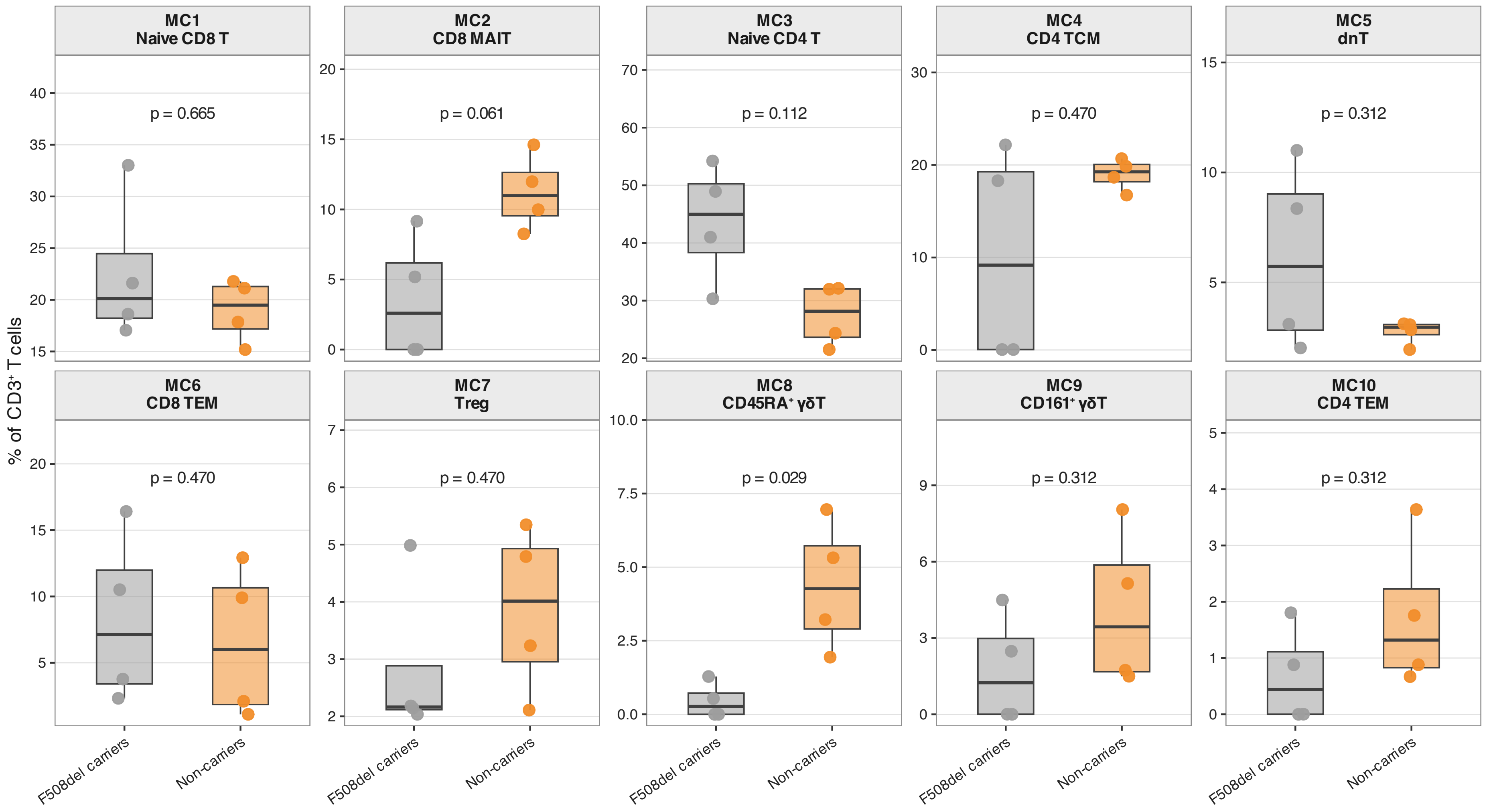

**Supplementary Figure 5. FlowSOM-based annotation and abundance analysis of CD3$^+$ T-cell metaclusters in healthy controls. (A)** Representative preprocessing workflow used to define the input population for downstream FlowSOM analysis. Sequential preprocessing steps are shown for a representative sample, including exclusion of margin events, selection of the PBMC/lymphocyte population, singlet selection, live-cell gating, CD3$^+$ T-cell gating, and final CD3$^+$ T-cell output used for unsupervised clustering. Excluded events are shown in grey. **(B)** FlowSOM metacluster tree generated from healthy-control CD3$^+$ T cells. Each node represents a FlowSOM cluster, and metacluster assignment is indicated by background colour. Star plots visualise relative marker expression patterns across the markers used to construct the model. **(C)** Median expression of type markers across FlowSOM metaclusters. Marker-expression patterns were used to support phenotypic annotation of the ten metaclusters as naïve CD8 T cells, CD8 MAIT cells, naïve CD4 T cells, CD4 central-memory T cells, double-negative T cells, CD8 effector-memory T cells, regulatory T cells, CD45RA$^+$ γδ T cells, CD161$^+$ γδ T cells, and CD4 effector-memory T cells. **(D)** Relative abundance of annotated FlowSOM metaclusters among CD3$^+$ T cells in healthy non-carriers and healthy *F508del* carriers. Dots represent individual samples, and boxplots show the group distribution. Unadjusted Wilcoxon rank-sum p-values are shown for exploratory comparison of metacluster abundance between groups.

#### Supplementary Figure 6

##### A. Serum cytokine profiles in healthy *F508del* carriers and non-carriers

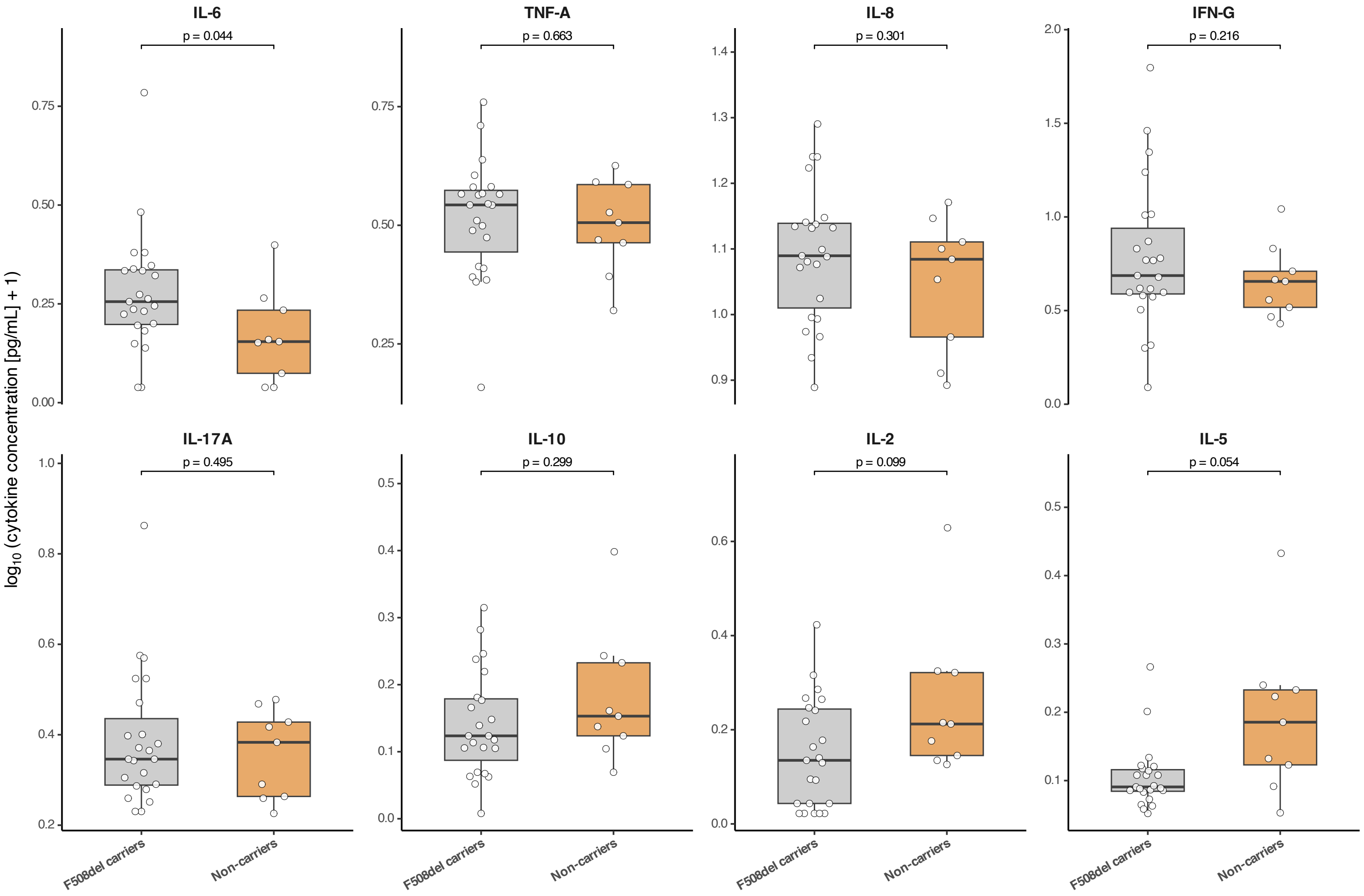

**B. Standardised effect sizes for serum cytokine differences in healthy *F508del* carriers**

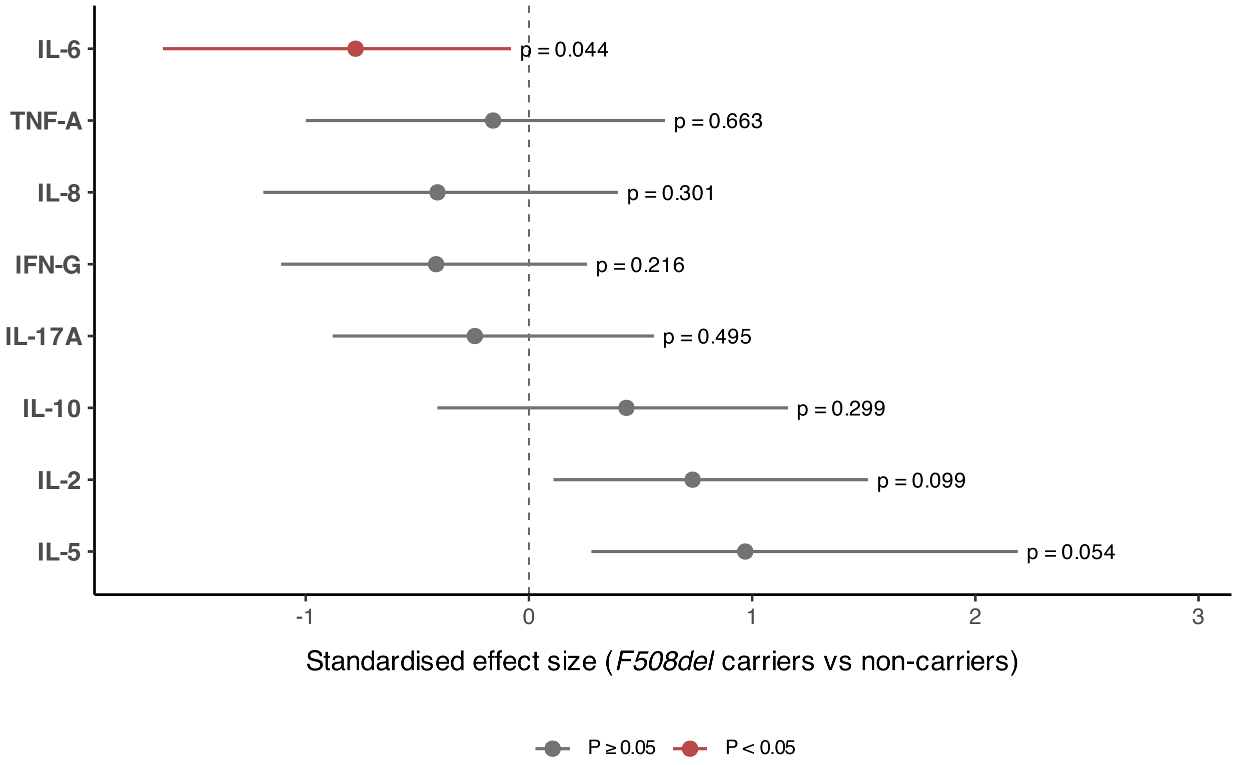


**Supplementary Figure 6. Serum cytokine profiling in healthy *F508del* carriers and non-carriers. (A)** Boxplots showing serum cytokine concentrations in healthy *F508del* carriers and non-carrier controls. Cytokine concentrations were log10-transformed following substitution of values below the detection limit with half of the lower detection limit. Each point represents one individual. Boxes indicate the median and interquartile range (IQR), with whiskers extending to 1.5 × IQR. Statistical comparisons were performed using two-sided Welch's t-tests on log10-transformed concentrations. Exact P values are shown. **(B)** Forest plot summarising standardised effect sizes for serum cytokine differences between healthy *F508del* carriers and non-carrier controls. Points indicate Hedges' g estimates and horizontal lines represent 95% confidence intervals. Positive effect sizes indicate higher cytokine concentrations in *F508del* carriers relative to non-carriers. Cytokines with nominal $P < 0.05$ are highlighted in red.

#### Supplementary Figure 7

##### A. Hallmark GSEA in CD14⁺ monocytes from pwCF at baseline

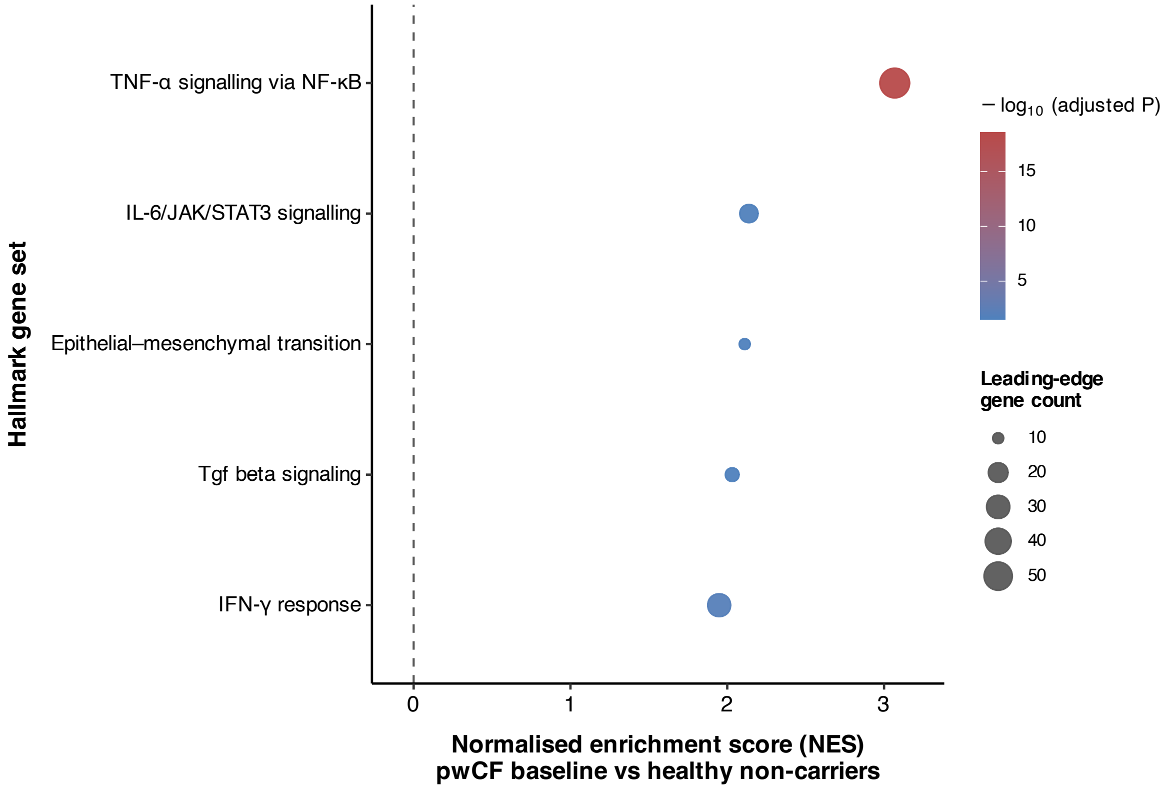


Hallmark GSEA in CD14+ monocytes; positive NES indicates enrichment in pwCF at baseline.

#### B. Gene-level concordance of CD14⁺ monocyte transcriptional changes in pwCF and healthy *F508del* carriers

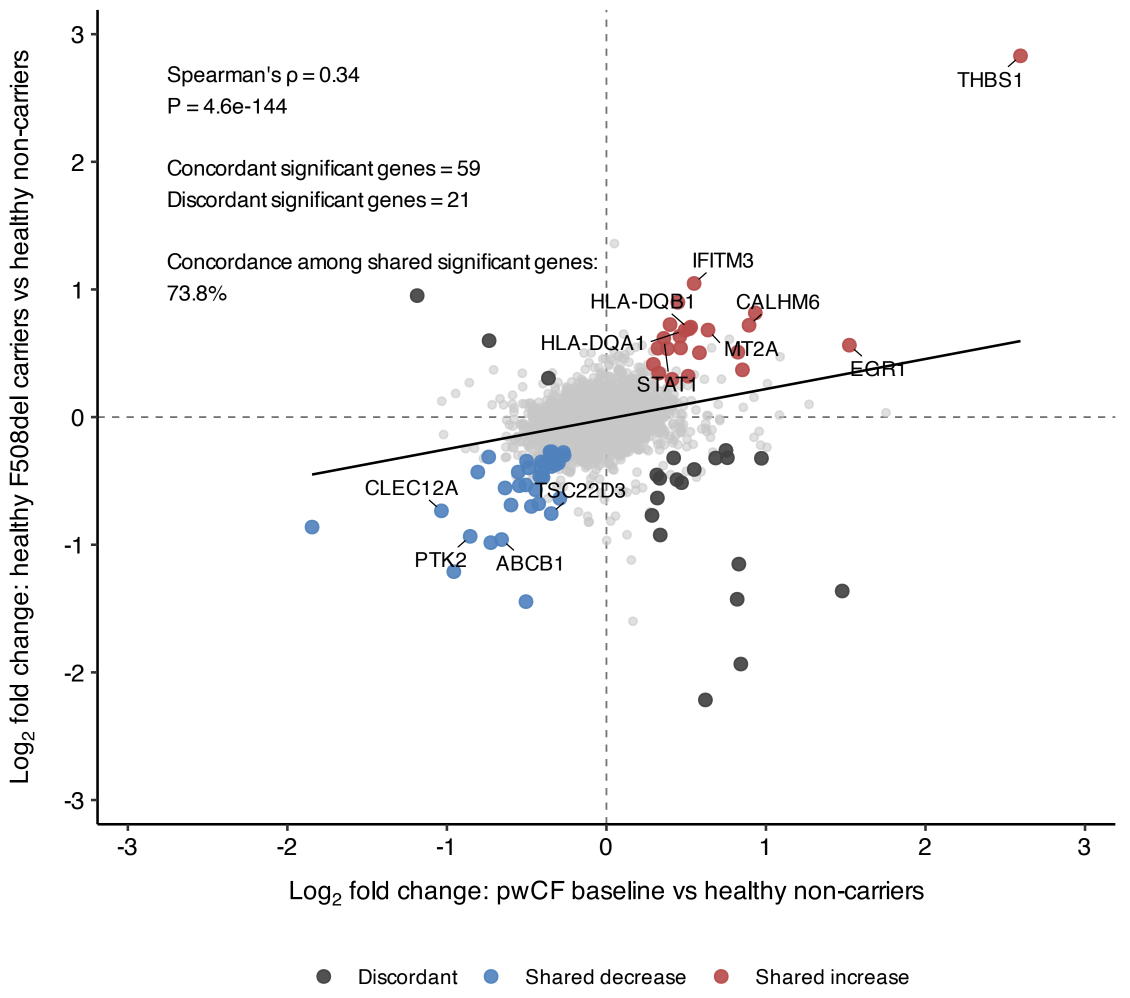


Each point represents one gene tested in both CD14+ monocyte comparisons. Highlighted genes were significant

**C. Pseudobulk expression of representative concordantly altered CD14⁺ monocyte genes**

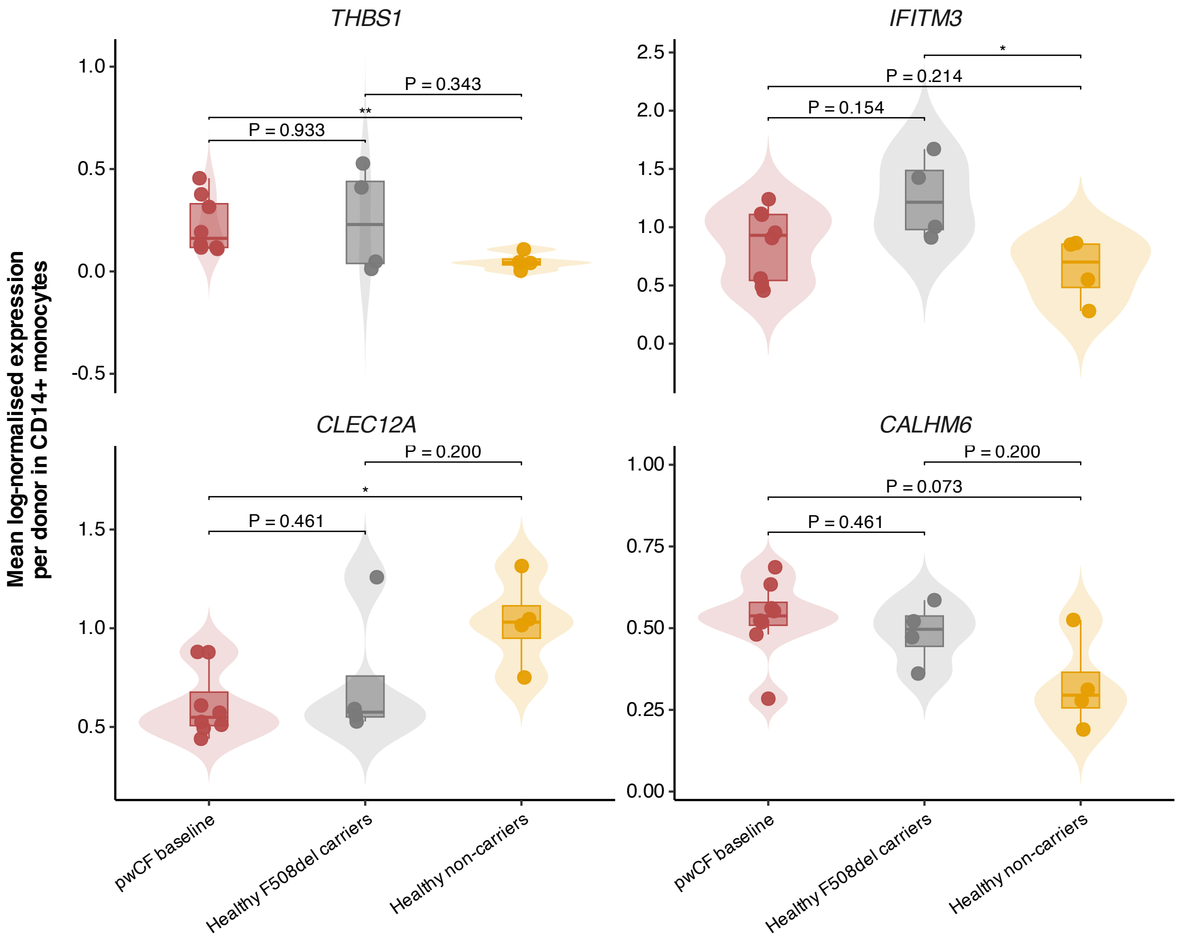


Each point represents one donor. Expression values represent sample-level mean log-normalised RNA expression in CD14+ monocytes.

**Supplementary Figure 7. Shared CD14⁺ monocyte transcriptional alterations in pwCF and healthy *F508del* carriers. (A)** Hallmark gene set enrichment analysis (GSEA) of CD14⁺ monocytes comparing pwCF at baseline (pre-modulator therapy) with healthy non-carriers. Pathways are ranked by normalised enrichment score (NES), with positive NES indicating enrichment in pwCF and negative NES indicating enrichment in healthy non-carriers. Only significantly enriched Hallmark pathways (FDR-adjusted $P < 0.05$) are shown. **(B)** Gene-level concordance analysis comparing differential gene expression in CD14⁺ monocytes from pwCF at baseline versus healthy non-carriers and healthy F508del carriers versus healthy non-carriers. Each point represents a gene significantly differentially expressed in at least one comparison. Genes were classified as shared increase, shared decrease, discordant, or other based on the direction of differential expression across both comparisons. Selected representative genes are labelled. The Spearman correlation coefficient, number of concordant and discordant significant genes, and concordance rate among shared significant genes are indicated. **(C)** Sample-level pseudobulk expression of representative concordantly altered genes in CD14⁺ monocytes. Each point represents one donor, with expression values corresponding to the mean log-normalised RNA expression across all CD14⁺ monocytes from that individual. Genes were selected from the concordant gene set identified in panel B and include *THBS1*,

*IFITM3*, *CLEC12A*, and *CALHM6*. Pairwise comparisons were performed using Wilcoxon rank-sum tests. Asterisks denote significant differences ($P < 0.05$, P < 0.01, $P < 0.001$), whereas exact $P$ values are shown for non-significant comparisons.

#### Supplementary Figure 8

##### A. ETI-associated transcriptional changes in CD14⁺ monocytes of pwCF

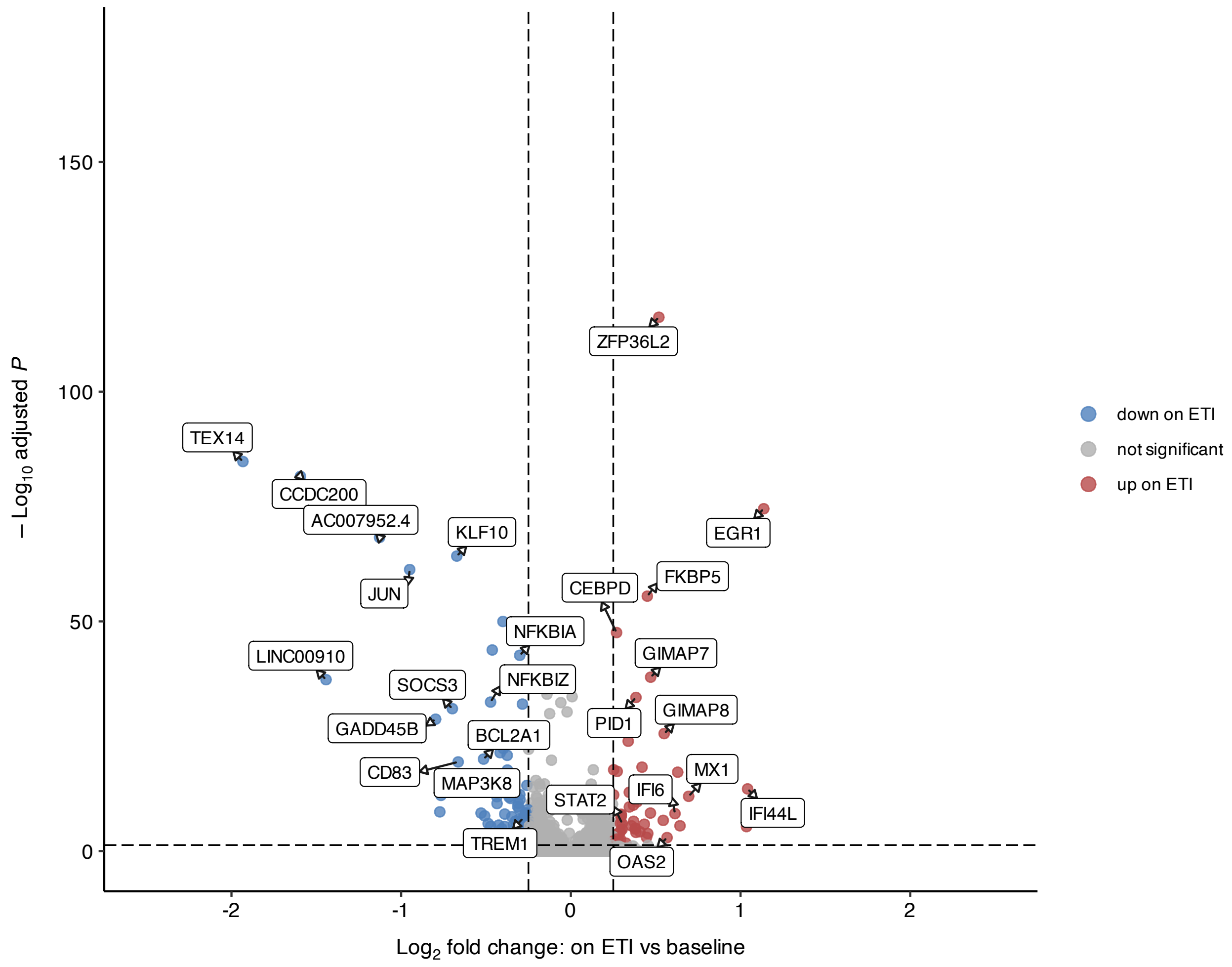

**B. Hallmark gene set enrichment analysis of CD14⁺ monocytes in pwCF on ETI versus baseline**

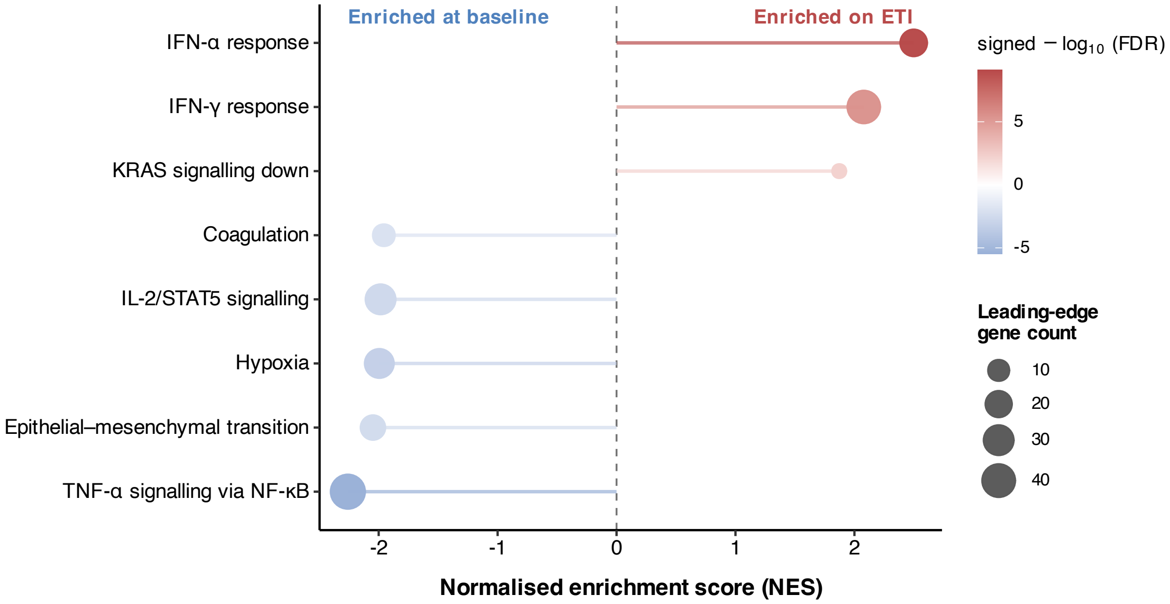


Positive NES indicates enrichment on ETI; negative NES indicates enrichment at baseline.

##### C. Expression of representative patient-level leading-edge genes underlying interferon and TNF/NF-κB pathway changes in CD14$^+$ monocytes from pwCF on ETI versus baseline

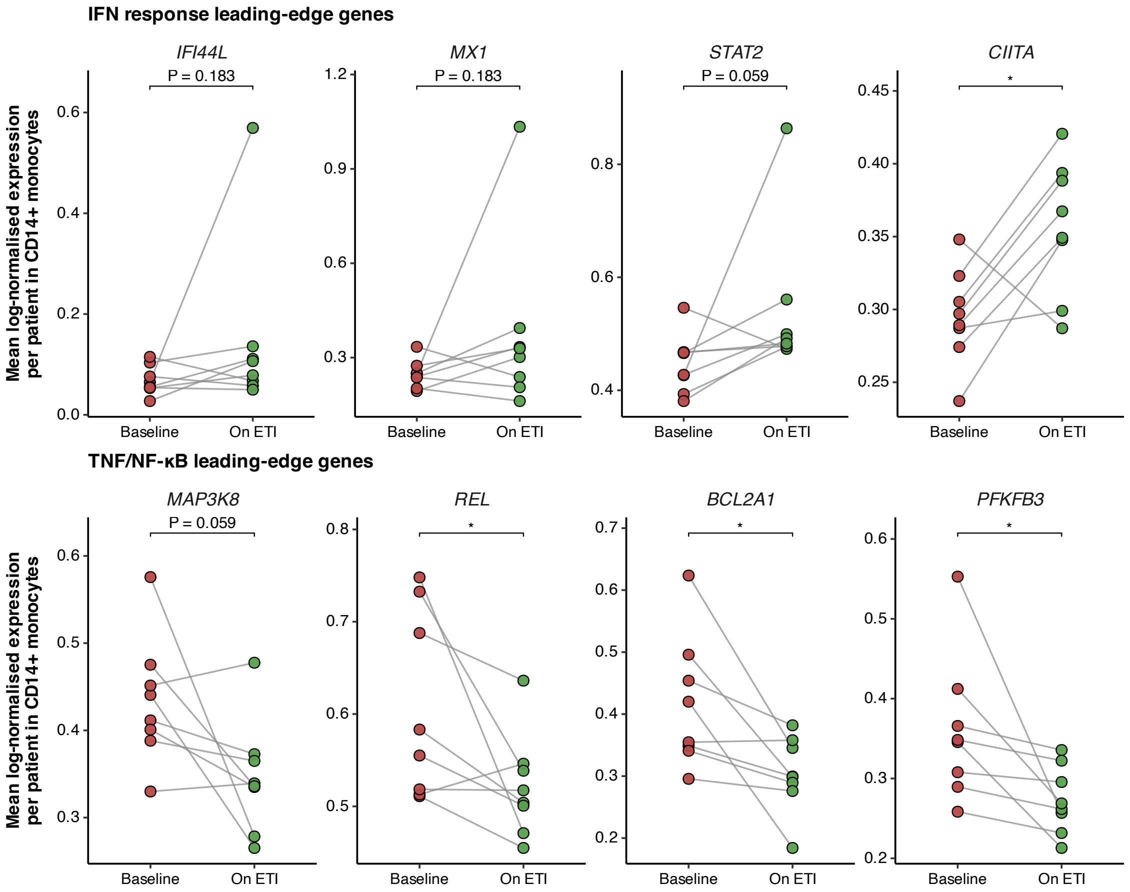


**Supplementary Figure 8. ETI-associated transcriptional changes in CD14$^+$ monocytes from pwCF. (A)** Volcano plot showing differential gene expression in CD14$^+$ monocytes from people with cystic fibrosis (pwCF) on elexacaftor/tezacaftor/ivacaftor (ETI) compared with baseline. Differential expression was assessed using MAST with patient identity included as a latent variable. The x-axis shows the $\log_2$ fold change on ETI versus baseline, and the y-axis shows $-\log_{10}$ adjusted *P*. Red dots indicate genes upregulated on ETI, blue dots indicate genes downregulated on ETI, and grey dots indicate genes not meeting the significance threshold. Selected differentially expressed genes are labelled. **(B)** Hallmark gene set enrichment analysis (GSEA) of CD14$^+$ monocytes from pwCF on ETI compared with baseline, using genes ranked by $\log_2$ fold change from the differential-expression analysis shown in panel A. Positive normalised enrichment scores (NES) indicate enrichment on ETI, whereas negative NES indicate enrichment at baseline. Lollipop length represents the magnitude and direction of enrichment. Dot size indicates the number of leading-edge genes, and colour represents signed $-\log_{10}$ false discovery rate (FDR). Significant ETI-enriched pathways and the top significant baseline-enriched pathways are shown. ETI was associated with enrichment of interferon-related programmes and reduced enrichment of TNF-α/NF-κB-associated inflammatory pathways. **(C)** Patient-level expression of representative leading-edge genes from the interferon response and TNF/NF-κB Hallmark pathways highlighted in panel B. For each gene, expression was summarised per patient as the mean log-normalised expression across $CD14^+$ monocytes at baseline and on ETI. Each connected pair represents one pwCF. The upper row shows representative interferon response leading-edge genes (*IFI44L*, *MX1*, *STAT2*, *CIITA*), whereas the lower row shows representative TNF/NF-κB leading-edge genes (*MAP3K8*, *REL*, *BCL2A1*, *PFKFB3*). Paired comparisons were performed using Wilcoxon signed-rank tests; nominal $P$ values are shown. Gene symbols are italicised.

### Supplementary Figure 9

#### A. Patient-level expression, ligand activity and specificity supporting prioritised CD14$^{+}$ monocyte-derived ligand–receptor interactions at baseline and on ETI in pwCF

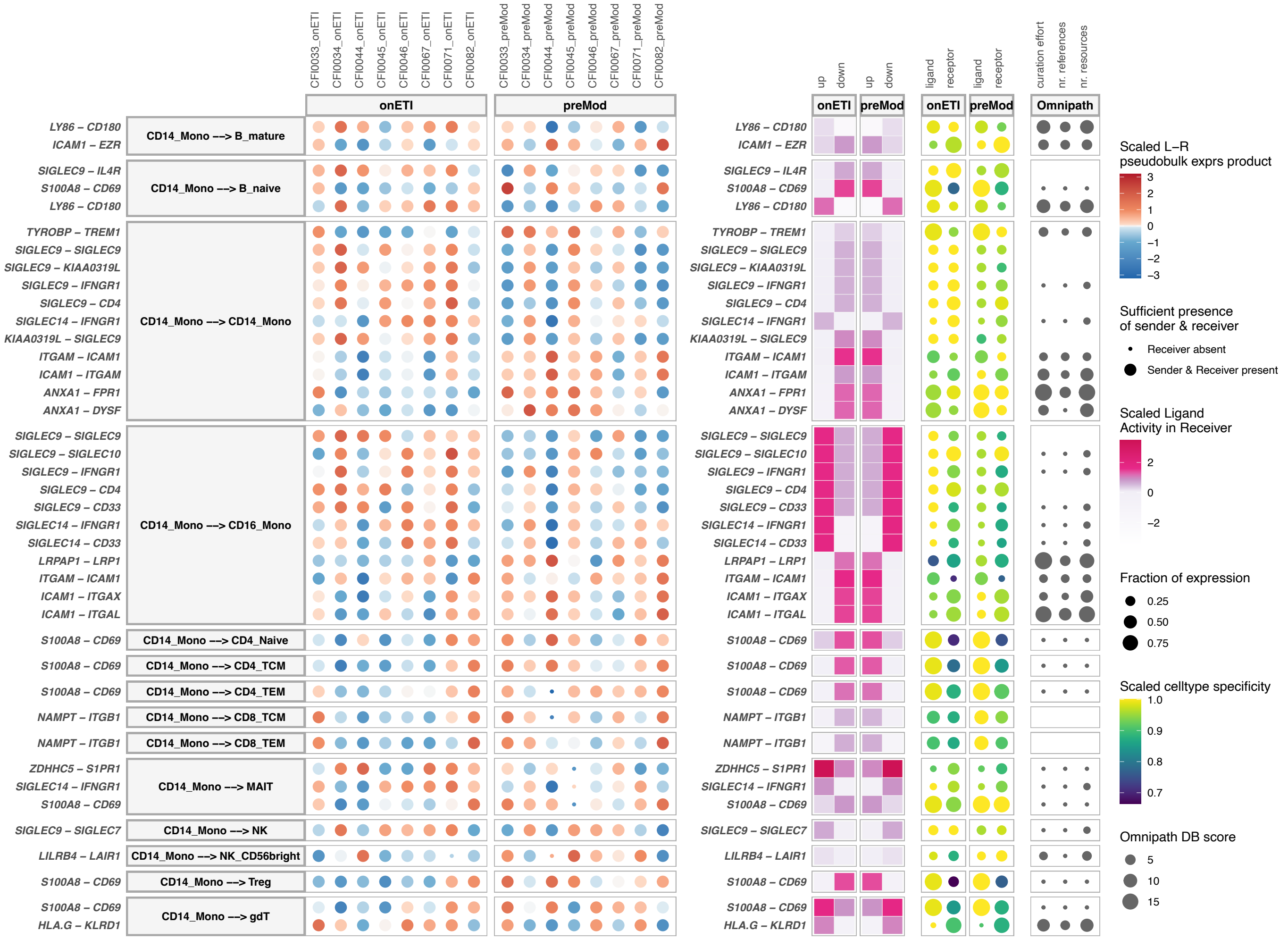

#### B. Predicted ligand activity, regulatory potential and patient-level target-gene expression supporting baseline-prioritised signalling towards MAIT cells in pwCF

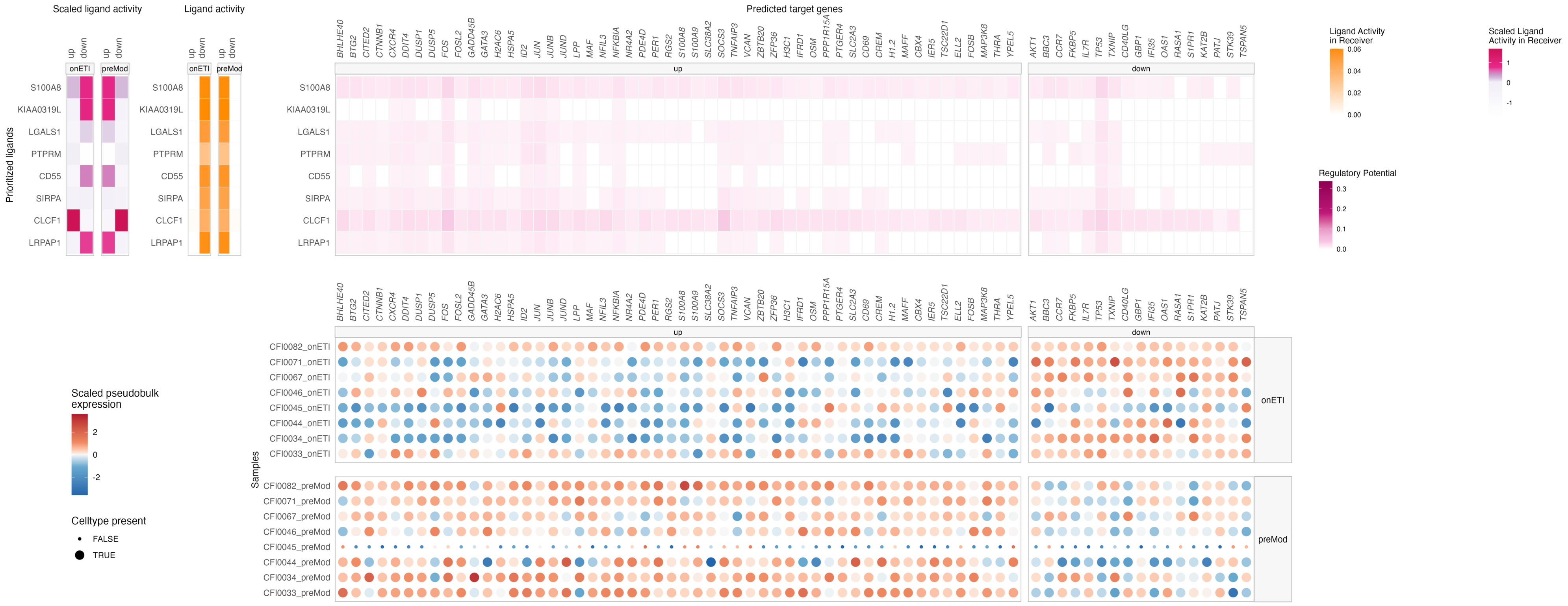

**Supplementary Figure 9. MultiNicheNet output supporting ETI-associated changes in CD14⁺ monocyte-derived predicted intercellular communication in pwCF.** Exploratory ligand-receptor analysis was performed using MultiNicheNet in paired peripheral-blood mononuclear-cell samples from eight pwCF collected at baseline and on elexacaftor–tezacaftor–ivacaftor (ETI). Patient identity was included as a covariate to account for the paired study design. MultiNicheNet integrates sample-level ligand and receptor expression, differential expression, cell-type and condition specificity, expression prevalence, predicted ligand activity in receiver cells and prior ligand–receptor evidence to prioritise condition-associated intercellular interactions. **(A)** Patient-level expression, ligand activity and specificity supporting prioritised $CD14^+$ monocyte-derived ligand–receptor interactions at baseline and on ETI in pwCF. The plot shows the highest-prioritised ligand–receptor interactions originating from $CD14^+$ monocytes across baseline and on-ETI samples. Each row represents a specific $CD14^+$ monocyte sender–receiver cell-type combination and ligand–receptor pair. The individual plot components show the evidence layers contributing to MultiNicheNet prioritisation, including sample-level scaled ligand–receptor pseudobulk expression, predicted ligand activity in the receiver population, ligand and receptor cell-type specificity, the fractions of sender and receiver cells expressing the ligand and receptor, and ligand–receptor database support. At baseline, the most consistent prioritised interaction was S100A8–CD69, with $CD14^+$ monocyte-derived S100A8 directed towards multiple lymphocyte populations, including MAIT cells, Treg, γδ T cells and conventional $CD4^+$ and $CD8^+$ T-cell subsets. This pattern was supported by consistently higher ligand–receptor expression and ligand-activity evidence across baseline samples. On ETI, the most prominent $CD14^+$ monocyte-derived interactions instead involved SIGLEC9–IFNGR1, particularly towards $CD14^+$ and $CD16^+$ monocytes and MAIT cells. **(B)** Predicted ligand activity, regulatory potential and patient-level target-gene expression supporting baseline-prioritised signalling towards MAIT cells in pwCF. Ligand–target analysis was performed for MAIT cells as receiver population at baseline. The left heatmaps show scaled and absolute predicted ligand activity for the prioritised ligands across baseline and on-ETI conditions. The central heatmaps show the predicted regulatory potential of each ligand for genes upregulated or downregulated in baseline MAIT cells, with greater colour intensity indicating stronger predicted ligand–target regulatory potential. The right-hand dot heatmaps show sample-level pseudobulk expression of the corresponding predicted target genes in MAIT cells across individual baseline and on-ETI samples. S100A8 showed the strongest and most coherent baseline-associated ligand-activity signal and was linked to a broad set of predicted target genes that were generally more highly expressed across evaluable baseline MAIT-cell samples. Abbreviations: ETI, elexacaftor–tezacaftor–ivacaftor; LR, ligand–receptor; MAIT, mucosal-associated invariant T; pwCF, people with cystic fibrosis; Treg, regulatory T cells.

#### Supplementary Figure 10

##### A. Exploratory predicted ligand-receptor interactions prioritised in healthy *F508del* carriers and non-carriers

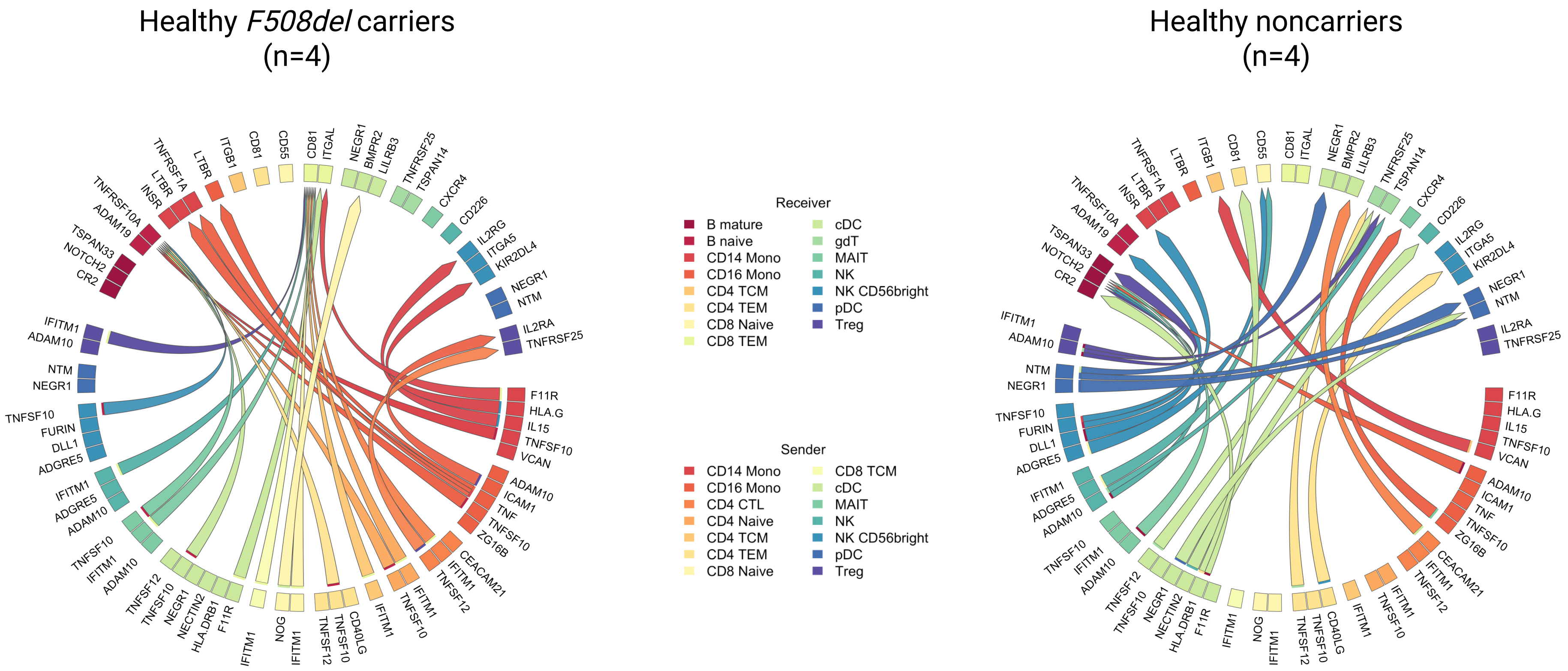

#### B. MultiNicheNet bubble plot of *F508del* carrier-prioritised ligand–receptor interactions originating from CD14⁺ and CD16⁺ monocytes

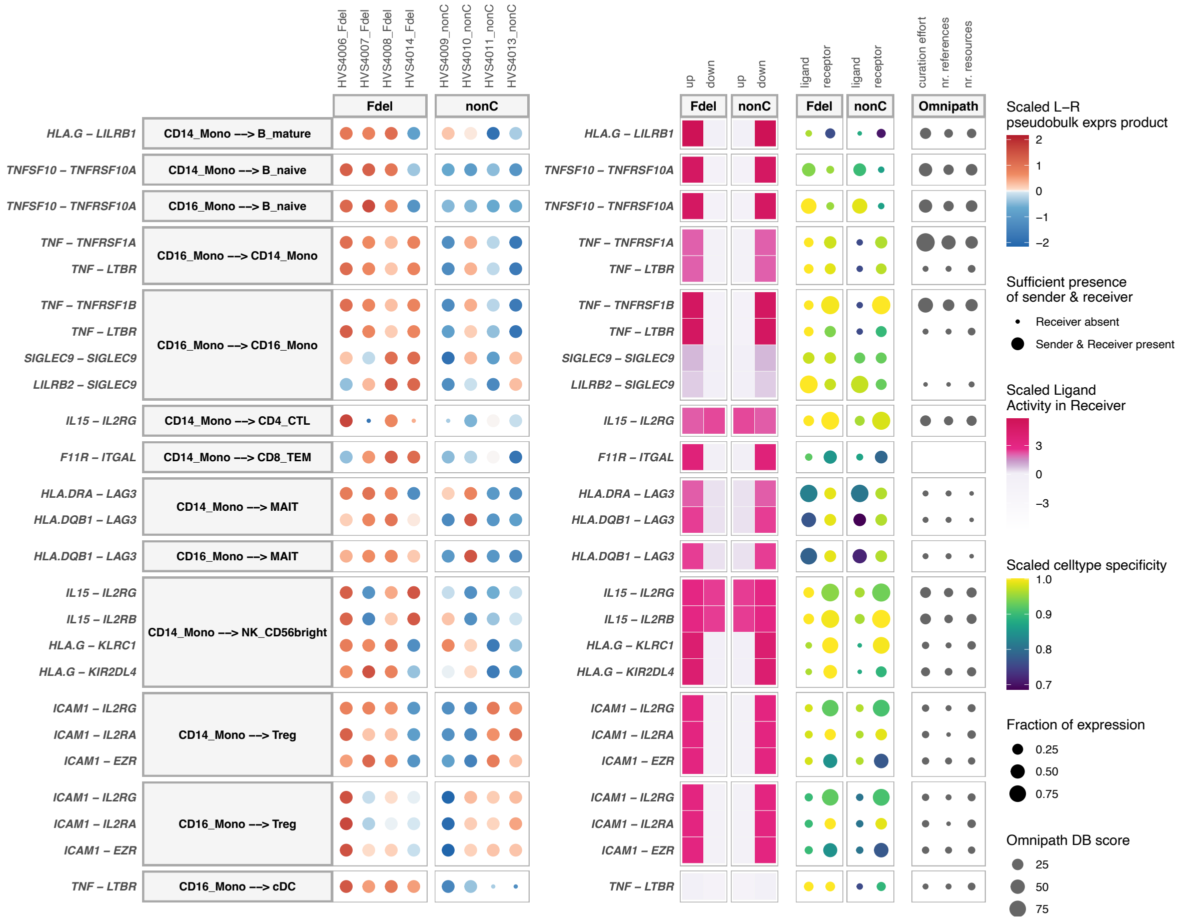

C. **Predicted ligand activity, ligand–target links and target-gene expression in CD14⁺ monocyte receivers from healthy *F508del* carriers versus non-carriers**

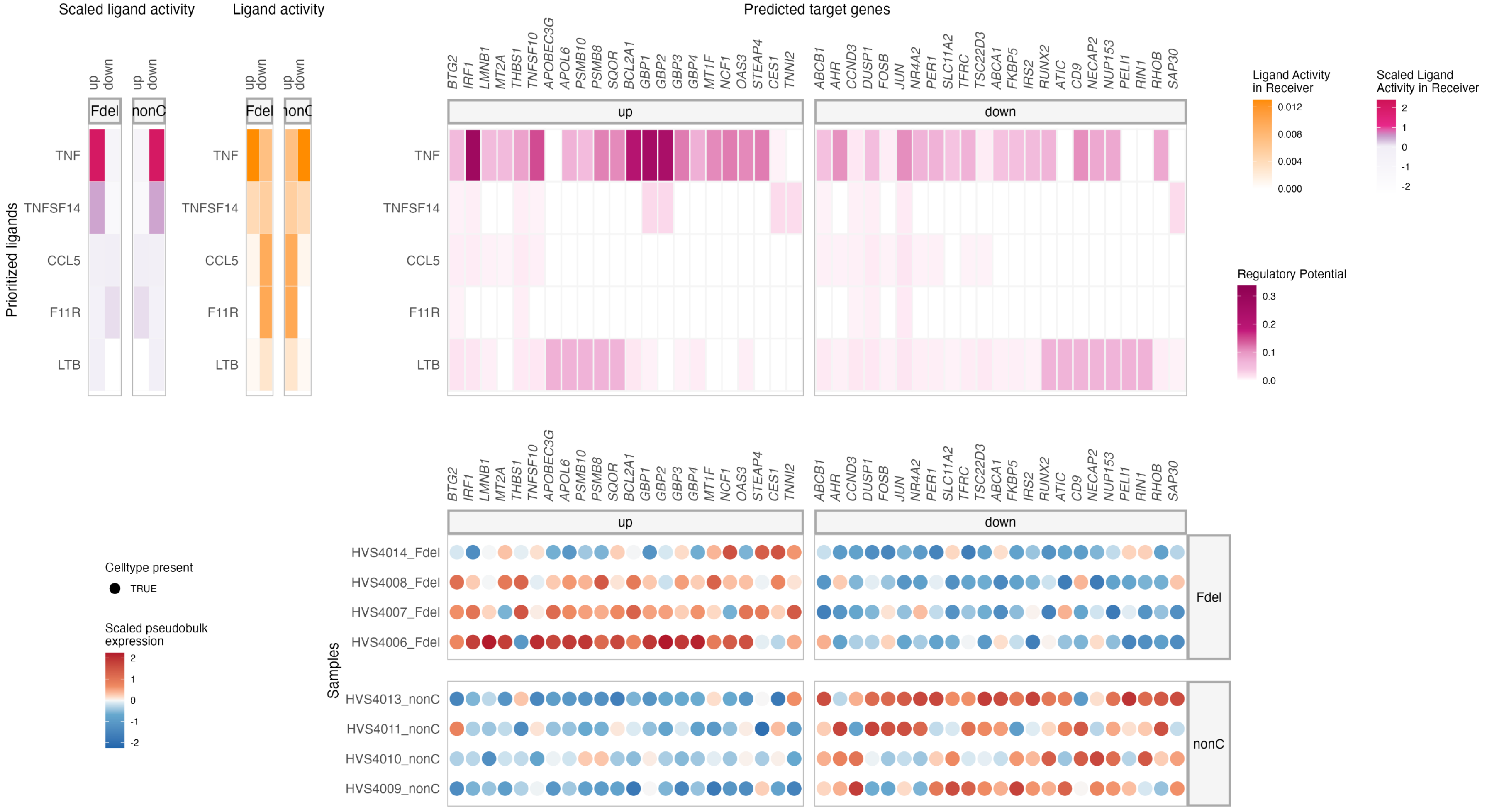

**Supplementary Figure 10. Exploratory MultiNicheNet analysis of predicted intercellular communication in healthy *F508del* carriers and non-carriers.** Exploratory ligand–receptor analysis was performed using MultiNicheNet to compare predicted intercellular communication patterns between healthy *F508del* carriers and healthy non-carrier controls in the CITE-seq PBMC dataset. Because the healthy-control single-cell cohort included four *F508del* carriers and four non-carriers, these analyses were considered hypothesis-generating and were used to complement, rather than independently establish, carrier-associated immune alterations. **(A)** Global overview of prioritised ligand–receptor interactions in healthy *F508del* carriers and non-carriers. Circos plots show the top 25 prioritised ligand–receptor interactions per group, ranked separately for healthy *F508del* carriers and healthy non-carriers. Each sector represents an immune-cell population acting as either a sender or receiver cell type. Curved links represent prioritised ligand–receptor interactions between sender and receiver populations. The colour of each sector corresponds to the cell type, with sender and receiver identities shown in the accompanying legend. These plots provide a global overview of the predicted communication topology in each group. In healthy *F508del* carriers, prioritised interactions included monocyte-centred TNF-superfamily signalling, including $CD14^+$ monocyte-derived TNFSF10–TNFRSF10A interactions towards $CD14^+$ monocytes and $CD16^+$ monocyte-derived TNF interactions towards $CD14^+$ and $CD16^+$ monocytes. Additional carrier-prioritised interactions included $CD14^+$ monocyte-derived IL15–IL2RG and HLA-G–KIR2DL4 interactions towards CD56^bright^ NK cells, ICAM1-associated $CD16^+$ monocyte-to-Treg interactions, and recurrent IFITM1-associated interactions originating from $CD4^+$ T-cell subsets and converging on CD8 TEM cells. In contrast, the non-carrier network showed a different prioritisation pattern, including interactions involving VCAN, ADAM10, ZG16B, CEACAM21, CD40LG, NECTIN2 and NEGR1, with ADAM10-associated signalling converging on NOTCH2. **(B)** MultiNicheNet bubble plot of carrier-prioritised ligand–receptor interactions from $CD14^+$ and $CD16^+$ monocytes. Bubble plots show the data layers supporting the top prioritised ligand–receptor interactions originating from $CD14^+$ and $CD16^+$ monocytes in healthy *F508del* carriers. Each row represents a ligand–receptor pair in a specific sender–receiver context. The left heatmap shows the scaled product of ligand expression in the sender population and receptor expression in the receiver population at the sample level; red indicates relatively higher ligand–receptor pseudobulk expression product and blue indicates lower expression product. The adjacent ligand-activity heatmap shows the scaled predicted activity of each ligand in the receiver population in *F508del* carriers and non-carriers. Dot plots show cell-type specificity of ligand and receptor expression, with dot size indicating the fraction of cells expressing the ligand or receptor and colour indicating scaled specificity. The rightmost OmniPath evidence layer indicates the level of database support for the corresponding ligand–receptor interaction. The most consistent carrier-prioritised signal involved $CD16^+$ monocyte-derived TNF interactions towards $CD14^+$ and $CD16^+$ monocytes, including TNF–TNFRSF1A and TNF–TNFRSF1B. These interactions showed higher ligand–receptor pseudobulk expression across *F508del* carrier samples, increased predicted ligand activity in the receiver population, high cell-type specificity, substantial expression fractions and strong OmniPath support. A second module involved $CD14^+$ and $CD16^+$ monocyte-to-MAIT HLA-DQB1–LAG3 interactions, characterised by high ligand–receptor expression, increased predicted ligand activity in *F508del* carriers, high receptor specificity and substantial ligand/receptor expression fractions, although with lower OmniPath evidence scores. Together, panel B supports a carrier-prioritised monocyte-derived communication pattern involving intra-myeloid TNF-family signalling and predicted monocyte–MAIT MHC-II/LAG3-associated communication. **(C)** Predicted ligand activity, ligand–target links and target-gene expression in $CD14^+$ monocyte receivers from healthy *F508del* carriers versus non-carriers. Ligand–target visualisation was performed for healthy *F508del* carriers with $CD14^+$ monocytes defined as the receiver population. This panel shows the predicted downstream target-gene layer underlying prioritised ligand activity towards $CD14^+$ monocytes. The ligand-activity heatmaps display scaled and raw predicted ligand activity for prioritised ligands in $CD14^+$ monocyte receivers. The ligand–target heatmap shows predicted regulatory links between prioritised ligands and target genes in $CD14^+$ monocytes, based on the NicheNet ligand–target prior model. The sample-level expression heatmap shows pseudobulk expression of predicted target genes in $CD14^+$ monocytes across individual healthy *F508del* carrier and non-carrier samples. In this receiver-cell analysis, TNF emerged as the dominant predicted active ligand in *F508del* carriers. TNF was linked to a coherent set of predicted target genes in $CD14^+$ monocytes, including inflammatory, NF-κB feedback and signalling-regulatory genes such as *NFKBIA*, *TNFAIP3*, *IRF1*, *SOCS3*, *DUSP2*, *TNFSF10* and related targets.

#### Supplementary Figure 11

##### A. Mean paired cytokine changes across adult CFTR modulation subgroups

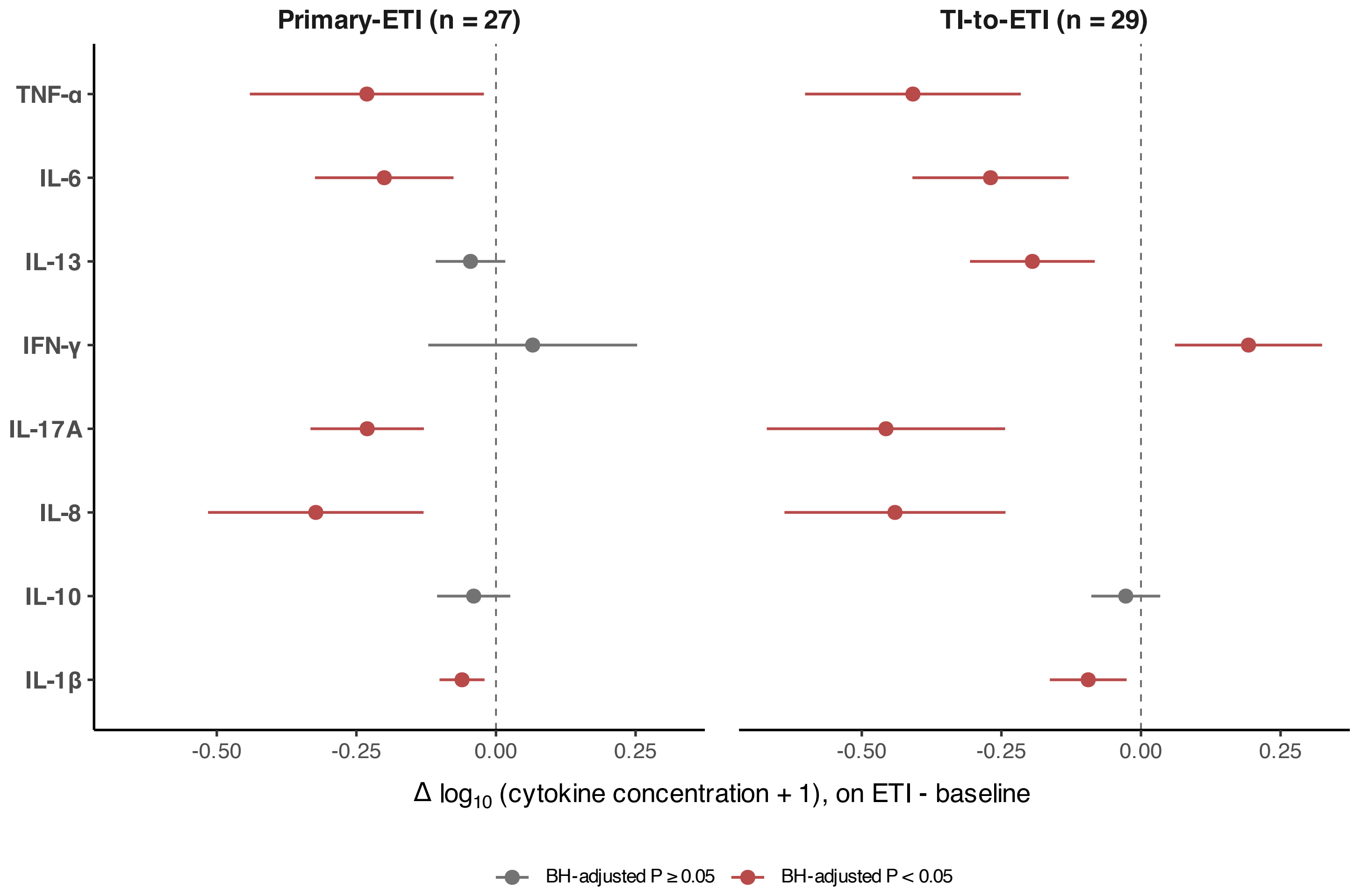

#### B. Three-timepoint cytokine trajectories in the TI-to-ETI cohort

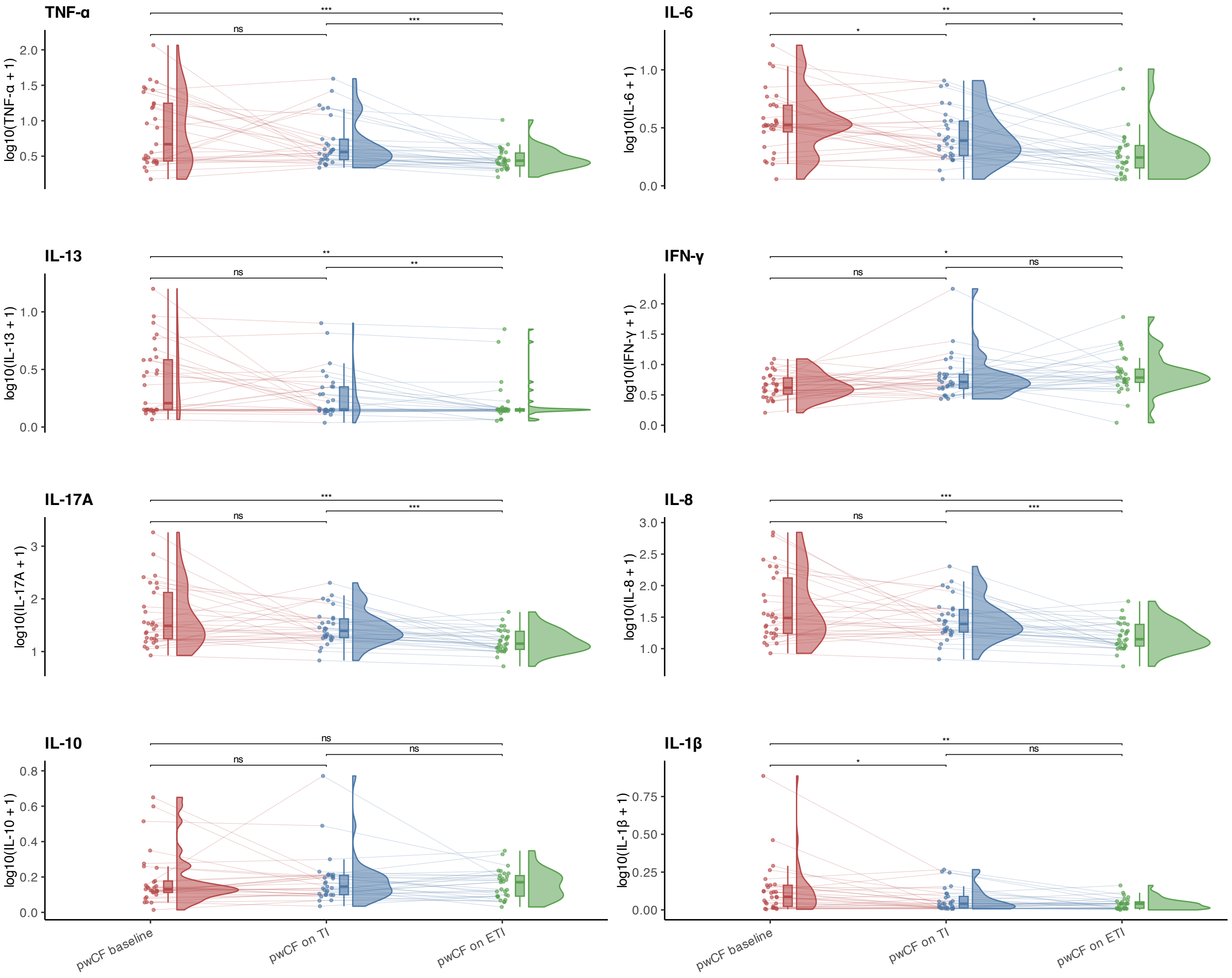

**C. Baseline systemic cytokine profiles in adult primary-ETI versus TI-to-ETI subgroups**

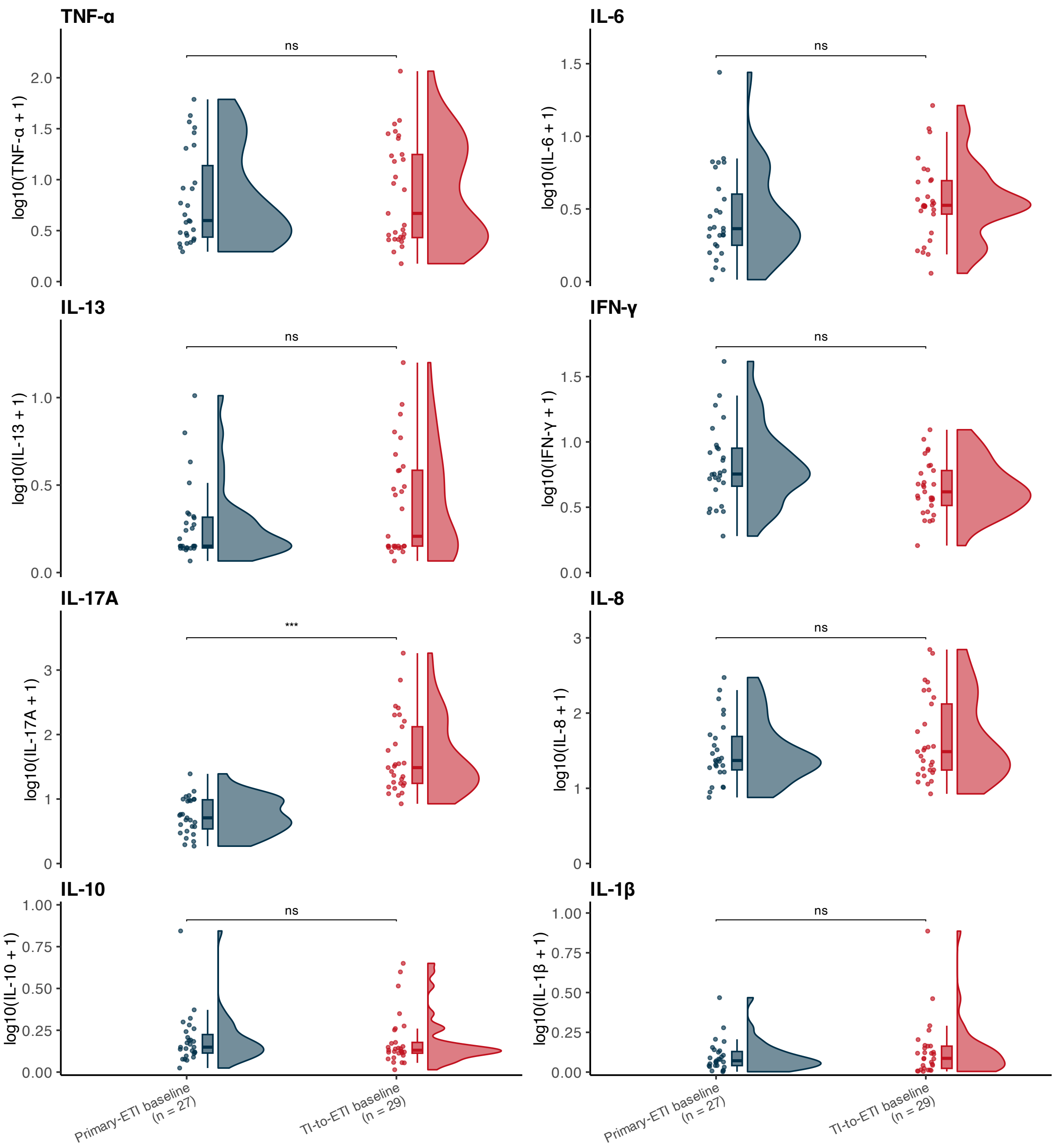


**Supplementary Figure 11. Longitudinal and baseline systemic cytokine profiling across adult CFTR modulation subgroups. (A)** Summary of paired cytokine changes following ETI initiation in adult pwCF. Forest plots display mean paired changes in log10-transformed serum cytokine concentrations between baseline and on-ETI samples, with 95% confidence intervals. Results are shown separately for the primary-ETI subgroup (baseline to ETI) and the TI-to-ETI subgroup (transition from tezacaftor/ivacaftor to ETI). Positive values indicate increased cytokine concentrations on ETI, whereas negative values indicate reductions following ETI initiation. Colour indicates whether the corresponding paired Wilcoxon signed-rank test remained significant after Benjamini–Hochberg correction across cytokines. **(B)** Longitudinal cytokine trajectories in the adult TI-to-ETI subgroup. Raincloud plots display log10-transformed serum cytokine concentrations at baseline, on tezacaftor/ivacaftor (TI), and on ETI in paired adult pwCF with three longitudinal timepoints (n = 29). Individual lines connect paired samples from the same participant. Statistical comparisons were performed using paired Wilcoxon signed-rank tests with Holm correction for multiple testing within each cytokine. Significance annotations indicate adjusted pairwise comparisons between timepoints. **(C)** Baseline systemic cytokine profiles in adult primary-ETI and TI-to-ETI subgroups. Raincloud plots show log10-transformed baseline serum cytokine concentrations in adult pwCF initiating ETI directly (primary-ETI, n = 27) or transitioning from prior TI therapy to ETI (TI-to-ETI, n = 29). Statistical comparisons were performed using two-sided Wilcoxon rank-sum tests with Benjamini–Hochberg correction across cytokines. Significance annotations indicate adjusted p-values. In all panels, cytokine concentrations were transformed using log10(x + 1).

### Supplementary Figure 12

#### A. Longitudinal cytokine trajectories in paediatric pwCF on ETI

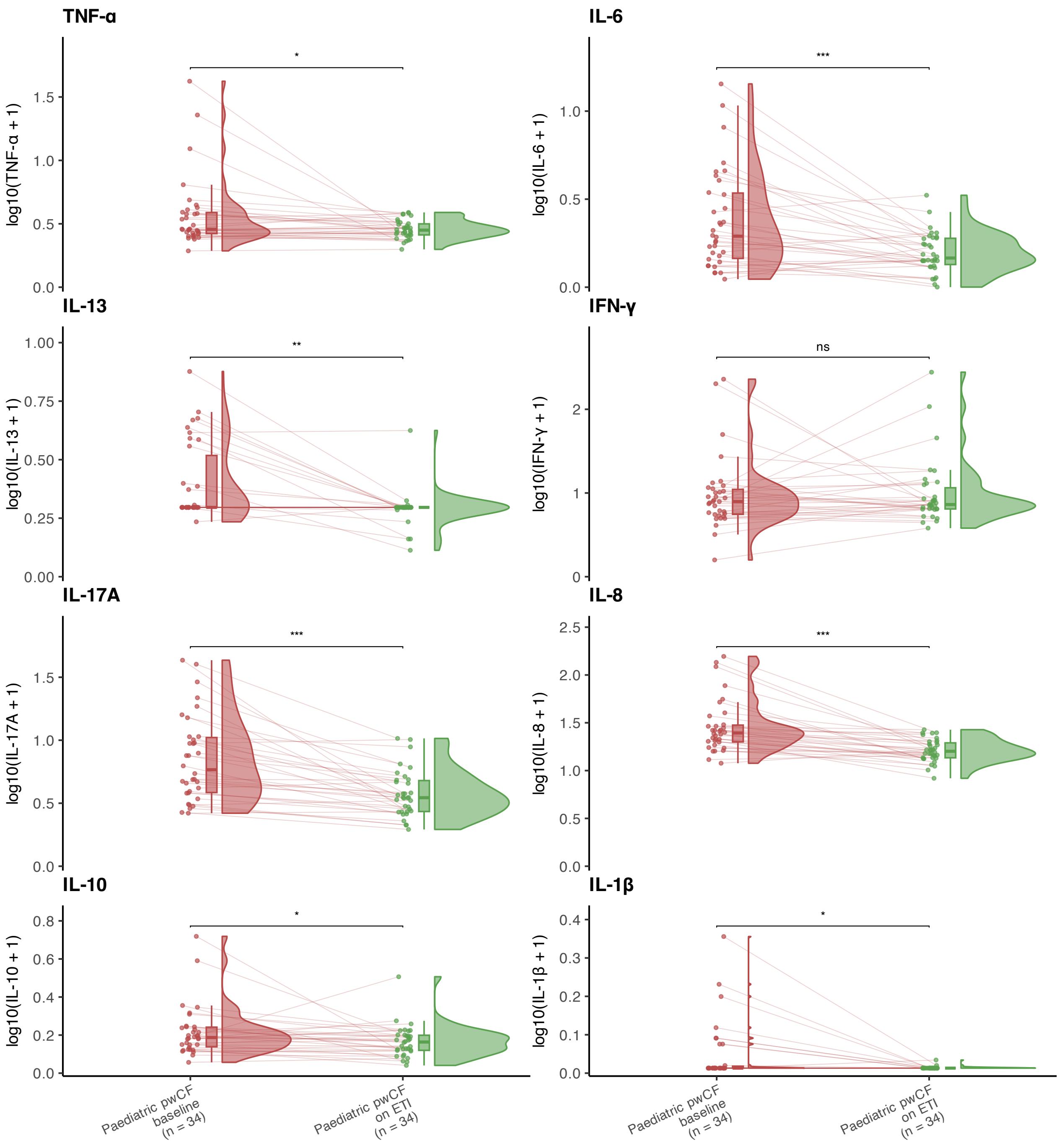

**B. Mean paired cytokine changes in paediatric pwCF on ETI**

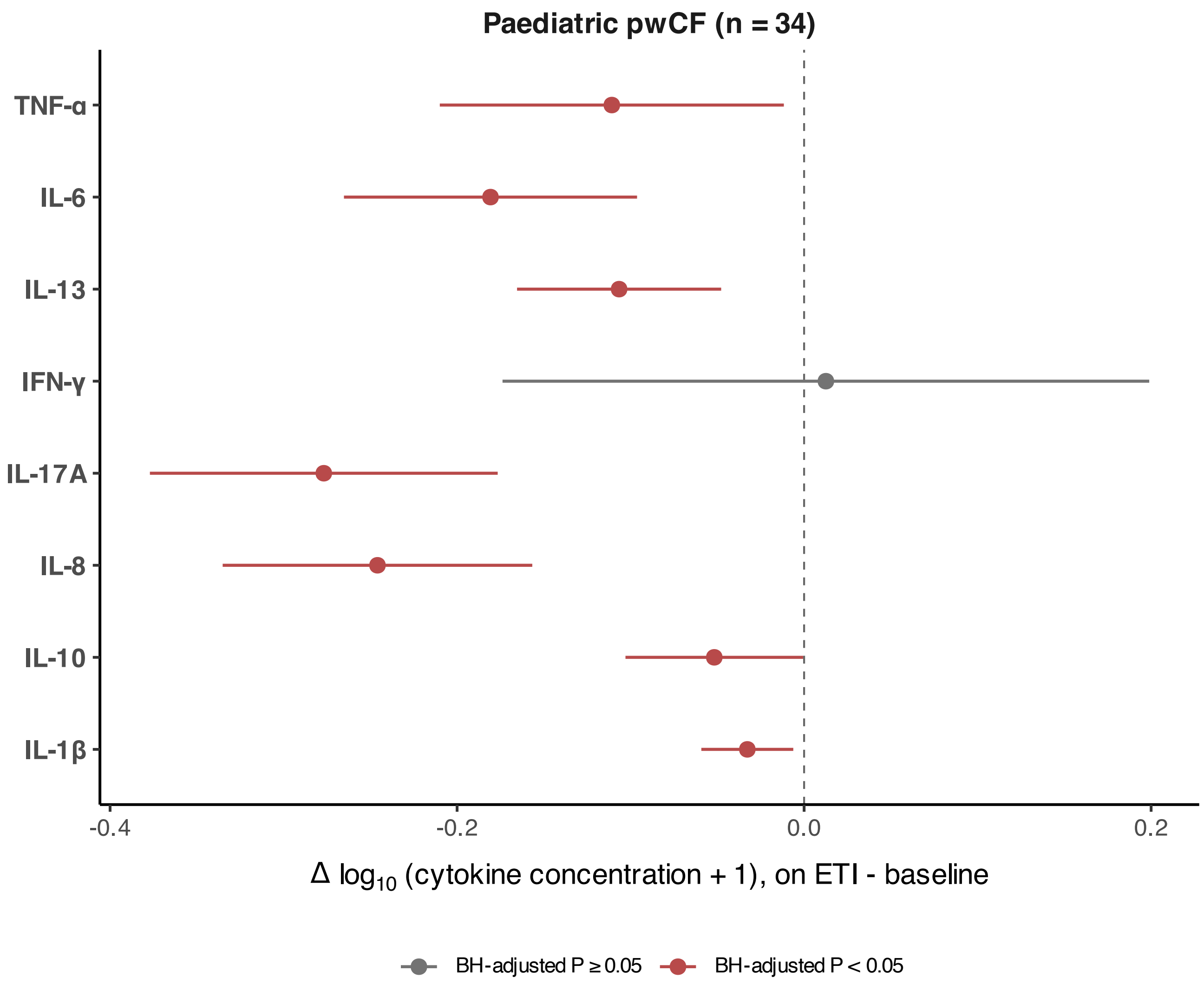

#### C. Comparison of longitudinal cytokine responses to ETI between adult and paediatric pwCF

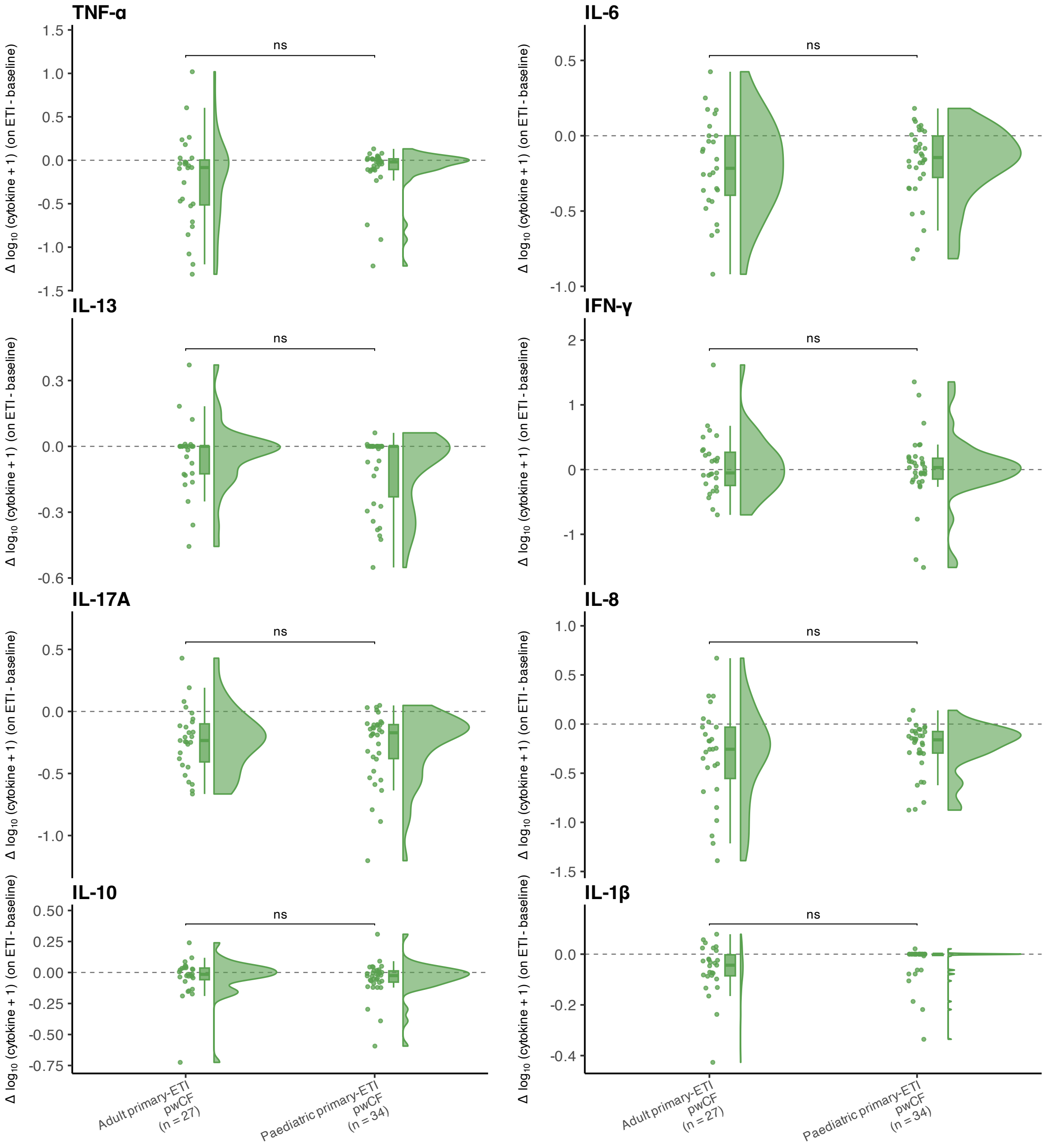


**Supplementary Figure 12. Paediatric systemic cytokine responses to ETI and comparison with adult ETI-associated cytokine remodelling. (A)** Longitudinal cytokine trajectories in paediatric pwCF on ETI. Raincloud plots display log10-transformed serum cytokine concentrations at baseline and on ETI in paired paediatric pwCF initiating ETI (n = 34). Individual lines connect paired samples from the same participant. Statistical comparisons were performed using paired Wilcoxon signed-rank tests with Benjamini–Hochberg correction across cytokines. Significance annotations indicate adjusted p-values. **(B)** Mean paired cytokine changes following ETI initiation in paediatric pwCF. Forest plots display mean paired changes in log10-transformed serum cytokine concentrations between baseline and on-ETI samples, with 95% confidence intervals. Positive values indicate increased cytokine concentrations on ETI, whereas negative values indicate reductions following ETI initiation. Colour indicates whether the corresponding paired Wilcoxon signed-rank test remained significant after Benjamini–Hochberg correction across cytokines. **(C)** Comparison of longitudinal cytokine responses to ETI between adult and paediatric pwCF. Raincloud plots display paired changes in log10-transformed serum cytokine concentrations between baseline and on-ETI samples in adult primary-ETI pwCF (n = 27) and paediatric pwCF (n = 34). Statistical comparisons were performed using two-sided Wilcoxon rank-sum tests with Benjamini–Hochberg correction across cytokines. Significance annotations indicate adjusted p-values. In all panels, cytokine concentrations were transformed using log10(x + 1).

#### Supplementary Figure 13

##### A. Baseline covariates show limited association with ETI-associated cytokine changes

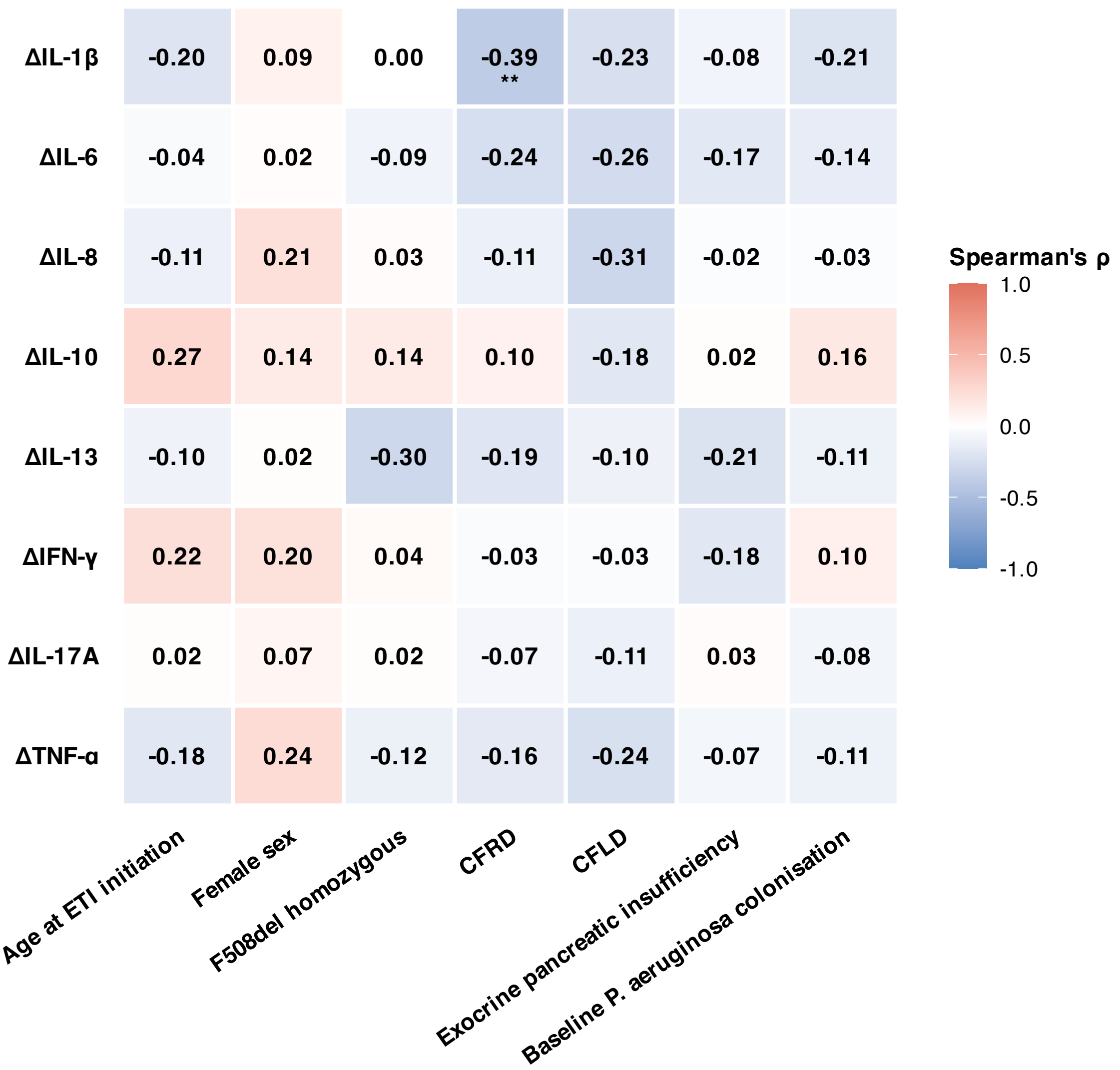

#### B. Higher baseline inflammation correlates with broader cytokine reductions in pwCF on ETI

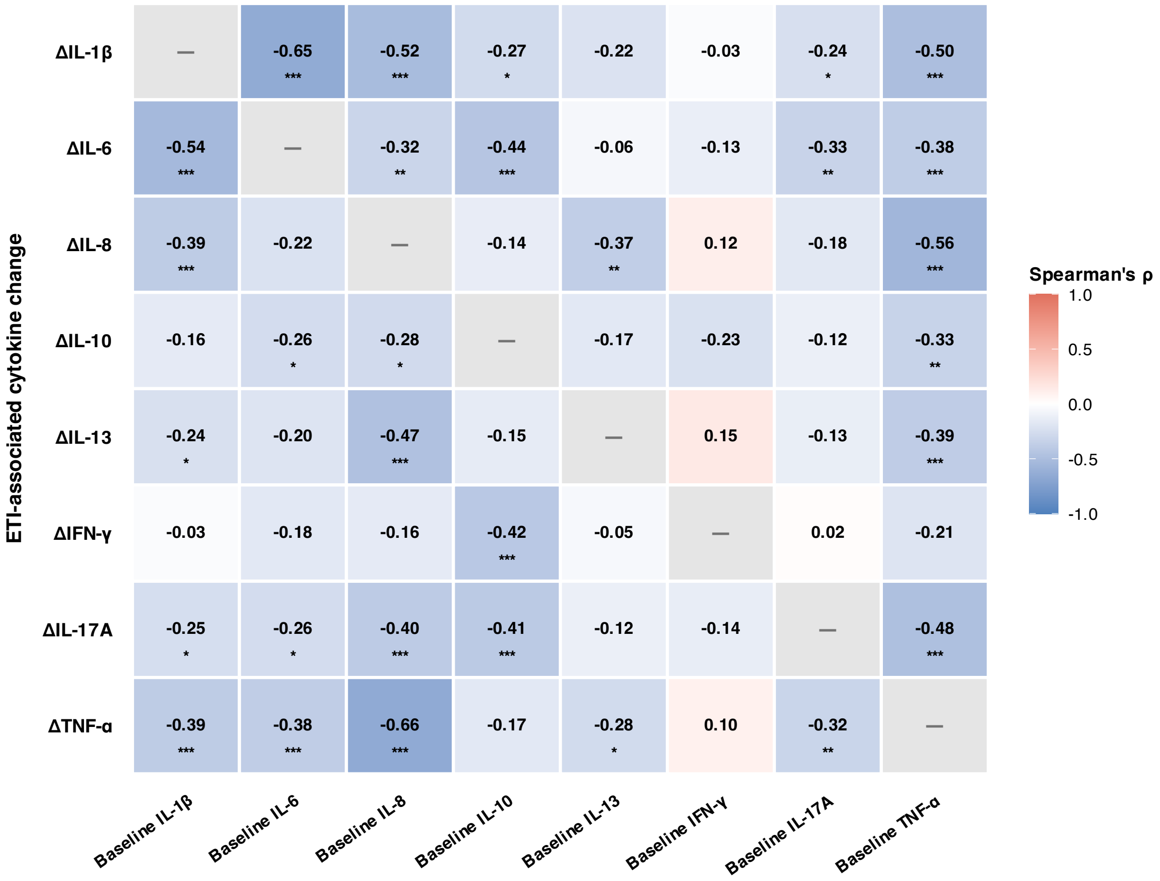

**C. IL-1β reduction is associated with BMI gain on ETI**

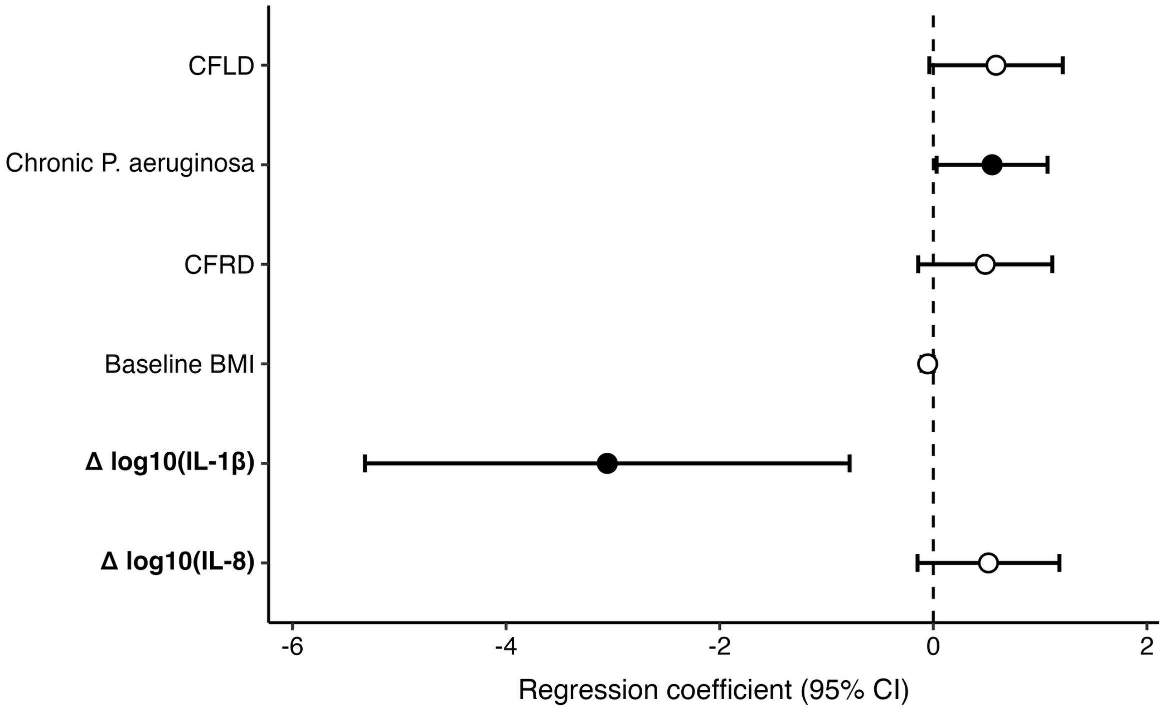


**Supplementary Figure 13. Baseline clinical and cytokine features associated with ETI response. (A)** Correlation heatmap showing associations between baseline clinical or demographic variables and longitudinal serum cytokine changes following ETI initiation. Cytokine changes were calculated as Δlog10(cytokine concentration + 1), defined as on-ETI minus baseline values. Correlations were assessed using Spearman's rank correlation. Colour intensity indicates Spearman's ρ, with negative values indicating that the presence or higher value of a baseline covariate was associated with a more negative cytokine change, corresponding to a greater cytokine reduction on ETI. P-values were adjusted across all displayed comparisons using the Benjamini–Hochberg method. **(B)** Correlation heatmap showing associations between baseline serum cytokine levels and subsequent ETI-associated cytokine changes. Baseline cytokine concentrations and cytokine changes were calculated from log10-transformed values. Same-cytokine baseline/change correlations were omitted because baseline values are mathematically included in the corresponding change score. Correlations were assessed using Spearman's rank correlation, with Benjamini–Hochberg correction across displayed cross-cytokine comparisons. Negative correlations indicate that higher baseline cytokine levels were associated with larger reductions in other cytokines on ETI. **(C)** Multivariable linear regression model of BMI change following ETI initiation. The forest plot displays regression coefficients with 95% confidence intervals for the retained model. Cytokine changes were defined as log10-transformed on-ETI values minus log10-transformed baseline values. Negative coefficients for cytokine-change terms indicate that larger cytokine reductions were associated with greater BMI gain. Filled points indicate predictors with $p < 0.05$ in the retained model. Abbreviations: BMI, body mass index; CFRD, cystic fibrosis-related diabetes; CFLD, cystic fibrosis-related liver disease; ETI, elexacaftor/tezacaftor/ivacaftor; IL, interleukin; *P. aeruginosa, Pseudomonas aeruginosa*.
